# The Entwined African and Asian Genetic Roots of the Medieval Peoples of the Swahili Coast

**DOI:** 10.1101/2022.07.10.499442

**Authors:** Esther S. Brielle, Jeffrey Fleisher, Stephanie Wynne-Jones, Nasreen Broomandkhoshbacht, Kim Callan, Elizabeth Curtis, Lora Iliev, Ann Marie Lawson, Jonas Oppenheimer, Lijun Qiu, Kristin Stewardson, J. Noah Workman, Fatma Zalzala, George Ayodo, Agness O. Gidna, Angela Kabiru, Amandus Kwekason, Audax Z.P. Mabulla, Fredrick K. Manthi, Emmanuel Ndiema, Christine Ogola, Elizabeth Sawchuk, Lihadh Al-Gazali, Bassam R. Ali, Salma Ben-Salem, Thierry Letellier, Denis Pierron, Chantal Radimilahy, Jean-Aimé Rakotoarisoa, Brendan Culleton, Kendra Sirak, Swapan Mallick, Nadin Rohland, Nick Patterson, Mohammed Ali Mwenje, Khalfan Bini Ahmed, Mohamed Mchulla Mohamed, Sloan Williams, Janet Monge, Sibel Kusimba, Mary E. Prendergast, David Reich, Chapurukha M. Kusimba

## Abstract

The peoples of the Swahili coast of eastern Africa established a literate urban culture by the second millennium CE. They traded across eastern Africa and the Indian Ocean and were among the first sub-Saharan practitioners of Islam. An open question has been the extent to which these early interactions between Africans and non-Africans were accompanied by genetic admixture. We report genome-wide ancient DNA from 80 individuals in five medieval and early modern (1300-1800 CE) coastal towns, as well as people from an inland town postdating 1650 CE. Over half of the ancestry of most coastal individuals came from African ancestors; these African ancestors were primarily female. A slightly smaller proportion of ancestry was from Asia. This Asian component was approximately eighty to ninety percent from Near Eastern males and ten to twenty percent from Indian females. Peoples of African and Asian origins began to mix by around 1000 CE, a time when archaeological evidence documents changes on the coast that are often interpreted as marking the large-scale adoption of Islam. Before roughly 1500 CE, the Near Eastern ancestry detected in the individuals was mainly Persian-related, consistent with the narrative of the Kilwa Chronicle, the oldest history told by the Swahili themselves. After this time, the sources of Near Eastern ancestry became increasingly Arabian, consistent with the archaeological and historical evidence of growing interactions between the Swahili coast and parts of southern Arabia. Subsequent interactions of Swahili coast peoples with other Asian and African groups further changed the ancestry of present-day peoples relative to the ancient individuals we sequenced, highlighting how Swahili genetic legacies can be more clearly understood with ancient DNA.

## Introduction

The Swahili culture of eastern Africa has been defined by a set of shared features: a common language of African origin (Kiswahili), a shared predominant religion (Islam), and a geographic distribution in coastal towns, many of which have long served as trading centers, linking peoples across the Indian Ocean to peoples of inland Africa. In the late medieval and early modern period, Swahili people lived over a vast coastal region spanning present-day northern Mozambique to present-day southern Somalia and including islands (Madagascar) and archipelagoes (Comoros, Kilwa, Mafia, Zanzibar, Lamu) (yellow outlines in **Figure 1**A) [1, 2].

**Figure 1:**
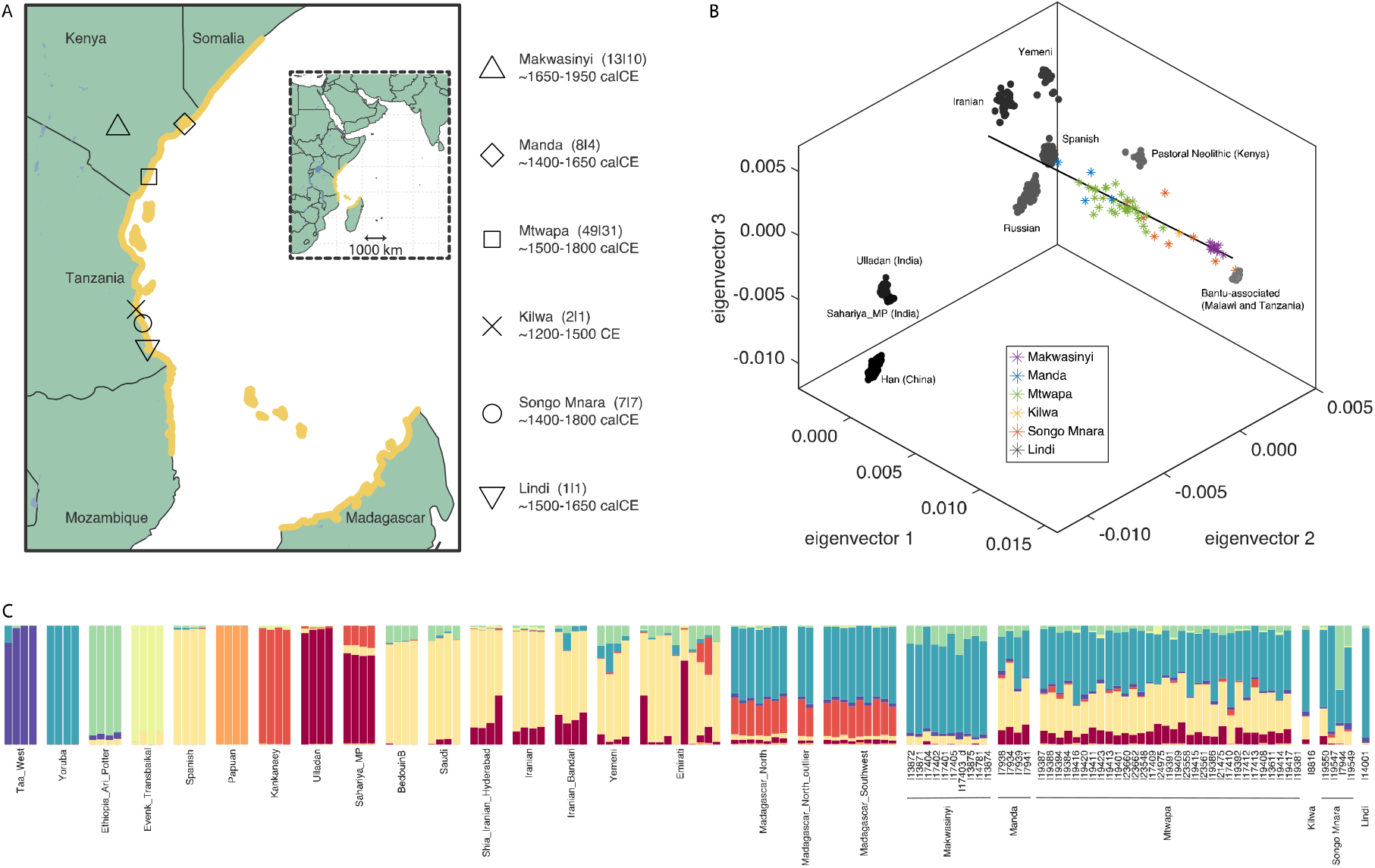
Dataset overview. A) The coastal areas associated with the Swahili culture are outlined in yellow. Sites included in this study are marked with black shapes and labeled on the zoom-in inset with the number of individuals, number of non-first-degree relatives with sufficient data to analyze extensively, and approximate date range. B) A three-dimensional principal component analysis shows present-day and ancient people studied in order to contextualize the ancestry of the Swahili individuals, labeled with asterisks. Eigenvector 1 correlates to variation maximized in sub-Saharan Africa. Together, eigenvectors 2 and 3 describe variation maximized in Eurasia. For 2D PCAs to aid visualization of the same data, see **Extended Data Figure 3. C**) Ancestry component assignment using ADMIXTURE with K=8 ancestral reference populations. Mtwapa, Manda, Kilwa, and some Songo Mnara individuals have evidence of shared ancestry with Near Eastern, South Asian, and African sources. Songo Mnara individuals appear to be heterogenous in their ancestry components and proportions, reflecting cosmopolitanism along the Swahili coast. The Lindi individual falls near people with western African ancestry including present-day Bantu speakers.

Narratives of Swahili identity have dominated research over the last 50 years, which has demonstrated the autochthonous development of coastal settlement, yet questions of genetic origins have remained unanswered. First millennium CE sites on the littoral, beginning in the 7th century CE, were part of a shared material culture and practice network across the eastern African region [3, 4] and were fully engaged in Indian Ocean trading networks [5]. The annual southeast monsoons from May to October allowed trading vessels to travel from India or the Arabian Peninsula to the eastern African coast, and the northeast monsoons from November to March enabled their return in the same year [5, 6]. Muslims were present on the coast from the 8^th^ century CE, although probably in a minority [7]. A transition can be identified in the archaeology of the coast during the 11th century, with the establishment of new settlements and the elaboration of older ones with coral-built mosques and tombs, all the while remaining deeply connected to sites throughout eastern Africa [8]. This archaeological transition has been discussed as reflecting the majority adoption of Islam [3, 9]; it can also be seen in the distinction of coastal ceramics and material traditions from hinterland assemblages [10].

In the 16th century, Portuguese naval and economic dominance in the Indian Ocean disrupted prior power relations [11, 12]. As Portuguese influence waned in the early 18^th^ century, the Sultanates of Oman and later Zanzibar ruled the eastern African coast [13]. In the 19^th^ century, growth of overseas trade, particularly the slave trade, led to large-scale movements of people from central regions of Africa to the coast and the settlement of people from the Yemenite region of Hadramawt [13, 14]. In the mid-19^th^ century, other European powers, especially the British, took administrative and military control, sent settlers, and precipitated significant population movements, including importing workers from South Asia. In addition, there was continued interaction with other eastern African groups, including pastoralist and farming groups along the coast and in the interior. Because present-day people, identified as Swahili, may differ in their genetic ancestry compared to populations of the 11^th^ to 18^th^ centuries, the origins of earlier peoples of the Swahili culture cannot be reconstructed without the aid of ancient DNA.

A set of common oral traditions recorded along the coast relate the founding of coastal towns to the arrival of a group known as the Shirazi, referring to a region in Persia [15–17]. This Shirazi tradition was first put into writing in the Kilwa Chronicle, transcribed in Portuguese in the 16th century [18]. The Swahili accounts of Shirazi roots were central to the narrative constructed by mid-20^th^ century colonialist archaeologists. They interpreted second-millennium coastal eastern African sites as the remains left by Persian and Arab settlers. Largely ignoring the evidence of a complex society created by peoples of African origin [10], colonial archaeologists focused on cultural traits showing connections with non-African peoples of the Indian Ocean world [19, 20].

Studies in the last decades have shown how Swahili elites have emphasized foreign origins as part of status claims and signals of cultural and religious affinities to support social stratification [21, 22]. Postcolonial research has challenged narratives of medieval Swahili peoples having roots outside Africa [7, 23]. Historical linguistics has revealed Kiswahili as a Bantu language, albeit with Asian loanwords [24, 25]. Archaeological material culture of the people who inhabited second-millennium villages and towns shows continuity with earlier settlements, including the persistence of crops, domesticated animals, ceramics, and craft styles [7, 26]. Thus, postcolonial research has shown that Swahili culture had autochthonous roots, coexisting with long-standing engagement with Indian Ocean trade networks.

The intercontinental connections that Swahili groups maintained as part of the Indian Ocean trading network probably meant that foreigners were consistently present along the coast. The extent to which these foreigners immigrated, intermarried, and otherwise intermingled with African residents has long been debated, thus raising questions about Swahili coastal groups’ ancestry [27].

This study presents the first genetic analysis of the medieval people of the Swahili coast of eastern Africa. We generated ancient DNA data from the skeletal remains of individuals found at five coastal or island towns: Manda, Mtwapa, Kilwa, Songo Mnara, and Lindi. These samples date to the mid-second millennium 1300-1800 CE but provide insight into genetic admixture events of the medieval period spanning from the 10^th^-11^th^ centuries CE onwards. We also generate DNA from the skeletal remains of individuals found at the site of Makwasinyi (∼1650-1950 CE), about 100 km inland from the southern Kenyan coast, which has been populated by people who were in cultural contact with coastal groups. We compare the newly reported data from the ancient individuals to that of present-day coastal Swahili speakers [28, 29] and to published data from diverse ancient and present-day eastern African [30, 31] and Eurasian groups [31–38].

### Data set

We used in-solution enrichment for a targeted set of about 1.2 million single nucleotide polymorphisms (SNPs; see Methods) to generate genome-wide ancient DNA data from skeletal remains of 80 individuals buried at six second-millennium CE sites in eastern Africa (black shapes in **Figure 1**A, analyzed individuals listed in **Extended Data Table 1**, see Methods and SI for an ethics statement and descriptions of archaeological and genetic permissions and sampling). We also radiocarbon dated enough individuals from each site to establish chronology, although from two sites (Kilwa and Lindi) the samples we analyzed had insufficient collagen preservation to obtain dates. Two coastal sites were from northern towns: Mtwapa (49 individuals spanning ∼1500-1800 cal CE (calibrated radiocarbon years)) and Manda Island (8 individuals dated to ∼1400-1650 cal CE). Three additional sites were from southern towns: Kilwa Island (2 individuals spanning ∼1300-1600 CE based on archaeological context), Songo Mnara Island (7 individuals spanning ∼1300-1800 cal CE), and Lindi (present-day Tanzania, 1 individual dated to ∼1500-1650 CE). The remains of individuals buried at Mtwapa, Manda, and Songo Mnara were mainly recovered from elite Muslim burials often located near mosques. We do not have enough context for the Kilwa and Lindi burials to know if they followed the same pattern. We also analyzed 13 individuals spanning ∼1650-1950 CE from an inland site, Makwasinyi (∼100 km inland from the coast of present-day Kenya). While these burials post-date coastal sites in our study, the Makwasinyi community traded with people in the coastal towns in past centuries and has remained relatively isolated. We thus hypothesized that the genetic ancestry of the Makwasinyi people might be a good proxy for inland African groups that may have been in contact with people from medieval cities on the northern Swahili coast in previous centuries [39, 40].

Of the 80 individuals for which we acquired genetic data, we removed 26 from our main analyses. Of these 26, 2 had contaminated sequences, 18 had too little data for high-resolution analyses, 5 were first- or possibly second-degree relatives of other individuals in the dataset with better quality data, and 1 was unreliable as it was a significant outlier with relatively low coverage (**Online Table 2 | newly_reported_individuals**). Of the remaining 54 individuals, 24 have direct Accelerator Mass Spectrometry (AMS) radiocarbon dates (**Extended Data Table 1** and **Online Table 3 | 14C_Results**).

### Genetic affinities of newly analyzed individuals

We performed principal component analysis (PCA, **Figure 1**B and **Extended Data Figure 3**), computing the eigenvector axes using 1286 present-day Eurasian and African individuals. Many of the newly reported ancient individuals projected onto this modern variation form a cline, with one end overlapping with ancient and present-day African groups and the other falling between present-day Persian and Indian groups. This cline suggests that the individuals might be mixed in different proportions of two source populations, where each can have multiple ancestry components. However, some coastal individuals, particularly from Songo Mnara and Lindi, do not fall along this cline. Individuals who deviate from the cline cannot be described by this same two-way mixture model and must harbor alternative ancestral contributions. Similar patterns of ancestry composition are evident with unsupervised clustering using ADMIXTURE (**Figure 1**C, **Extended Data Figure 1**), which suggests sub-Saharan African-associated components, Near Eastern-associated components, and East Asian or Indian-associated components.

**Figure 3:**
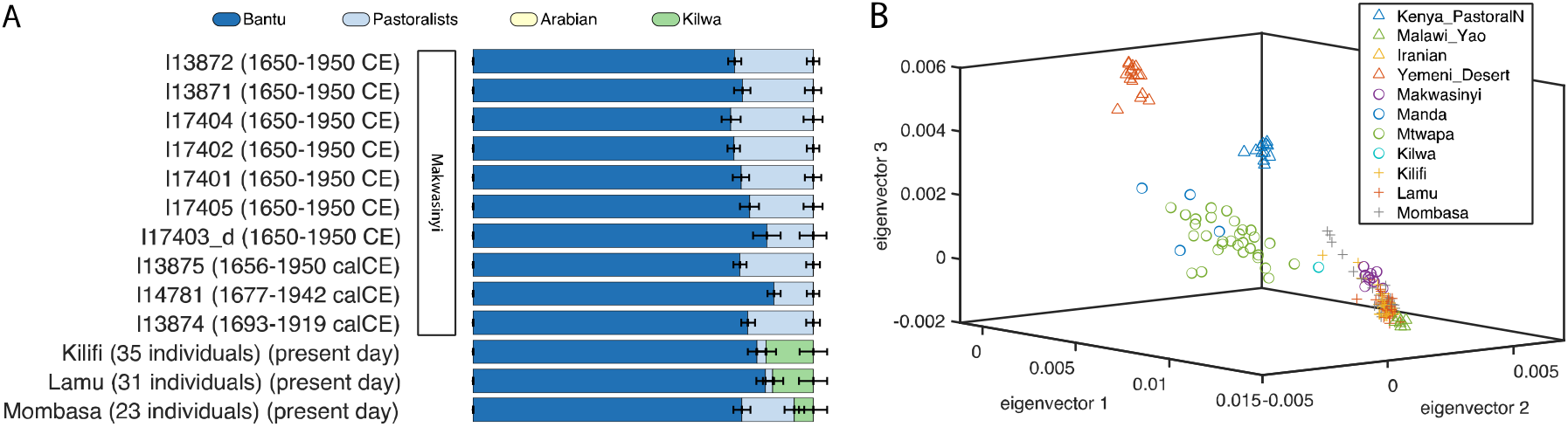
Relation of medieval Swahili people to modern and ancient eastern Africans. A) Graphical representation of individual and population level qpAdm models for Makwasinyi individuals (this study) and present-day people from Kilifi (n=35), Lamu (n=31), and Mombasa (n=23) who identify as Swahili [28]. The green ancestry component represents the modeled contribution from medieval Swahili coast people, as represented by the individuals buried in Kilwa. B) PCA with the same present-day people from Kilifi, Lamu, and Mombasa, as well as present-day people from Malawi, Iran, and Yemen (triangles) [31, 32, 37, 53], and ancient individuals from eastern Africa published in this study (open circles) and previously (triangles) [30, 31] projected onto the first three eigenvectors.

### Three-way mixture of Sub-Saharan African, Persian, and Indian ancestry

Using *qpAdm* [34], we identified a common three-source model that fits the data for most Swahili individuals (see **Supplementary qpAdm modeling** for a description of formal modeling). We used a local ancient African source and present-day Iranian and Indian groups as proxies for the three ancestry components (**Figure 2**A and **Extended Data Table 2**). Any model that does not include all three ancestries can be rejected with high statistical confidence. This model fits the pools of Mtwapa (P value for fit 0.23) and Manda (P=0.28) individuals, the Kilwa individual with good quality data (I8816) (P=0.27), and at least one Songo Mnara individual (I19550) (P=0.38). In individuals with relatively high proportions of African ancestry, such as the Kilwa individual, other contributions are small, and the Indian proportion can fall below the threshold of definitive detection.

**Figure 2:**
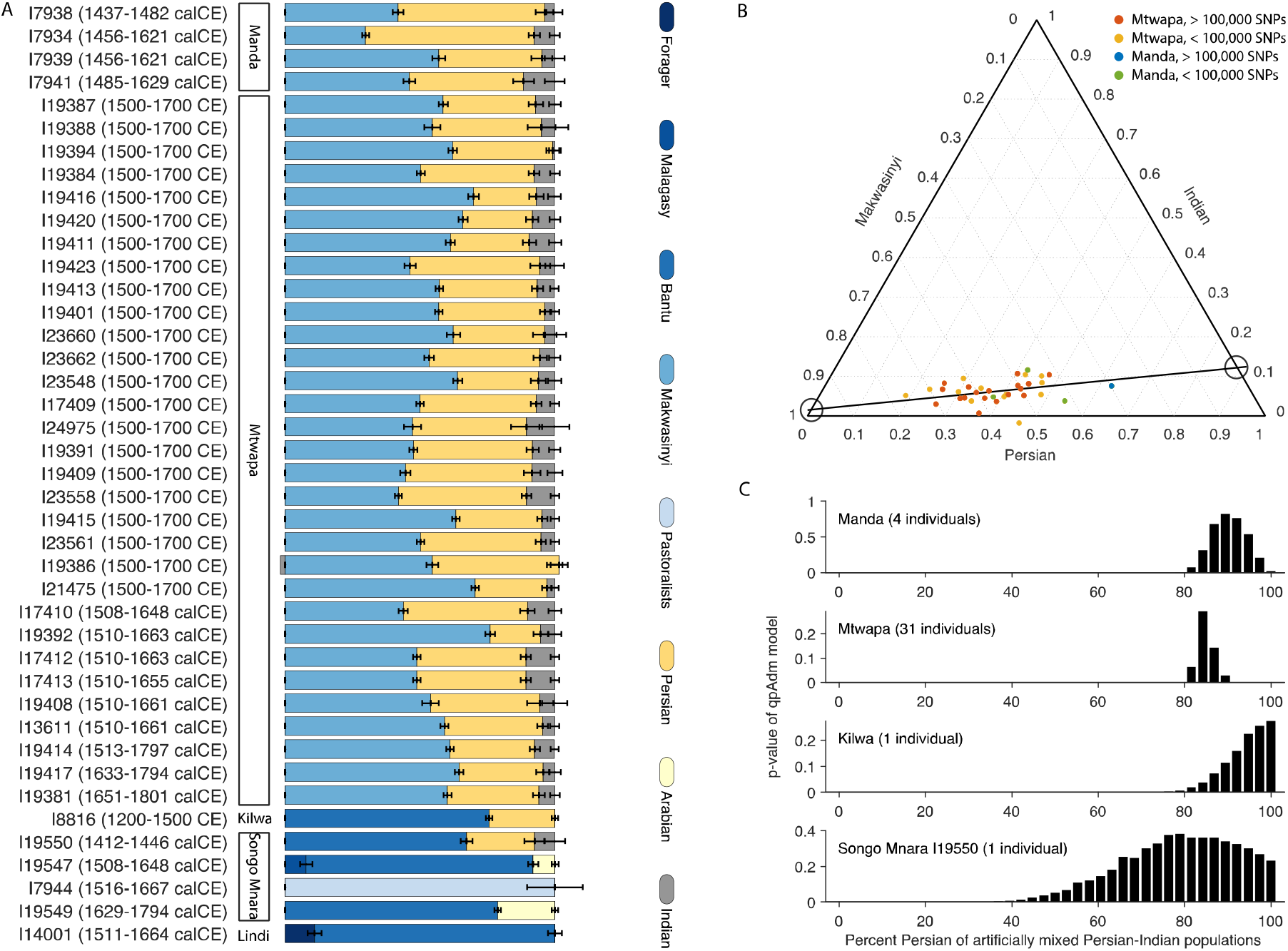
Individual mixture proportions. A) Inferences from qpAdm (bars show one standard error, see **Extended Data Table 2** and **Supplementary qpAdm modeling** for model details including source populations, proportions of admixture, and P-values for fit). Blue represents African components of ancestry: the most common are Bantu-associated (medium blue, common at southern sites), and Makwasinyi associated (medium-light blue, common at northern sites), which itself is ∼80% Bantu-related and ∼20% pastoralist-related. The African pastoralist-related ancestry in this diagram encompasses both Neolithic and Iron Age ancestry components from Kenya and Tanzania. Yellow represents Near Eastern ancestry: Persian (light yellow) or Arabian associated (dark yellow). Gray represents Indian ancestry. B) Ternary plot of Makwasinyi, Persian, and Indian components in Mtwapa (red points - high coverage, yellow points - low coverage) and Manda individuals (blue points - high coverage, green points - low coverage). Higher coverage individuals (>100,000 SNPs overlapping positions on the Human Origins array) are used to fit a linear regression (dashed line), which intersects at nearly 100% Makwasinyi and nearly 0% Persian and Indian ancestries, consistent with a Makwasinyi-related population with little or no recent Near Eastern ancestry mixing with an already mixed Persian-Indian population to form the ancestry of northern Swahili coast people. C) Bar graph showing P-values for a two-way qpAdm model with a local African source (using Makwasinyi as the source for Mtwapa and Manda individuals, and the Lindi individual as the source for the Kilwa individual and the Songo Mnara individual) and a mixed Persian-Indian source. The x-axis specifies the proportion of Near Eastern ancestry in this mixed source.

The African source of ancestry needed to make the models fit differed between the northern individuals in present-day Kenya (Mtwapa and Manda) and the southern individuals in present-day Tanzania (Kilwa and Songo Mnara). Among individuals from the Mtwapa and Manda sites, the best-fitting proxy source for the sub-Saharan African-associated ancestry is the group from the inland Makwasinyi site (**Extended Data Table 2**). Makwasinyi individuals themselves are well-modeled as mixtures of about 80% ancestry related to western African groups (an ancestry common today in speakers of Bantu languages and prevalent in eastern Africa; henceforth called ‘Bantu-associated’) and about a quarter ancestry from people related to ancient eastern African Pastoral Neolithic individuals [30] (**Figure 3**A and **Extended Data Table 3**, see **Supplementary qpAdm modeling** for a description of formal modeling). Among individuals from Kilwa and Songo Mnara, the best-fitting African proxy source is Bantu-associated without a Pastoral Neolithic contribution. We use the individual from this study buried at ∼1600 CE in Lindi as a proxy Bantu-associated source for the Kilwa individual and for the Songo Mnara I19550 individual (**Extended Data Table 2**).

The ancestry proportions for individuals from Manda and Mtwapa form a cline consistent with a mixture of just two source populations (**Figure 2**B). Using linear regression, we extrapolated the ancestry of these two sources and inferred that one source population was consistent with a 100% African origin (one standard error, **Supplementary qpAdm modeling**). The same analysis concluded that the other source had Indian and Persian ancestry. This data are consistent with sub-Saharan Africans mixing with a group that already had a mix of Persian and Indian ancestry components. The finding of multiple mixture events would be expected if people of mixed Persian and Indian ancestry had children with people from different local African populations at different locations along the coast.

When we separately analyze the Mtwapa, Manda, Kilwa, and Songo Mnara individuals, we infer possibly consistent proportions of Indian and Persian ancestry (the ranges are close and plausibly within normal variation, **Figure 2**C). Some variation in the ratio of Persian-to-Indian ancestry among the early immigrants is expected, making it difficult to distinguish between scenarios of one or many streams of Persian-Indian ancestral populations. However, there was high uncertainty for individual estimates of Indian proportions, and in some individuals (e.g., the one buried at Kilwa), it may in fact be absent (**Supplementary qpAdm modeling**). Finally, our analysis cannot determine if the Persian-Indian mixture occurred before these people arrived in eastern Africa or after.

### Male ancestors mainly from Persia; female ancestors mainly from Africa and India

We tested whether male and female ancestors (based on genetically determined sex) contributed equal proportions of the three ancestry components in the ancient individuals sampled at Mtwapa, Manda, and Kilwa (**Table 1**). We could not carry out the same analysis for Songo Mnara because there were no individuals who fit the three-way model and had data of sufficient quality to permit this analysis.

**Table 1:** Inferences about mixture for three Swahili coast groups.

|  | **Mtwapa (31 individuals)** |  |  | **Manda (4 individuals)** |  |  | **Kilwa (1 individual)** |  |
| --- | --- | --- | --- | --- | --- | --- | --- | --- |
|  | Africa | Persia | India | Africa | Persia | India | Africa | Persia |
| mtDNA haplogroups (N) | 30 | -- | 1 | 3 | -- | -- | 1 | -- |
| Y chrom. haplogroups (N) | 4 | 16 | -- | -- | 3 | -- | -- | 1 |
| Ancestry sources autosomes | 56-58% | 34-39% | 4-9% | 30-34% | 55-65% | 3-13% | 72-76% | 24-28% |
| Ancestry sources chrom. X | 65-75% | 0-24% | 7-39% | 49-70% | 0-74% | 0-75% | 83-100% | 0-17% |
| Z score autosomes vs. X for Makwasinyi | -4.7 | 3.4 | -2.0 | -5.0 | 1.4 | -0.5 | -3.3 | 3.3 |
| Prop. of this ancestry female | 76-90% | 0% | 100% | 100% | 0% | 23-100% | 80-98% | 0% |
| Prop. of this ancestry male | 10-24% | 100% | 0% | 0% | 100% | 0-77% | 2-20% | 100% |
| Prop. Persian (of total Asian) | 84-87% |  |  | 84-97% |  |  | 89-100% |  |
| Estimated admixture dates | 938-1242 CE |  |  | 629-1412 CE |  |  | 614-1336 CE |  |

*Notes: Ancestry estimates on chromosomes 1-22 and chromosome X are compared to infer proportions from female or male ancestors (Methods). Ranges for rows 3-4 are 95% confidence intervals based on estimated $\pm 1.96$ standard errors, truncated if extending outside 0-100%.*

We first analyzed mitochondrial haplogroups transmitted entirely by females, restricting to non-first-degree relatives. All but one individual carries an L* haplogroup, which today is almost wholly restricted to sub-Saharan Africans [41–43]. The one individual with a non-L haplogroup is from Mtwapa and carries haplogroup M30d1, which is largely restricted to South Asia [42]. The mtDNA haplogroup distribution of sampled coastal individuals is consistent with female ancestry deriving almost entirely from African or Indian sources, with a higher proportion of African female ancestors. We also expanded our mtDNA analysis to include ancient coastal and island individuals with low coverage or who did not pass quality control on the autosomes but did have sufficient mtDNA coverage. With this larger group of individuals, all but two have L* haplogroup indicative of sub-Saharan Africa. In addition to the M30d1 haplogroup, common in South Asia, we identified a male individual with haplogroup R0+16189, which is common in Saudi Arabia and the Horn of Africa [44].

We also examined male-transmitted Y chromosome sequences, which tell a different story than female-transmitted mitochondrial DNA. Two of three non-first-degree related males from Manda belong to haplogroup J2, while the third belongs to G2; both haplogroups are characteristic of Near Eastern ancestry (plausibly Persian) and are largely absent in sub-Saharan Africa [45–47]. The Kilwa individual also carries a J2 haplogroup. Fourteen of the twenty male Mtwapa individuals belong to J-family haplogroups, and two belong to R1a haplogroups, all of which are typically non-African lineages. The four remaining individuals belong to E1 family haplogroups, characteristic of sub-Saharan Africa. The Y chromosome haplogroup distribution is consistent with the male ancestry of sampled individuals from Mtwapa deriving mostly, but not entirely, from the Near East. The Y chromosome haplogroup distribution in Mtwapa differs qualitatively from that in Manda and Kilwa in that it includes more J1’s than J2’s, but is still consistent with the male ancestry of sampled individuals from Mtwapa deriving mostly, but not entirely, from the Near East.

Finally, we compared ancestry estimates from chromosomes 1-22 (the autosomes), which equally reflect the contributions of females and males, to those from chromosome X, which occurs in two copies in females compared to one in males, and thus predominantly reflects the contributions of female ancestors. African ancestry estimates from the autosomes are significantly lower in the individuals from Mtwapa, Manda, and Kilwa than estimates made from chromosome X: the 95% confidence intervals are non-overlapping, and the absolute value of the Z-score for a difference between the two estimates is greater than 2. This implies that the African ancestry in medieval Swahili coast populations primarily came from females, consistent with the mitochondrial DNA and Y chromosome evidence (**Table 1**). Similarly, the Persian ancestry in all three groups primarily came from males (**Table 1**). Under the simplifying assumption that the mixture occurred in a single event over just a few generations, rather than over a period that spanned many generations, we estimate the proportion of African ancestry contributed by females: 100% in sampled individuals at Manda (95% confidence interval entirely above this value), 76-90% at Mtwapa, and 80-98% at Kilwa (Methods, **Table 1**). Using the same approach, we estimate the proportion of Persian ancestry contributed by males to be 100% in sampled individuals in all three locations.

### All three ancestry components began mixing by about 1000 CE

We used the software DATES [48] to estimate when mixture occurred. We detect strong signals of linkage disequilibrium driven by a mixture between ancestral populations of sub-Saharan African and Asian ancestry, with 95% confidence intervals for the inferred dates of 938-1242 CE for Mtwapa, 629-1412 CE for Manda, and 614-1336 CE for Kilwa (**Table 1** and **Extended Data Figure 2**). The uncertainty intervals of the three groups overlap from 938 to 1242 CE. The Songo Mnara individual that fits the three-way model does not have sufficient data to estimate a date of mixture, but since the other three date ranges overlap, it is reasonable to hypothesize that they derive from mixture in the same period. Our analysis suggests that already admixed males with both Indian and Persian ancestry were present along the coast by ∼1000 CE and mixed further with primarily female sub-Saharan Africans.

### In recent centuries, the coast was impacted by Arabian (plausibly Omani) migration

For the earliest individuals in our study with a Near Eastern ancestry component (individuals from Manda, Kilwa, and the I19550 individual from Songo Mnara), there is no evidence of Arabian-associated ancestry that is not well proxied by Persian sources. However, later individuals can be modeled as having Arabian-related ancestry, such as those buried at Mtwapa with radiocarbon dates in the 16^th^ to 18^th^ centuries. We were not able to determine the exact source of the Arabian-associated ancestry in the Mtwapa group. However, we know that it is somewhere on the genetic gradient between Arabians and Persians. We are also not able to determine if there is added Persian-Arabian ancestry or only added Arabian ancestry since the significant Persian admixture from ∼1000 CE would mask any smaller or partial subsequent additions of Persian ancestry. In one of our analyses, this added Persian-Arabian ancestry source for Mtwapa is well proxied by groups bordering the Strait of Hormuz, which separates the Arabian Peninsula from present-day Iran. The Strait of Hormuz and the Swahili coast were under Omani control simultaneously (beginning at the end of the 17^th^ century). While Mtwapa individuals have Arabian and Persian ancestries, we do not detect Arabian-associated ancestry when we apply a similar analysis to the Manda individuals who lived earlier than the Mtwapa individuals. Since Manda individuals lived sometime between the 15^th^ and 17^th^ centuries, they predate Omani political dominance along the Swahili coast.

Further evidence for Arabian immigration affecting parts of the Swahili coast in the later period comes from two individuals from Songo Mnara (I19547 and I19549). Both date to after 1500 CE and can be modeled with Arabian-related ancestry in *qpAdm* (Figure 2A).

### Relationship between modern and medieval peoples of the eastern African coast

We analyzed genome-wide data from 89 people previously reported by Brucato et al. After removing two outliers (according to the ADMIXTURE graph in Extended Data Figure 1) from analysis, we found that these present-day people who live in coastal areas of Kilifi, Lamu, and Mombasa in Kenya and who identify as Swahili [28] typically have a small genetic inheritance from ancient people who lived in nearby coastal areas (proxied by the individual buried at Kilwa) (**Figure 3**A and **Extended Data Table 3**, **Supplementary qpAdm modeling**). We infer point estimates in the three coastal populations of 79-86% Bantu-associated ancestry proxied by the Lindi individual (all standard errors about 3%); 2-15% Pastoral Iron Age-associated ancestry [30] (all standard errors about 3%); and 6-14% ancestry from medieval Swahili-associated ancestry (all standard errors about 4%) (**Figure 3**A).

We also examined mitochondrial and Y chromosome haplotype determinations published in the same study [28]. A total of 231 of 237 individuals with mitochondrial DNA haplogroup determinations (97%) were from the L and M1 lineages, which are widespread in Africa, consistent with almost all female-lineage ancestry being from Africa [49–52]. Of 107 Y chromosome haplogroups, 102 (95%) are typically African, which is very different from medieval coastal individuals from whom almost all Y chromosome haplogroups are Near Eastern associated (Table 1).

The Brucato et al. Y chromosome haplogroup frequencies are not characteristic, however, of all modern coastal populations that identify as Swahili. Raaum et al. carried out an independent study of these frequencies, investigating mitochondrial and Y chromosomes of 179 present-day Swahili men from thirteen locations along the Kenyan coast (8 in the Lamu archipelago, 3 near Mtwapa and 2 near the Tanzanian border) [29]. The Raaum et al. study found that the African-associated Y haplogroup frequency was 45%, which was much less than the 95% in Brucato et al. albeit also substantially larger than the 17% in the medieval individuals (see Table 2). Thus, although Brucato and Raaum are sampling Swahili groups from roughly the same areas in Kenya, their findings are qualitatively different, implying population structure in modern day Swahili. The Raaum et al. study may have been biased toward upper-class Swahili society because they recruited participants through snowball sampling relying on initial contacts known to field assistants, while the Brucato et al. sampling strategy relied on public meetings. The medieval Swahili sample may more closely resemble the sample in the Raaum et al. study because our excavations focused on graves located near mosques, likely biasing toward more elite residents. Still, both studies were in agreement in showing a large increase in the proportion of African haplogroups relative to the medieval sample.

**Table 2:** Comparison of medieval Swahili haplogroup distribution with those from two studies of present-day Swahili individuals.

| **Likely Y chromosome haplogroup origin** | **Present-day Raaum et al [29]** | **Present-day Brucato et al [28]** | **Medieval Present study** |
| --- | --- | --- | --- |
| Africa | 45% | 95% | 17% |
| Near East | 35.5% | 3% | 83% |
| Other | 19.5% | 2% | 0% |
| **Likely mtDNA HG origin** | **Raaum et al [29]** | **Brucato et al [28]** | **Present study** |
| Africa | 94% | 97% | 97% |
| South or East Asia | 4% | 2% | 3% |
| Other | 2% | 1% | 0% |

### Ancient non-coastal groups are genetically similar to present-day coastal groups

The predominant genetic profile of Bantu-associated and pastoralist-associated ancestry found in present-day Kenyan coastal populations is similar to that in the Makwasinyi individuals located 100 km inland from the Kenyan coast (**Figure 3**A). The Makwasinyi individuals date to the last three centuries. They are from a relatively isolated population located deep into the Tsavo region and nearly 50 kilometers from their nearest neighbors. The Makwasinyi group fits as an African ancestry proxy source for *qpAdm* modeling of the Mtwapa and Manda individuals. Unlike the Mtwapa and Manda individuals, Makwasinyi individuals have no evidence of any recent Asian ancestry according to *qpAdm*. Instead, they are well modeled as 21.3±1.2% Pastoral Neolithic-associated ancestry and 78.7±1.2% Bantu-associated ancestry (**Figure 3**A). Herders were present in eastern Africa from at least 3000 BCE but are not documented beyond northernmost inland Kenya until two millennia later [30]. They persisted after people who practiced farming spread from west to east Africa in the first millennium CE, often associated with the Bantu expansion [26]. The sex-bias determination in the Bantu-Pastoral Neolithic mixture in Makwasinyi varies depending on the individual used as the Bantu-associated proxy source. When we use the 600BP individual buried at Makangale Cave on Pemba Island, we infer that Bantu-related ancestors were 37-50% female and 50-63% male, whereas KPN-related ancestors were 48-100% female and 0-52% male (Methods, **Supplementary qpAdm modeling**). However, when we calculate male and female ancestry proportions using the individual buried at Lindi, we infer that Bantu-related ancestors were 48-74% female and 26-52% male, whereas KPN-related ancestors were 0-60% female and 40-100% male. The variation could reflect the inherently poor statistical resolution of the X chromosome estimates and different degrees of sex-biased mixture in the source populations themselves. The average date for the Bantu-Pastoral Neolithic mixture is ∼300-1200 CE (**Extended Data Figure 2**), with most of this date range consistent with the archaeological evidence for the impact of the Bantu expansion on this region.

### Outliers past and present highlight the cosmopolitanism of the Swahili coast

While the main patterns of variation in the dataset are driven by admixture involving people with African, Persian, and Indian ancestries; there are some individuals at Songo Mnara and Lindi that do not have any evidence of recent Asian ancestry (I14001 dating to 1511-1664 calCE and I7944 dating to 1516-1667 calCE) (**Figure 3**A and **Extended Data Table 3**, **Supplementary qpAdm modeling**). We detect possible Arabian or Omani-associated ancestry in the Mtwapa group (**Supplementary qpAdm modeling**) and perhaps in two third-degree relatives from Songo Mnara (I19547 dating to 1508-1648 calCE and I19549 dating to 1629-1794 calCE) (**Figure 3**A and **Extended Data Table 3**, **Supplementary qpAdm modeling**). In addition, we find the possibility of Malagasy-associated ancestry in Songo Mnara (I19547 dating to 1508-1648 calCE) (**Figure 3**A and **Extended Data Table 3**, **Supplementary qpAdm modeling**). These coastal individuals who differ genetically compared to the others from similar times or geographic locations testify to the cosmopolitan nature of this region and its involvement within the inter-continental Indian ocean trading network.

## Discussion

A key finding of this study (**Figure 4** and **Table 1**) is genetic evidence of admixture at roughly 1000 CE between people of African and people of Persian ancestry. This admixture is consistent with one strand of the history recorded by the Swahili themselves, the Kilwa Chronicle, which describes the arrival of seven Shirazi (Persian) princes on the Swahili coast. At Kilwa, coin evidence has dated a ruler linked to that Shirazi dynasty, Ali bin al-Hasan, to the mid-11^th^ century [54]. Whether or not this history has a basis in an actual voyage, ancient DNA provides direct evidence for Persian-associated ancestry deriving overwhelmingly from males and arriving on the eastern African coast by about 1000 CE. This timing corresponds with archaeological evidence for a substantial cultural transformation along the coast, including the widespread adoption of Islam [3, 8, 9]. It is difficult to distinguish between a scenario in which a mixture event occurred within a few generations and one drawn out over dozens of generations. While we can be confident that mixture had begun by 1000 CE, in theory, it could have started earlier and continued later. We emphasize that the people of African ancestry made the largest genetic contribution to the mixed population.

**Figure 4:**
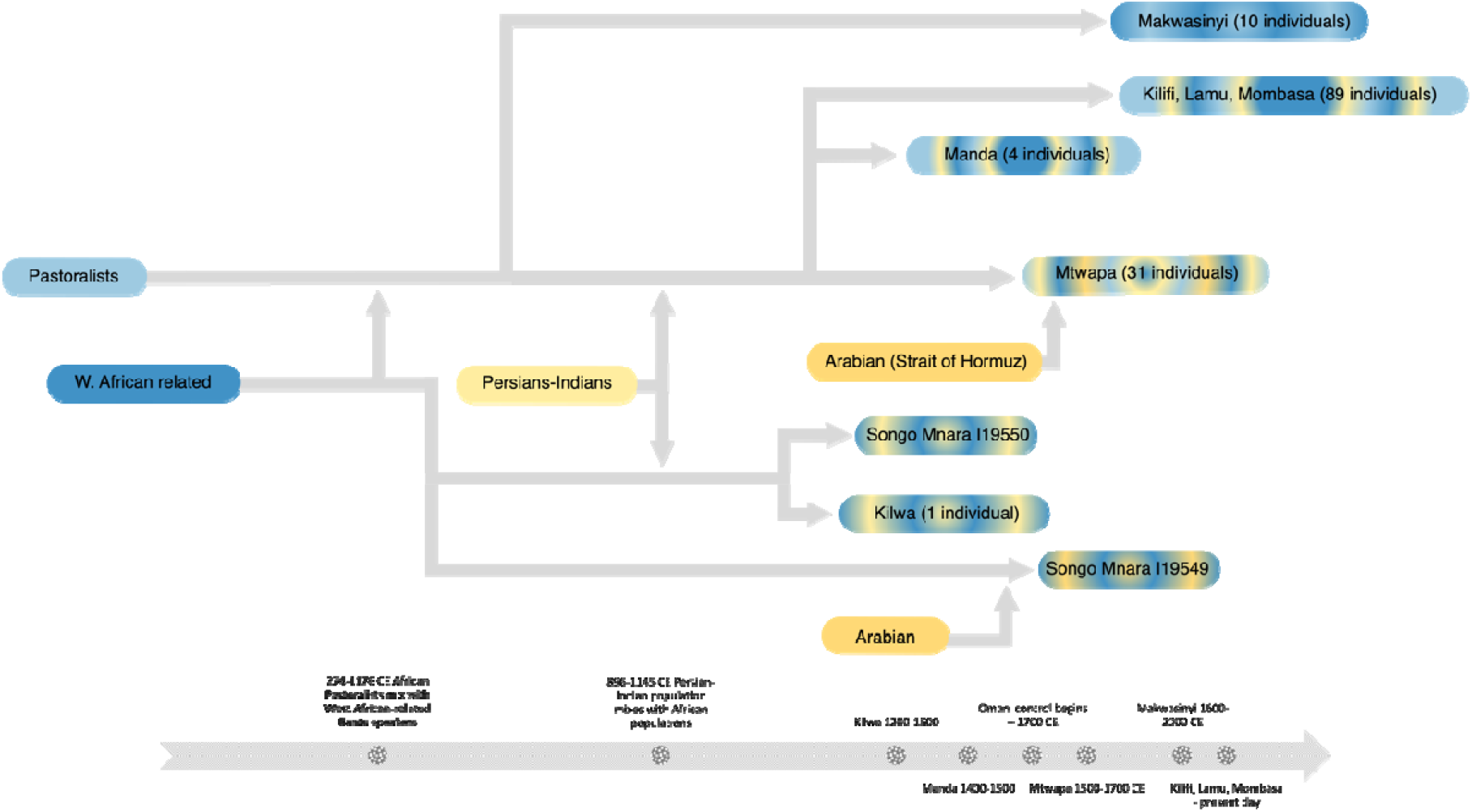
Proposed admixture events of groups along the eastern African coast. African populations are depicted in variations of blue. Near eastern and south Asian populations are depicted in variations of yellow. Mixtures of different colors represent populations that are admixed between the respective proxy source populations with those colors.

The genetic findings are important to interpret in light of the archaeological evidence. The samples analyzed here are from the 14^th^ – 18^th^ centuries, and mostly from elite contexts. While it is notable that elite burials of this period had significant Persian ancestry, we do not fully understand the contexts in which this admixture first occurred. Coastal sites of around 1000 CE had less evidence for elite distinction, and it is notable that these later elites developed from admixed populations. Two of the sites (Mtwapa and Songo Mnara) did not exist as towns during the period of admixture, and so these admixed populations moved to those towns during the succeeding centuries.

The genetic findings are also important to interpret in light of the linguistic evidence. Kiswahili is a Bantu language, and given our finding that most of the female ancestry of individuals in this study derived from Africans, our results point to a scenario in which the children of immigrant men of Asian origin adopted the languages of their mothers, as is common in matrilocal cultures [55]. But Kiswahili also has non-African influences consistent with the genetic and archaeological findings and reflecting 1500 years of interaction with societies around the Indian Ocean rim. Persian loanwords contribute up to 3% of Kiswahili, but it is unclear if they derive directly from Persian or through adoption into other Indian Ocean languages [56]. Arabic loanwords are the single largest non-Bantu element in Kiswahili (16-20%) [56]. However, these words, which predominantly relate to religion, jurisprudence, trade, and maritime vocabulary, may be primarily due to incorporations after around 1500 CE [25].

A repeated theme in this study is the different participation of males and females in population mixture events both on the coast and inland throughout the medieval period. For example, we find evidence of predominantly male Near Eastern ancestors mixing with predominantly female African and, to a much lesser extent, female Indian ancestors in the lineages of medieval Swahili coast people. This finding testifies to asymmetric social interactions between groups as cultural contact occurred, although genetic data cannot determine the processes that contributed to these patterns. We can learn more about the nature of these interactions through archaeological, ethnographic, linguistic, and historical evidence.

A limitation of this work is its sparse sampling of present-day and ancient populations both in space and time. The shortage of present-day and ancient samples means that the data that we have access to may not represent the precise sources or precise regions of Persia or Arabia, but rather be proxies for the true sources. The sampling of ancient Africans is similarly limited. The geographical coverage is skewed toward Kenya—with the individuals from Tanzanian sites such as Songo Mnara, Kilwa, and Lindi sufficient to identify similarities and differences in ancestry profiles from Kenya but not sufficient to define a general pattern—and does not incorporate groups from present-day Somalia, Mozambique, the Comoros Islands, and Madagascar. In addition, the individuals we analyzed were not fully representative of all social and economic groups in Swahili society. Nearly all the coastal graves and tombs occupied prominent positions within medieval and early modern townscapes (see **Supplementary Archaeological site summaries**). With the exceptions of Makwasinyi and possibly Kilwa and Lindi, the individuals we analyzed were from elite contexts within high-profile coastal sites. However, the Swahili cultural world included many non-stone and non-elite settlements whose residents are not represented here and whose ancestry would potentially be systematically different [57], and indeed modern Swahili populations retain differences today. Finally, none of the burials we analyzed pre-dated the 13^th^ century; sampling of individuals who lived before and after the mixture events around 1000 CE would provide insights about the population changes that occurred that cannot be discerned based on the datasets we analyze here.

Narratives of ancestry have a complex history on the eastern African coast. The data presented here add to that complexity. Ancient DNA data shows that there was also a complicated and ongoing cultural mix on the coast, where African people and society interacted and had families with visitors from other parts of Africa and many parts of the Indian Ocean world.

## Methods

### Ethics Statement

The present-day communities of the Swahili coast have strong traditions of connection to the people of the medieval coastal towns—including a tradition of descent from the people who lived in these communities, as well as shared language and religion—and hence community consultation is an important part of this work. Many medieval and present-day coastal people are also Muslim, and hence it is important to carry out any analysis in ways that are sensitive to Muslim proscriptions against disturbing the dead. The sampling for this study emerged from decades’-long community-based archaeology projects—including by co-authors Sibel and Chapurukha Kusimba in the Kenya region, and by co-authors Wynne-Jones, Fleisher, Tanzanian Antiquities, and the Songo Mnara ruins committee in the Tanzania region—that involved participation in and return of results to local communities, following rules for handling and reburying of remains agreed by the communities. Prior to submission of this study for publication, the corresponding authors of this study held return-of-results consultation meetings with local groups in Lamu, Songo Mnara, and Kilwa Kisiwani, and feedback from these engagements helped us to ensure that the final research was reflective of community perspectives.

### Ancient DNA sample preparation, DNA extraction, library preparation, enrichment, sequencing, and bioinformatic analysis

To generate the genetic data for the samples in **Extended Data Table 1** and **Online Table 2 | newly_reported_individuals** (and by-library metrics in **Online Table 1 | newly_reported_libraries**), we used established protocols in dedicated clean rooms, involving first sampling about 40 milligrams of powder from skeletal remains, then extracting DNA using methods designed to retain short, damaged fragments [58], then building individually barcoded libraries [59–61] after incubation with uracil DNA glycosylase (UDG) with the goal of removing characteristic errors typical of ancient DNA in the interiors of sequences [62, 63]. We amplified these libraries and carried out in-solution enrichment of DNA libraries for about 1.2 million single nucleotide polymorphisms (SNPs) [64–66] as well as baits targeting the mitochondrial genome [67], and sequenced the enriched products along with a small amount of unenriched library on Illumina NextSeq 500 or HiSeq X Ten instruments. The resulting sequenced paired-end reads were separated into respective libraries and stripped of identification tags and adapter related sequences. Read pairs were merged prior to alignment, requiring a minimum of 15 base pair overlap, allowing one mismatch if base quality was at least 20 or, up to three mismatches at lower base quality. We restricted to sequences that were at least 30 base pairs long. The resulting sequences were aligned using *samse* from *bwa-v0.6.1* [68] separately to (a) the human genome reference sequence (*hg19*), and (b) the mitochondrial RSRS genome [69] to allow the targeted nuclear and mitochondrial specific sequences to be assessed.

We performed a series of quality control measures. Specifically, we inferred the rate of mismatch of the sequences mapping to the mitochondrial consensus sequence (of libraries with at least 2-fold coverage) and tested if the upper bound of the inferred match rate was at least 95% [64]; we tested if the ratio of Y chromosome to the sum of X and Y chromosome sequences was in the range expected for DNA entirely from a female (<3%) or male (>35%); we tested if the rate of cytosine to thymine substitution in the final nucleotide was at least 10% for single stranded libraries or 3% for double stranded libraries as expected for authentic ancient DNA; and we examined the rate of polymorphism on the X chromosome to estimate a rate of contamination flagging an individual as appreciable contaminated if the lower bound of the inferred contamination rate was at least 1% [70].

We represented each position in the genome based on a single randomly selected sequence, and 80 individuals passed quality control screening. The analyses in this study focus on the 54 non-outlier individuals that are not first- or second-degree relatives of another individual in the study for which we have better quality data and for which we have data on a minimum of 15,000 SNPs. **Online Table 1 | newly_reported_libraries** provides a full report of sequencing results on both libraries that yielded data passing standard measures of ancient DNA authenticity and libraries that failed screening; we report negative results as a guide to help in future sampling efforts.

### Human Origin sample collection and data generation

We generated genotype data for 29 present-day individuals from the United Arab Emirates and Madagascar on the Affymetrix Human Origins array [34]. We applied quality control analyses from [32].

The Madagascar samples analyzed in this study were collected during 2007–2014 with ethical approval by the Human Subjects’ Ethics Committees of the Health Ministry of Madagascar and by French committees (Ministry of Research, National Commission for Data Protection and Liberties and Persons Protection Committee). Individuals all gave written consent before the study. DNA was purified from saliva using the Oragen Kit (DNA Genotek Inc.). The north and south Madagascar sampling was based on a grid sampling approach, in which “spots” 50 km in diameter were placed all over Madagascar (taking into account population density data), and three to four villages were sampled in each spot. Sampled villages were founded before 1900, and sampled individuals were 61 ± 15 y old, with the maternal grandmother and paternal grandfather born within a 50-km radius of the sampling location. Subjects were surveyed for current residence, familial birthplaces, and a genealogy of three generations to establish lineage ancestry and to select unrelated individuals. Global Positioning System locations were obtained during sampling.

The Emirati samples were collected from Emirati nationals from the city of Al Ain, United Arab Emirates. Ethical approval was obtained from the Al Ain District Human Research Ethical Committee. The samples were collected from healthy adults and initially used as controls for a study of rare disease using a different SNP array [71].

A list of the newly reported HO individuals and some relevant information can be found in Online Table 4 | newly_reported_HO_data.

### Principal component analysis (PCA)

We used smartpca version 18180 from EIGENSOFT version 8.0.0 [72, 73] with optional parameters (numoutlieriter: 2, numoutevec: 3, lsqproject: YES, newshrink: YES, and hiprecision: YES). We computed eigenvectors of the covariance matrix of SNPs from present-day individuals from Africa and Eurasia genotyped on the Affymetrix Human Origins (HO) SNP array which targets approximately 600,000 SNPs that are a subset of those targeted by in-solution enrichment (see SI for a list of populations used). Ancient individuals and present-day individuals (most genotyped on the HO SNP array, and some also genotyped on the Illumina Human Omni5 Bead Chip [28]) were projected into the three-dimensional (3D) eigenspace determined by the first three principal components.

### ADMIXTURE analysis

For the ADMIXTURE [74] plots in **Figure 1C** and **Extended Data Figure 1**, we use a dataset of 728 populations containing 7279 non-American Human Origins present-day populations and Swahili coast samples that were converted with convertf from the PACKEDANCESTRYMAP format to the PACKEDPED format for pruning in PLINK2 [75]. We applied the maf 0.01 option, which only includes SNPs that have a minor allele frequency of at least 0.01. We then pruned SNPs based on LD, using the indep-pairwise option with a window size of 200 variants, a step size of 25 variants, and a pairwise *r*^2^ threshold of 0.4. After pruning the SNP dataset, we ran 4 replicates of ADMIXTURE with random seeds with *K* = 4 to *K* = 12 ancestral reference populations.

### Quantitative estimation of mixture proportions

We used *qpAdm* (ADMIXTOOLS version 7.0.2) [34] to test if the Makwasinyi, Manda, Mtwapa, Kilwa, and Songo Mnara target populations can be formally modeled as having derived from a set of source populations (termed “left” groups) relative to a set of reference populations (termed “right” groups). See **Supplementary qpAdm modeling** for further details of this process that are population- or sample-specific.

The *qpAdm* workflow builds on the *qpWave* framework. The null hypothesis of *qpWave* is that a set of left populations (*L*_1_, *L*_2_, … *L*_*N_L_*_) and a set of right populations (*R*_1_, *R*_2_, … *R*_*N_R_*_) form clades with respect to one another. *qpWave* builds a matrix of *f*_4_(*L*_*i*_, *L*_*j*_, *R*_*m*_, *R*_*n*_) that encompass all non-redundant *f_4_*-statistics for the left and right population sets. If the left groups form clades with respect to the right groups, then these *f_4_*-statistics will be statistically consistent with zero. The rank of the matrix will be 1 since any additional rows or columns will not provide any linearly independent information. But if there is gene flow between the left and right groups, then the null hypothesis is rejected (we use a *p* < 0.1 threshold) and the rank of the matrix will increase for every independent gene flow between right and left groups.

*qpAdm* builds on *qpWave* by adding a target population to the left groups. The assumption is that the left groups and right groups still form a clade with respect to one another after the addition of this target; in other words, there is no independent gene flow between the target and the right populations that does not also pass through a common ancestral population with other left groups. The *qpAdm* P value for fit tests if this assumption is met, that is, if there is no *qpWave* rank increase after adding the target population. It then models the target as a linear combination of *f_4_*-statistic vectors associated with the other left groups, normalizing the weights to sum to one in which case they can be interpreted as unbiased estimates of mixture proportions.

In our use of *qpAdm,* we use a cycling approach, treating the target as a linear combination of all possible subsets of the candidate source populations, and moving the other candidate source population to the right. Cycling populations to the right allows us to test if a proposed set of left source populations is consistent with being more closely related to the target than other populations. Thus, we can build the closest admixture model within the constraints of our dataset and test its fit to the data. *qpAdm* uses a Block Jackknife, removing a different block of SNPs in each iteration, to determine the mean coefficient and its standard error for each source population in the model.

### mtDNA and Y-chromosome haplogroup determination

Mitochondrial haplogroups were determined using HaploGrep 2 [76]. Y chromosome haplogroups were determined according to the Yfull tree.

### Sex-bias determination based on the different autosome and X chromosome representation in the genome of males and females

We measured Sex-bias per source population by comparison of the inferences of proportions of ancestry on chromosomes 1-22 (the autosomes) which reflect 50% female and 50% male inheritance (*C_A_* = (*m* + *f*)/2), and on the X chromosome which reflect 67% female and 33% male ancestry (*C_X_* = (2*f* + *m*)/3). We determined Z-scores as in [77]. For each of Mtwapa, Manda, Kilwa, and Makwasinyi, we determine sex-bias in the following manner. (1) We sample 10^6^ sets of coefficients by generating random numbers from a multivariate normal distribution based on the *qpAdm*-determined jackknife mean coefficients and the error covariance matrix for both the autosomes the X-chromosome, separately. Since the values in the covariance matrix from *qpAdm* were rounded, we ensure that the eigenvalues of the matrices are all greater or equal to zero by adding a small offset to the matrices determined as the absolute value of its respective minimum eigenvalue. (2) We remove from consideration all infeasible sets of coefficients, namely those that include a coefficient below 0. (3) We take 95% confidence intervals of all the sampled sets of ancestry proportions. To do so, we determine how extreme each set of proportions are by finding their distance to the jackknife mean coefficients. We remove the 2.5% of samples with the greatest distance and the 2.5% of samples with the least distance. (4) We next calculate the female and male proportions of ancestry for each source population. Given the coefficient *C_A_* on the autosomes and the coefficient *C_X_* on the X chromosome, we can determine the likely proportion of female and male ancestry from any given source by solving the system of two above equations, giving *f* = 3*C_X_* − 2*C_A_* and *m* = *4C_A_* − 3*C_X_*. We calculate these *f* and *m* proportions for each source population for the sampled set of autosomal and X chromosome coefficients. (5) We normalize the proportions of female and male ancestry as 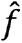 = *f*/(*f* + *m*) and 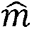 = /*m*(*f* + *m*) for each source for the sampled sets. (6) We find the mean and standard deviation of all the 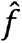 and 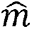 values from the sampled set for each of the source populations. The values are recorded as ranges in **Table 1**. Under the simple model of all the mixture occurring at once, these formulae can be interpreted as estimating the fraction of ancestors at the time of mixing that are on the male or female side. However, if the mixture was more gradual, the interpretation is more complicated albeit still informative about sex bias.

### Determining the composition of the two latest source populations in the African-Persian-Indian admixture

We plot all the Mtwapa and Manda individuals on a ternary plot with their respective proportions of Makwasinyi, Iranian, and Indian ancestry. We apply a linear regression to the high coverage individuals from Mtwapa (I19381, I19394, I19384, I19414, I19420, I19413, I19417, I19401, I23662, I17409, I19391, I13611, I17412, I17413, I23558, I19415, I23561, and I21475) and Manda (I7934) as seen in the ternary plot (see **Figure 2**B and **Figure S3**) with an equivalent Cartesian equation of *f*(*x*) = 0.09856 ⸱ *x* + 0.01301. The line intersects the Makwasinyi axis at 98±6% (left circle in **Figure 2**B and **Figure S3**), which would allow for 2±6% Indian ancestry. This is statistically consistent with 100% Makwasinyi and 0% Iranian and 0% Indian. A parsimonious scenario would thus be that the Iranian and Indian ancestry came from the same source population. The other end of the regression line intersects the Indian axis (right circle in **Figure 2**B and **Figure S3**) at 12±5% Indian ancestry and 88±5% Iranian ancestry. We further consolidated this uncertainty by jackknifing the qpAdm analyses and applying a separate linear regression to each bin. Through this we determine that one admixing source population is composed of 98.4±2.5% Makwasinyi ancestry, which is also consistent with a parsimonious model of 100% Makwasinyi ancestry at that end of the cline. The other source population is composed of 12.2±2.8% Indian ancestry with 87.8±2.8% Iranian ancestry (see **Supplementary qpAdm modeling**).

### Determining the date of mixture using admixture linkage disequilibrium

DATES measures the covariance coefficient between each two genome positions in an ancient individual for an admixture model with two source populations, as described in Narasimhan et al. [48]. With multiple individuals, this analysis is applied separately to all individuals and the curves are combined to increase resolution. The statistical power of DATES increases when we use groups with large sample sizes as sources. The surrogate source populations and estimated elapsed generations can be seen in **Extended Data Figure 2**. We use a 28±2 year-to-generation conversion estimate [78] to calculate the average date of the admixture, which is listed in the text and in **Table 1** and in **Extended Data Figure 2**.

### Data availability

The aligned sequences are available through the European Nucleotide Archive, accession PRJEBxxxxx [to be made available upon publication].

## Acknowledgments

We thank Nicole Adamski, Rebecca Bernardos, Brendan Culleton, Ilana Greenslade, Doug Kennett, Matthew Mah, Adam Micco, and Zhao Zhang, for their contribution to data generation, processing, and curation. Esther Brielle was supported by the EMBO Postdoctoral fellowship ALTF 242-2021. Chapurukha M. Kusimba, Sibel B. Kusimba, Sloan Williams, and Janet M. Monge’s research in Kenya was supported by the National Museums of Kenya and the Republic of Kenya through research permits and excavation permits: 0P/13/001/25C 86; MHE & T 13/001/35C264, and NCST/5/C/002/E/543. The bulk of the research at Manda, Mtwapa, and Makwasinyi was carried out when Chapurukha was at the Field Museum of Natural History. We acknowledge generous financial support from the US National Science Foundation SBR 9024683 (1991-3); BCS 9615291 (1996-8); BCS0106664 (2002-04); BCS 0352681 (2003-04); BCS 0648762 (2007-09; BCS-1030081 (2010-12), the US National Endowment for the Humanities (2012-14), the US IIE J. W. Fulbright Sr. Scholars Program 2002-3; 2012), and the National Geographic Society (1996-7). Excavations at Songo Mnara were supported by the National Science Foundation (BCS 1123091) and the Arts and Humanities Research Council (AH/J502716/1). The Madagascar HO sample collection was supported by the MAGE consortium. The ancient DNA work and analysis were supported by National Institutes of Health grant HG012287, by the John Templeton Foundation (grant 61220), by the Allen Discovery Center program, a Paul G. Allen Frontiers Group advised program of the Paul G. Allen Family Foundation, and by a gift from J.-F. Clin. D.R. is an Investigator of the Howard Hughes Medical Institute.

## Author contributions

ESB, JF, SWJ, SW, JM, MEP, DR, and CMK conceptualized the study; ESB, KS, and SM formally analyzed the genetic data; ESB, NB, KC, EC, LI, AL, JO, LQ, KS, NW, FZ, BC, KS, SM, NR, NP, KBA, MMM, SK, and DR were involved in the genetic data investigation; JF, SWJ, JO, GA, AG, AKa, AKw, AM, FKM, FM, EN, CO, KRB, ES, LAG, BRA, TL, DP, CR, JAR, BC, MAM, KBA, MMM, SW, JM, SK, MEP, DR, and CMK provided resources; ESB, JF, SWJ, MEP, DR, and CMK wrote the paper.

## Competing interests

The authors declare no competing interests.

## Supplementary Information

The online version contains supplementary material available at [to be made available upon publication].

## Correspondence and requests

for materials should be addressed to Esther S. Brielle, Jeffrey Fleisher, Stephanie Wynne-Jones, David Reich, and Chapurukha M. Kusimba.

## Peer review information

to be added after review.

## Reprints and permissions information

is available at [to be made available upon publication].

---

**Extended Data Table 1.**
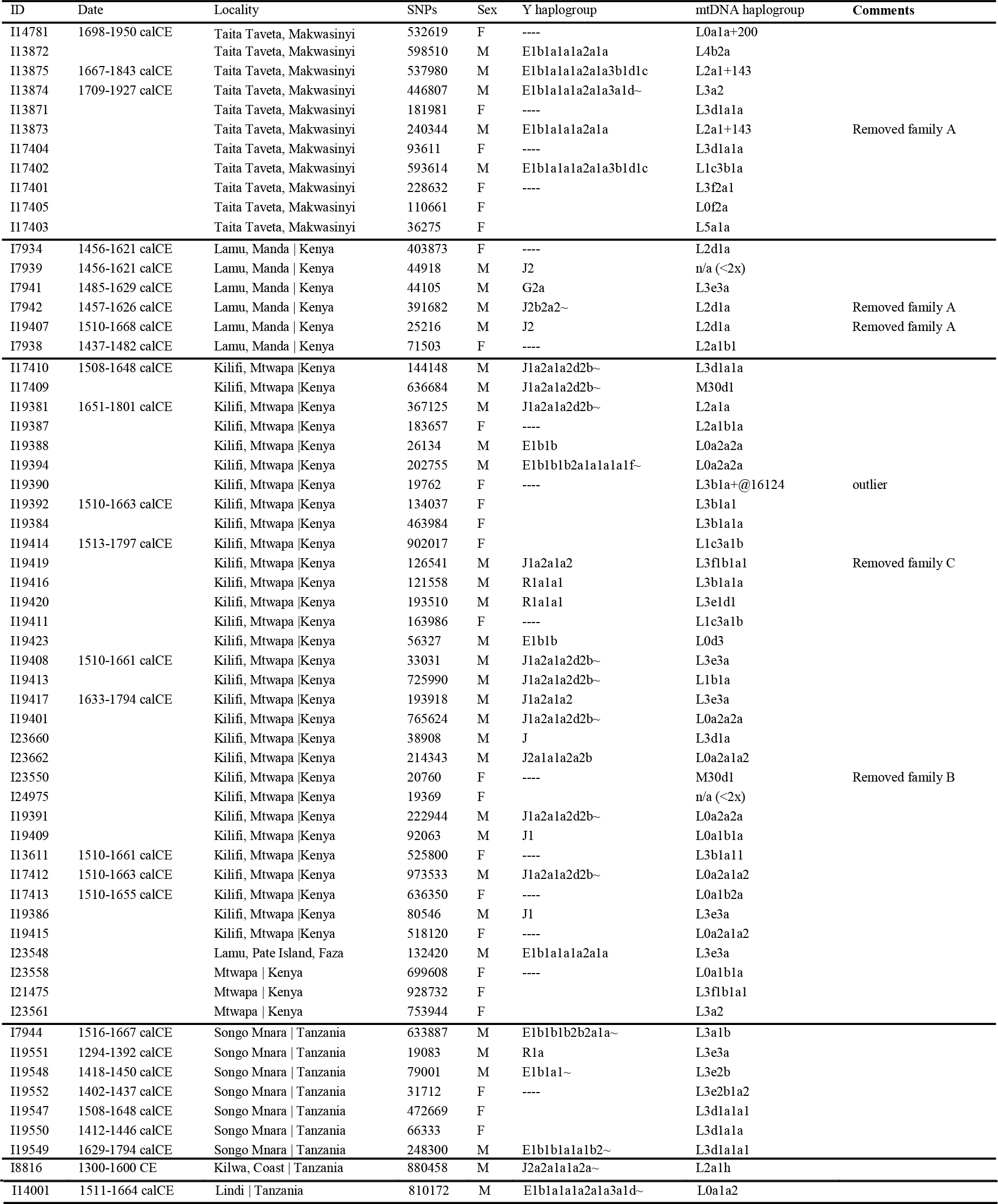
Ancient individuals with high coverage genome-wide data newly reported in this study. Makwasinyi Family A has 4 members. I13873 has a son I13872 and a brother I13875 (who is genetically a second or third degree relative of I13872), and all three males are second- or third-degree relatives of I17402. Manda Family A has 4 members. I7934, I7942, I7943, and I19407 are all first-degree relatives of one another. Mtwapa Family A has 2 members who are second- or third-degree relatives, I19401 and I13611. Mtwapa Family B has 8 members. Of them, I23550 and I17409 are first degree relatives. Second or third-degree relatives are I17409 to I19391, I19394 to I23558, I19394 to I19391, I19391 to I19393, I19394 to I19423, and I19394 to I19388. Mtwapa Family C has 3 members. I21475 and I19419 are first degree relatives. I21475 and I19384 are second- or third-degree relatives. Mtwapa Family D has 2 members who are second- or third-degree relatives, I19416 and I19417. Mtwapa Family E has 2 members who are second- or third-degree relatives, I19414 and I19415. Mtwapa Family F has 2 Members who are second- or third-degree relatives, I23662 and I19398. Songo Mnara Family A has 2 members who second or third-degree relatives, I19547 and I19549.

**Extended Data Table 2.**
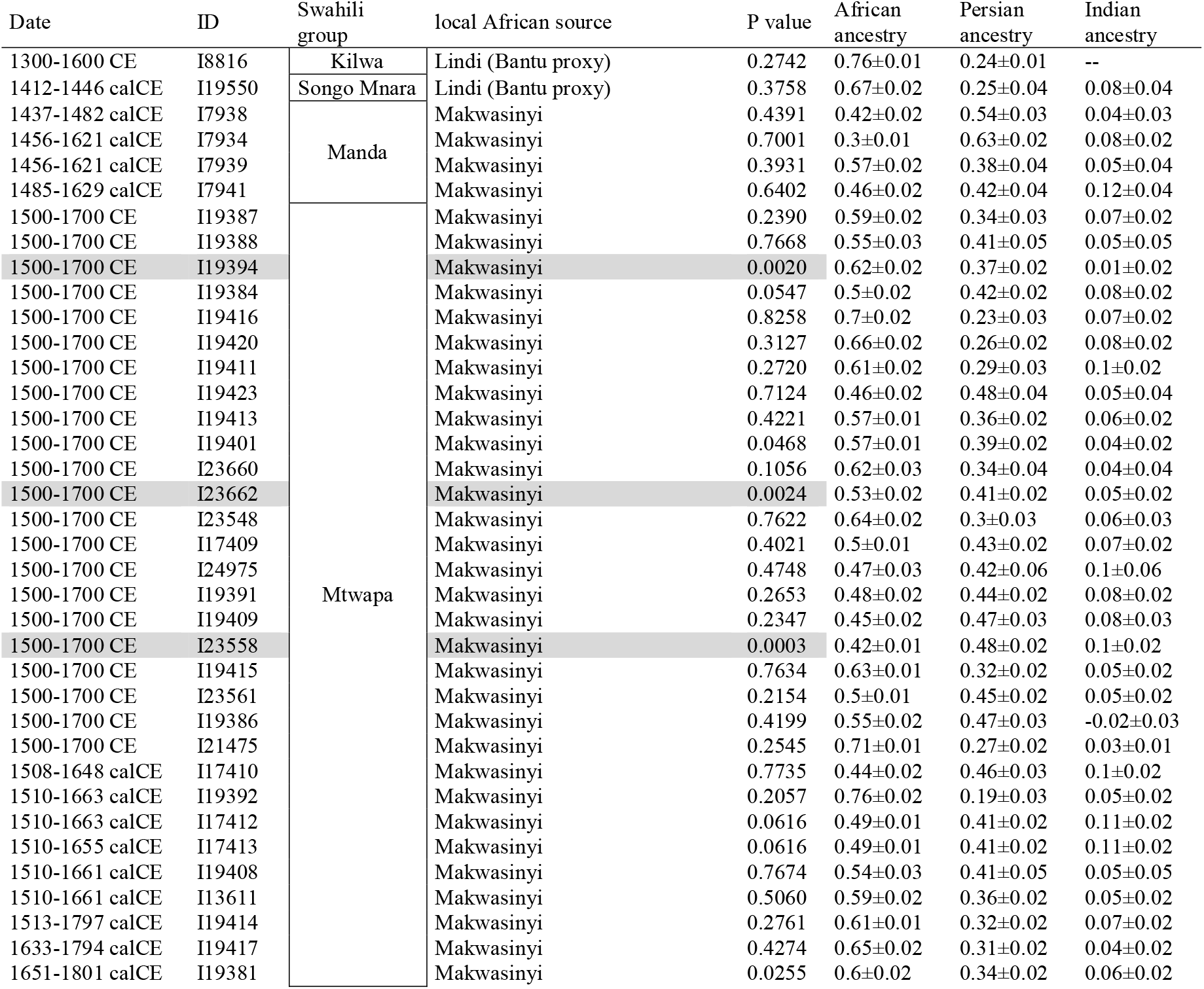
Ancient individuals with high coverage genome-wide data newly reported in this study that have Persian and often Indian ancestry. The African proxy source is the Makwasinyi group from this study for individuals buried at Mtwapa and Manda, and is the Lindi individual from this study for the Kilwa individual and the Songo Mnara individual I19550. The Persian proxy source in all cases is the HO Iranian group [32, 37]. The Indian proxy source in all cases is the HO Sahariya_MP

**Extended Data Table 3.**
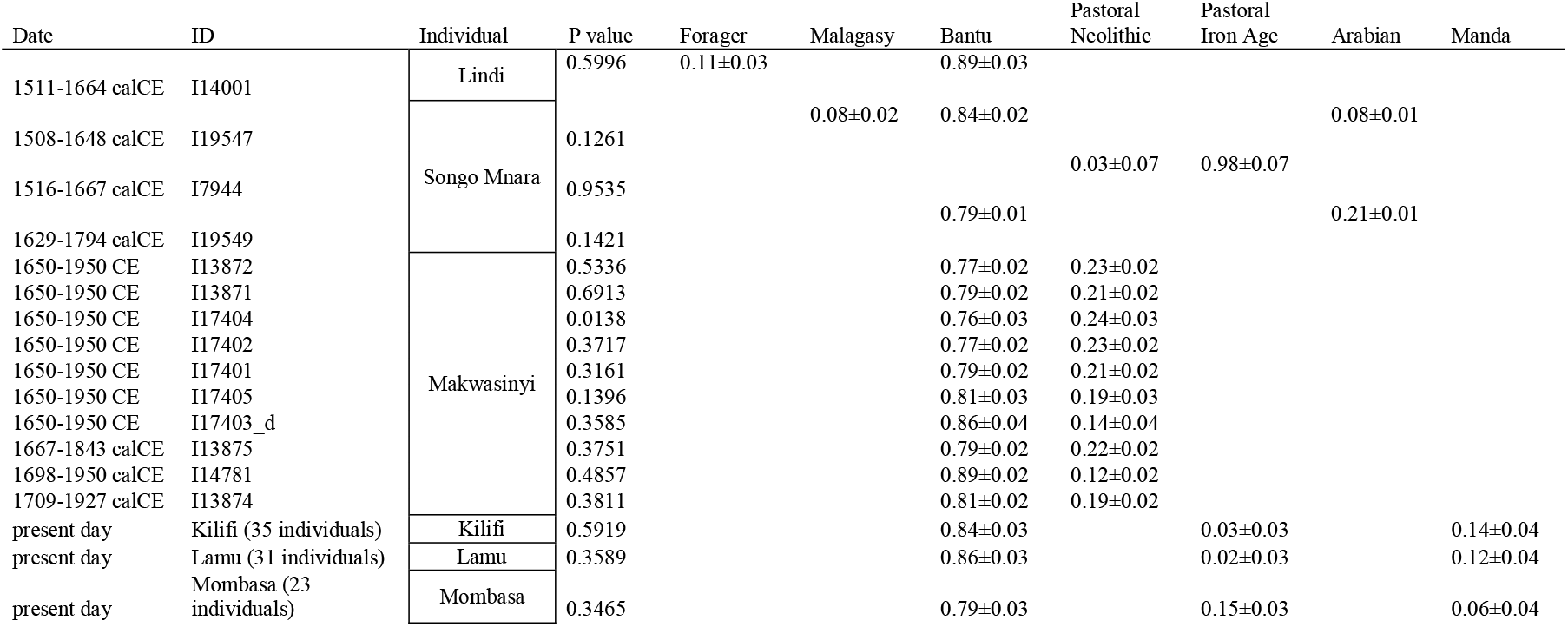
Ancient individuals with high coverage genome-wide data newly reported in this study that do not have evidence of Persian ancestry. The forager proxy source for the Lindi individual I14001 is the Hadza group genotyped on the Human Origins array [32] and the Bantu proxy source is the Malawi_Yao group [31]. The Pastoral Neolithic and Pastoral Iron Age proxy sources for the Songo Mnara individual I7944 are the 1240K capture Kenya_PastoralN group [30] and the 1240K capture Kenya_IA_Pastoral_published group [30], respectively. The Malagasy proxy source for the Songo Mnara individual I19547 is the Madagascar_North group, the Bantu proxy source is the Malawi_Ngoni group, and the Arabian proxy source is the Yemeni_Highlands_Raymah individual. The Bantu proxy source for the Songo Mnara individual I19549 is the 1240K capture Tanzania_Pemba_600BP_published individual [31] and the Arabian proxy source is the Emirati group. The Bantu proxy source for the Makwasinyi group is the 1240K capture Tanzania_Pemba_600BP_published individual [31] and the Pastoral Neolithic proxy source is the 1240K capture Kenya_PastoralN group [30]. The Bantu proxy source for all the present-day groups from Kilifi, Lamu, and Mombasa is the 1240K Lindi individual from the present study. The medieval Swahili-speaking proxy source is the Kilwa individual from the present study for the present-day groups. The Pastoral Iron Age proxy source for the present-day groups is two published individuals from [30], I8892_published and I12381.

**Extended Data Figure 1.**
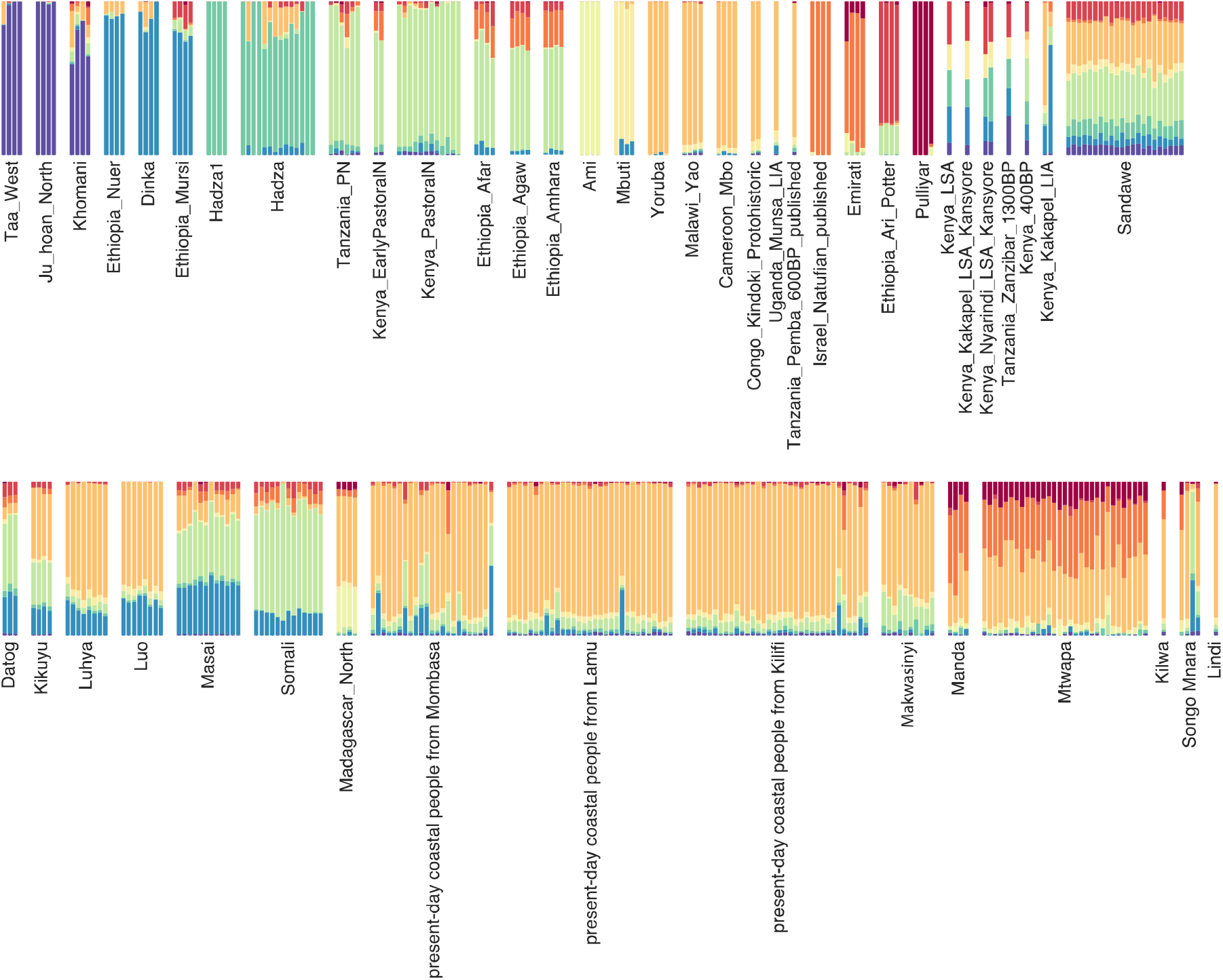
ADMIXTURE analysis, with K=10 ancestral reference populations. We used 262 populations (2248 individuals) including African populations genotyped with the Human Origins array along with some 1240K capture ancient samples from present-day Kenya, present-day Tanzania, and surrounding areas, Swahili coast present-day samples [28], ancient Israel_Natufian_published, and Pulliyar samples from present-day India. We were able to view a variety of African pastoral, farming, and forager ancestries within coastal and inland Swahili-speaking populations of the Human Origins dataset. Makwasinyi components of ancestry resemble those of the present-day coastal populations sampled by Brucato and colleagues [28], albeit with different proportions.

**Extended Data Figure 2.**
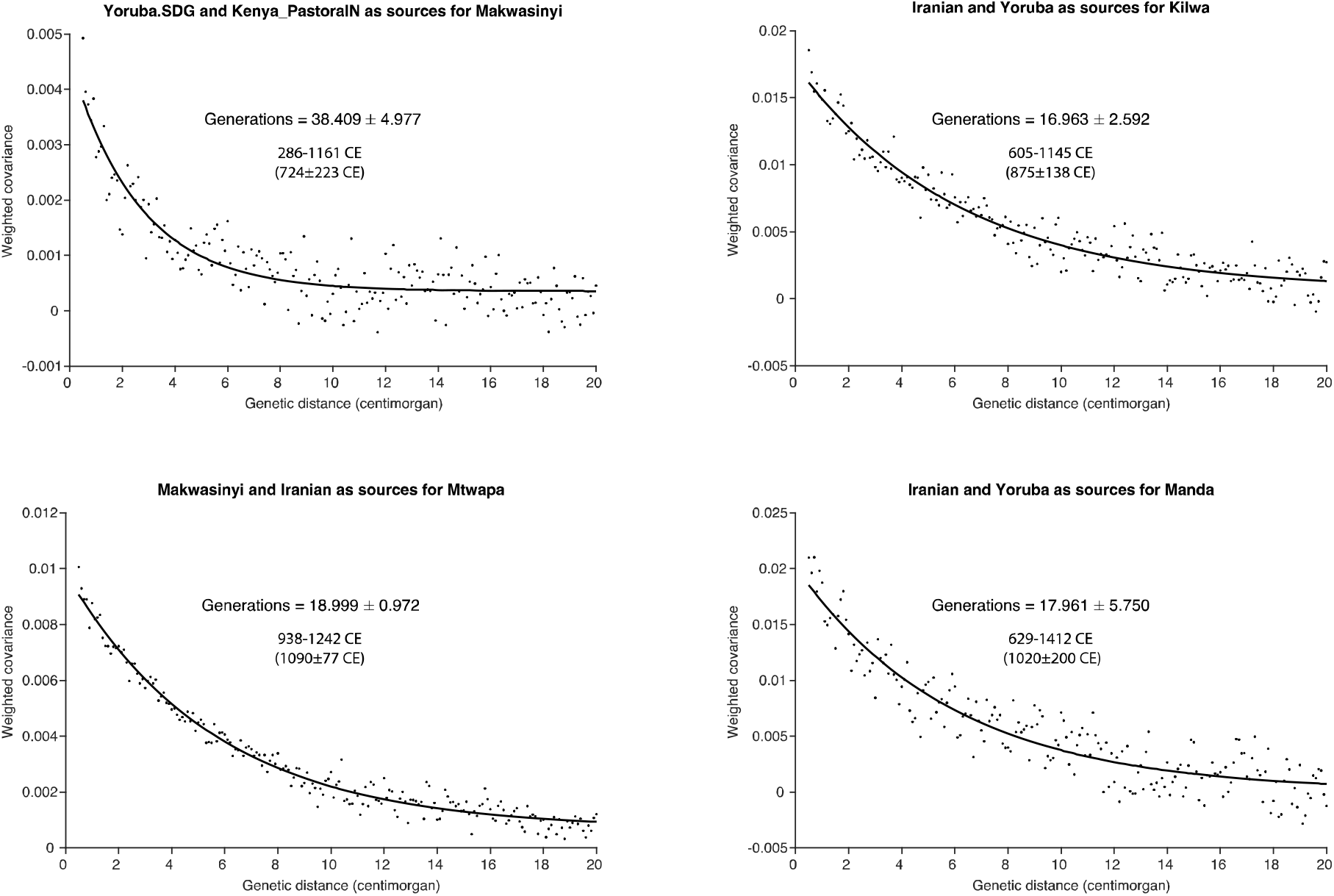
DATES curves. Curves that show the exponential decay of linkage disequilibrium generated by mixture between two populations related differentially to the two sources as a function of the number of elapsed generations since the mixture event. Multiplying by a 28±2 year-to-generation conversion estimate and subtracting from the average calibrated radiocarbon dates of the ancient individuals (or the archaeologically estimated date) gives calendar years [78].

**Extended Data Figure 3.**
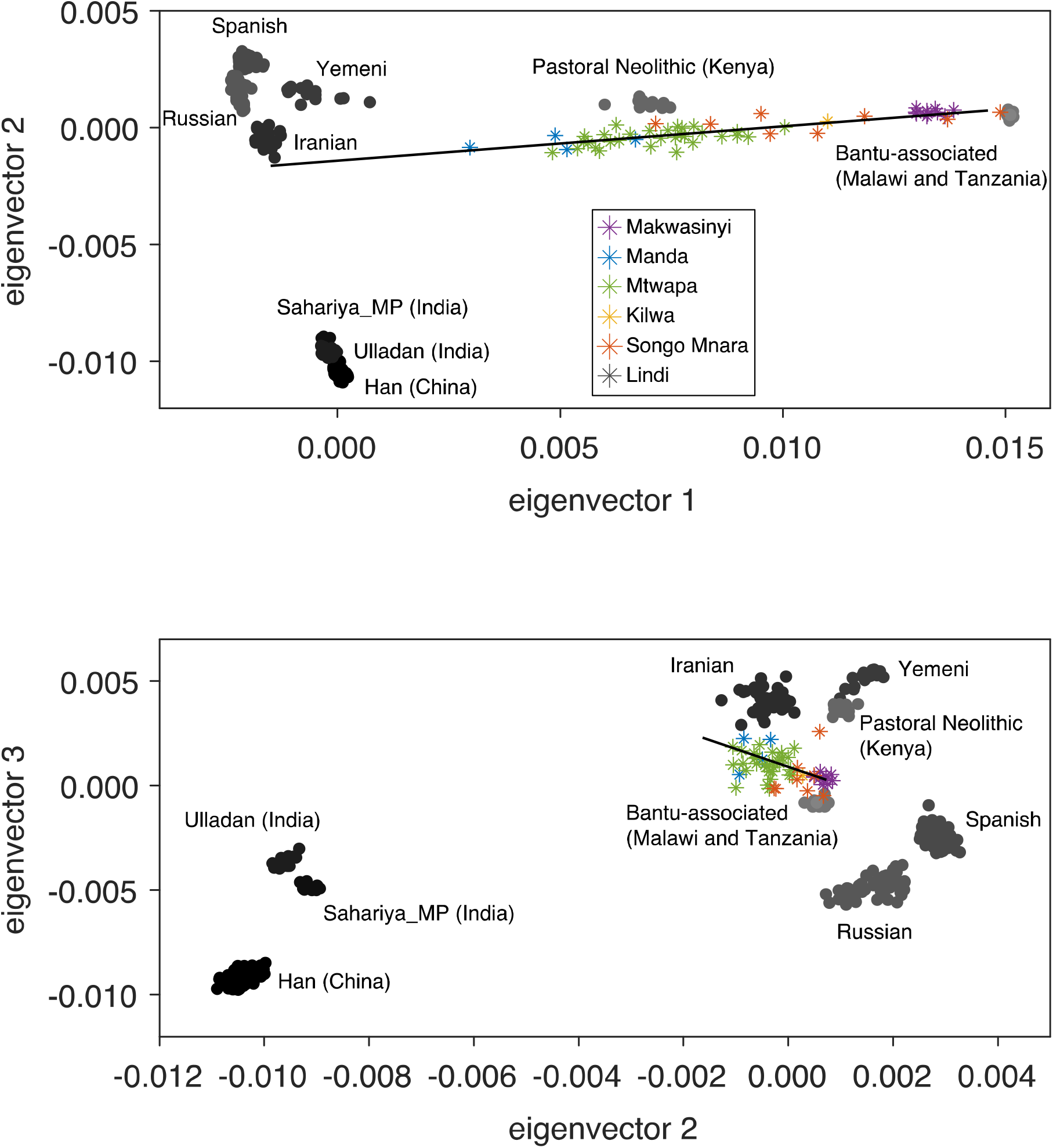
Two dimensional PCAs. (Top) A PCA of the first and second eigenvectors for the same data as in Figure 1B. (Bottom) A PCA of the second and third eigenvectors for the same data as in Figure 1B.

## Notes

### Competing Interest Statement

The authors have declared no competing interest.

## References

1. Wynne-Jones, S. and A.J. LaViolette, *The Swahili World*. Routledge Worlds Series. London: Routledge.

2. Ichumbaki, E.B. and E. Pollard, The Swahili Civilization in Eastern Africa, in Oxford Research Encyclopedia of Anthropology. 2021, Oxford University Press.

3. Chami, F., The Tanzanian Coast in the Early First Millennium AD: an archaeology of the iron-working, farming communities. Studies in African Archaeology. Vol. 7. 1994, Uppsala: Societas Archaeologica Uppsaliensis.

4. Fleisher, J. and S. Wynne-Jones, Ceramics and the early Swahili: deconstructing the Early Tana Tradition. African Archaeological Review, 2011. 28(4): p. 245–278.

5. Pollard, E. and O.C. Kinyera, The Swahili Coast and the Indian Ocean Trade Patterns in the 7th–10th Centuries CE. Journal of Southern African Studies, 2017. 43(5): p. 927–947.

6. Horton, M.C. and J. Middleton, The Swahili: The Social Landscape of a Mercantile Society. 2001, Oxford: Blackwell.

7. Horton, M.C., Shanga:An early Muslim trading community on the coast of Eastern Africa. Vol. 14. 1996: British Institute in Eastern Africa.

8. Fleisher, J., et al., When Did the Swahili Become Maritime? American Anthropologist, 2015. 117(1): p. 100–115.

9. Wright, H., Trade and Politics on the Eastern Littoral of Africa, AD 800-1300, in The Archaeology of Africa: Food, Metals and Towns, A.O. Bassey Andah, Thurstan Shaw, and Paul Sinclair, Editor. 1993, Routledge: London.

10. Kusimba, C.M., *The Rise and Fall of Swahili States*. 1999, Walnut Creek: Altamira Press.

11. Pearson, M.N., Port cities and intruders: The Swahili coast, India, and Portugal in the early modern era. 1998.

12. Vernet, T., Le territoire hors les murs des cités-États swahili de l’archipel de Lamu, 1600-1800. Journal des africanistes, 2004. 74(1-2): p. 381–411.

13. Sheriff, A., Slaves, Spices and Ivory in Zanzibar. Integration of an East African Commercial Empire into the World Economy, 1770-1873. 1987, Oxford: James Currey.

14. Glassman, J., Feasts and Riot: Revelry, Rebellion, and Popular Consciousness on the Swahili Coast 1856-1888. 1995, London: James Currey.

15. Spear, T.T., The Shirazi in Swahili Traditions, Culture, and History. History in Africa, 1984. 11: p. 291–305.

16. Tolmacheva, M., The Pate Chronicle: Edited and Translated from Mss 177, 321, 344, and 358 of the Library of the University of Dar Es Salaam. 1993, Ann Arbor, MI: University of Michigan Press.

17. Pawlowicz, M. and A. LaViolette, Pawlowicz M. and LaViolette, A. 2013. Swahili Historical Chronicles from an Archaeological Perspective: Bridging History and Archaeology, and Coast and Hinterland, in Southern Tanzania. In P. Schmidt and S. Mrozowski (Eds.), The Death of Prehistory (pp. 117–140). Oxford: Oxford University Press. 2013. p. 117–140.

18. Freeman-Grenville, G.S.P., The East African Coast. Select Documents from the First to the Earlier Nineteenth Centuries. 1962, London: Clarendon Press.

19. Kirkman, J.S., The Arab City of Gedi: Excavations at the Great Mosque, Architecture and Finds. 1954, London: Oxford University Press.

20. Chittick, H.N., *The Peopling of the East African coast*, in *East Africa and the Orient: Cultural Syntheses in Pre-Colonial Times*, H.N. Chittick and R.I. Rotberg, Editors. 1975, Africana Publishing Company: New York & London. p. 16-43.

21. Pouwels, R.L., *Horn and Crescent. Cultural change and traditional Islam on the East African coast, 800-*1900. African Studies Series. Vol. 53. 1987, Cambridge: Cambridge University Press.

22. Saad, E., Kilwa Dynastic Historiography: A Critical Study. History in Africa, 1979. 6: p. 177–209.

23. Allen, J.d.V., Swahili Origins: Swahili Culture and the Shungwaya Phenomenon. 1993, Ohio U Press: Athens.

24. Nurse, D. and T.J. Hinnebusch, *Swahili and Sabaki: A Linguistic History*. 1993, Berkeley: University of California Press.

25. Nurse, D. and T. Spear, The Swahili: Reconstructing the History and Language of an African Society, 800-1500. 1985: University of Pennsylvania Press.

26. Crowther, A., et al., Subsistence mosaics, forager-farmer interactions, and the transition to food production in eastern Africa. Quaternary International, 2018. 489: p. 101–120.

27. Caplan, P., ‘But the coast, of course, is quite different’: Academic and Local Ideas about the East African Littoral. Journal of Eastern African Studies, 2007. 1(2): p. 305–320.

28. Brucato, N., et al., The Comoros Show the Earliest Austronesian Gene Flow into the Swahili Corridor. American journal of human genetics, 2018. 102(1): p. 58–68.

29. Raaum, R.L., et al., Decoding the genetic ancestry of the Swahili, in The Swahili World. 2017, Routledge. p. 81–102.

30. Prendergast Mary, E., et al., Ancient DNA reveals a multistep spread of the first herders into sub-Saharan Africa. Science, 2019. 365(6448): p. eaaw6275.

31. Skoglund, P., et al., Reconstructing Prehistoric African Population Structure. Cell, 2017. 171(1): p. 59–71.e21.

32. Lazaridis, I., et al., Ancient human genomes suggest three ancestral populations for present-day Europeans. Nature, 2014. 513(7518): p. 409–413.

33. Nakatsuka, N., et al., The promise of discovering population-specific disease-associated genes in South Asia. Nature Genetics, 2017. 49(9): p. 1403–1407.

34. Patterson, N., et al., Ancient Admixture in Human History. Genetics, 2012. 192(3): p. 1065–1093.

35. Biagini, S.A., et al., People from Ibiza: an unexpected isolate in the Western Mediterranean. European journal of human genetics : EJHG, 2019. 27(6): p. 941–951.

36. Broushaki, F., et al., Early Neolithic genomes from the eastern Fertile Crescent. Science, 2016. 353(6298): p. 499–503.

37. Lazaridis, I., et al., Genomic insights into the origin of farming in the ancient Near East. Nature, 2016. 536(7617): p. 419-424.

38. Jeong, C., et al., The genetic history of admixture across inner Eurasia. Nature Ecology & Evolution, 2019. 3(6): p. 966–976.

39. Kusimba, C.M., S.B. Kusimba, and L. Dussubieux, Beyond the Coastalscapes: Preindustrial Social and Political Networks in East Africa. African Archaeological Review, 2013. 30(4): p. 399–426.

40. Kusimba, C.M. and J.R. Walz, When Did the Swahili Become Maritime?: A Reply to Fleisher et al. (2015), and to the Resurgence of Maritime Myopia in the Archaeology of the East African Coast. American Anthropologist, 2018. 120(3): p. 429–443.

41. Rowold, D.J., et al., Mitochondrial DNA geneflow indicates preferred usage of the Levant Corridor over the Horn of Africa passageway. Journal of Human Genetics, 2007. 52(5): p. 436–447.

42. Quintana-Murci, L., et al., Where West Meets East: The Complex mtDNA Landscape of the Southwest and Central Asian Corridor. The American Journal of Human Genetics, 2004. 74(5): p. 827–845.

43. Abu-Amero, K.K., et al., Mitochondrial DNA structure in the Arabian Peninsula. BMC Evolutionary Biology, 2008. 8(1): p. 45.

44. Gandini, F., et al., Mapping human dispersals into the Horn of Africa from Arabian Ice Age refugia using mitogenomes. Scientific Reports, 2016. 6(1): p. 25472.

45. Regueiro, M., et al., Iran: Tricontinental Nexus for Y-Chromosome Driven Migration. Human Heredity, 2006. 61(3): p. 132–143.

46. Luis, J.R., et al., The Levant versus the Horn of Africa: Evidence for Bidirectional Corridors of Human Migrations. The American Journal of Human Genetics, 2004. 74(3): p. 532–544.

47. Cadenas, A.M., et al., Y-chromosome diversity characterizes the Gulf of Oman. European Journal of Human Genetics, 2008. 16(3): p. 374–386.

48. Narasimhan Vagheesh, M., et al., The formation of human populations in South and Central Asia. Science, 2019. 365(6457): p. eaat7487.

49. Silva, M., et al., 60,000 years of interactions between Central and Eastern Africa documented by major African mitochondrial haplogroup L2. Scientific Reports, 2015. 5(1): p. 12526.

50. Quintana-Murci, L., et al., Genetic evidence of an early exit of Homo sapiens sapiens from Africa through eastern Africa. Nature Genetics, 1999. 23(4): p. 437–441.

51. Pipek, O.A., et al., Worldwide human mitochondrial haplogroup distribution from urban sewage. Scientific Reports, 2019. 9(1): p. 11624.

52. Rishishwar, L. and I.K. Jordan, Implications of human evolution and admixture for mitochondrial replacement therapy. BMC Genomics, 2017. 18(1): p. 140.

53. Vyas, D.N., A. Al-Meeri, and C.J. Mulligan, Testing support for the northern and southern dispersal routes out of Africa: an analysis of Levantine and southern Arabian populations. American Journal of Physical Anthropology, 2017. 164(4): p. 736–749.

54. Horton, M., et al., The Chronology of Kilwa Kisiwani, AD 800–1500. African Archaeological Review, 2022.

55. Lansing, J.S., et al., Kinship structures create persistent channels for language transmission. Proceedings of the National Academy of Sciences, 2017. 114(49): p. 12910–12915.

56. Haspelmath, M. and U. Tadmor, *Loanwords in the world’s languages : a comparative handbook*. 2009, Berlin, Germany: De Gruyter Mouton.

57. Fleisher, J. and A. LaViolette, *Elusive wattle-and-daub: Finding the hidden majority in the archaeology of the Swahili*. Azania: Archaeological Research in Africa, 1999. 34(1): p. 87–108.

58. Dabney, J., et al., Complete mitochondrial genome sequence of a Middle Pleistocene cave bear reconstructed from ultrashort DNA fragments. Proceedings of the National Academy of Sciences, 2013. 110(39): p. 15758–15763.

59. Gansauge, M.-T., et al., Single-stranded DNA library preparation from highly degraded DNA using T4 DNA ligase. Nucleic acids research, 2017. 45(10): p. e79–e79.

60. Gansauge, M.-T., et al., Manual and automated preparation of single-stranded DNA libraries for the sequencing of DNA from ancient biological remains and other sources of highly degraded DNA. Nature Protocols, 2020. 15(8): p. 2279–2300.

61. Rohland, N., et al., *Partial uracil-DNA-glycosylase treatment for screening of ancient DNA.* Philosophical transactions of the Royal Society of London. Series B, Biological sciences, 2015. 370(1660): p. 20130624-20130624.

62. Rohland, N., et al., Extraction of highly degraded DNA from ancient bones, teeth and sediments for high-throughput sequencing. Nature Protocols, 2018. 13(11): p. 2447–2461.

63. Briggs Adrian, W., et al., Patterns of damage in genomic DNA sequences from a Neandertal. Proceedings of the National Academy of Sciences, 2007. 104(37): p. 14616–14621.

64. Fu, Q., et al., DNA analysis of an early modern human from Tianyuan Cave, China. Proceedings of the National Academy of Sciences of the United States of America, 2013. 110(6): p. 2223–2227.

65. Fu, Q., et al., An early modern human from Romania with a recent Neanderthal ancestor. Nature, 2015. 524(7564): p. 216–219.

66. Rohland, N., et al., Three Reagents for in-Solution Enrichment of Ancient Human DNA at More than a Million SNPs. bioRxiv, 2022: p. 2022.01.13.476259.

67. Maricic, T., M. Whitten, and S. Pääbo, Multiplexed DNA sequence capture of mitochondrial genomes using PCR products. PloS one, 2010. 5(11): p. e14004–e14004.

68. Li, H. and R. Durbin, Fast and accurate long-read alignment with Burrows–Wheeler transform. Bioinformatics, 2010. 26(5): p. 589–595.

69. Behar, Doron M., et al., A “Copernican” Reassessment of the Human Mitochondrial DNA Tree from its Root. The American Journal of Human Genetics, 2012. 90(4): p. 675–684.

70. Korneliussen, T.S., A. Albrechtsen, and R. Nielsen, ANGSD: Analysis of Next Generation Sequencing Data. BMC Bioinformatics, 2014. 15(1): p. 356.

71. Reiff, R.E., et al., METTL23, a transcriptional partner of GABPA, is essential for human cognition. Human Molecular Genetics, 2014. 23(13): p. 3456–3466.

72. Patterson, N., A.L. Price, and D. Reich, Population Structure and Eigenanalysis. PLOS Genetics, 2006. 2(12): p. e190.

73. Price, A.L., et al., Principal components analysis corrects for stratification in genome-wide association studies. Nature Genetics, 2006. 38(8): p. 904–909.

74. Alexander, D.H., J. Novembre, and K. Lange, Fast model-based estimation of ancestry in unrelated individuals. Genome research, 2009. 19(9): p. 1655–1664.

75. Chang, C.C., et al., Second-generation PLINK: rising to the challenge of larger and richer datasets. GigaScience, 2015. 4(1): p. s13742–015-0047-8.

76. Weissensteiner, H., et al., HaploGrep 2: mitochondrial haplogroup classification in the era of high-throughput sequencing. Nucleic Acids Research, 2016. 44(W1): p. W58–W63.

77. Mathieson, I., et al., The genomic history of southeastern Europe. Nature, 2018. 555(7695): p. 197–203.

78. Moorjani, P., et al., A genetic method for dating ancient genomes provides a direct estimate of human generation interval in the last 45,000 years. Proceedings of the National Academy of Sciences, 2016. 113(20): p. 5652.

