## Supplementary Information for "The Entwined African and Asian Genetic Roots of the Medieval Peoples of the Swahili Coast"

### Permissions and sampling protocols

#### Permissions to export samples for destructive analysis and research

*Songo Mnara*

Permission to sample individuals from Songo Mnara was obtained from the United Republic of Tanzania, granted by the Commission for Science and Technology (COSTECH permit 2011-167-ER-2009-46 issued to co-authors J. Fleisher and S. Wynne-Jones) as well as the Ministry of Natural Resources and Tourism (Excavation License 06/2011). Permission to export the skeletal samples was granted by the Ministry of Natural Resources and Tourism, United Republic of Tanzania (Permit No. 5/2011, Ref. No. EA.402/606/01/9).

*Kilwa*

Permissions to sample individuals from Kilwa were obtained from both the United Republic of Tanzania (where the site is located) and the Republic of Kenya (where the skeletal remains were curated at the time of sampling, at the British Institute in Eastern Africa in Nairobi). Permission from Tanzania was granted by the Commission for Science and Technology (COSTECH permits 2017-220/221-NA-2012-50 issued to co-authors M. Prendergast and E. Sawchuk). Permission to sample the remains in Kenya was obtained from the National Commission for Science, Technology, and Innovation (NACOSTI permit P/17/34239/17088 issued to co-author M. Prendergast), through affiliation with the National Museums of Kenya (NMK), as well as by the then-director of the British Institute in Eastern Africa, J. Fontein. Permission to export the skeletal samples was granted by the Cabinet Secretary, Ministry of Sports and Heritage, Republic of Kenya.

*Lindi*

The individual from Lindi was sampled upon request by curators at the National Museums of Tanzania, under a permit from the Commission for Science and Technology (COSTECH permits 2017-220/221-NA-2012-50 issued to co-authors M. Prendergast and E. Sawchuk), through affiliation with the National Museums of Tanzania (NMT), and by NMT curator co-author A. Kwekason. Permission to export the skeletal samples was granted by the Division of Antiquities, Ministry of Natural Resources and Tourism (Export License 03/2018/2019).

*Mtwapa*

Research at Mtwapa was supported by the National Museums of Kenya and the Field Museum (Chicago, Illinois, USA). Permission to sample individuals from Mtwapa was obtained from the Republic of Kenya through research permits and excavation permits: 0P/13/001/25C 86; MHE & T 13/001/issued to co-authors C.M. Kusimba, S. Williams, and J. Monge. After collecting samples from all the individuals, all the human skeletal remains were reburied at Mtwapa.

*Manda*

Research at Manda was supported by the National Museums of Kenya and the Field Museum (Chicago, Illinois, USA). Permission to sample individuals from Manda was obtained from the Republic of Kenya through research permits and permits: NACOSTI/P/17/811175/16914 issued to co-authors C.M. Kusimba, S. Williams, and J. Monge. All the skeletal remains were reburied at the sites following sample collection and study.

*Makwasinyi*

Research at Makwasinyi in Tsavo was supported by the National Museums of Kenya and the Field Museum (Chicago, Illinois, USA). Permission to sample individuals from Makwasinyi was obtained from the Republic of Kenya through research permit NCST/5/C/002/E/543 issued to co-author C.M. Kusimba. The skulls were left intact at their sites after samples were collected.

#### Skeletal sampling protocols, sample return, and curation of derived products

*Songo Mnara*

Samples were selected from excavated skeletal remains at Songo Mnara. Samples from adult burials included 1-2 teeth for isotopic study as well as one metatarsal or metacarpal for aDNA analysis. Infants were not sampled during excavations. One infant mandible recovered during analysis of domestic faunal remains was also subjected to aDNA extraction. Teeth samples are stored at the University of Bristol. Bone samples are currently with the Reich laboratory at Harvard University; after this study is complete, they will be returned to Tanzania via the National Museums of Tanzania.

*Kilwa and Lindi*

For Kilwa and Lindi, samples were selected following protocols outlined by Prendergast and Sawchuk (2018) and Prendergast et al. (2019).

For Kilwa, three individuals were identified by E. Sawchuk among the commingled and largely unlabeled remains. Two samples were collected for one individual (KS.01.01, KS.01.02; the latter was not sampled and was returned intact), and one for each of the other individuals (KS.02.01, KS.03.01). Samples from Kilwa were exported and returned to the NMK by co-author C. Ogola in October 2017. They are due to be repatriated to the National Museum of Kenya during 2022 and permissions have been granted by co-author Dr. A. Kwekason.

For Lindi, two samples were chosen from the single individual present at this site, an adult male: the petrous portion of the left temporal bone (sample number LS1.02); and the right upper central incisor (LS1.01). Only the petrous was sampled, and the remainder of this petrous, and the complete unsampled incisor, were both returned to the NMT by M. Prendergast, along with a complete record of sampling activity and aDNA and C14 results, in May 2019. As described in a Memorandum of Agreement (MOA) between the Reich lab and the National Museum of Tanzania, the remaining powder, DNA extracts, and libraries remain under curation at the Reich Laboratory at Harvard University.

*Mtwapa*

Bone and tooth samples were taken over the course of the excavations at the site, which began under the direction of Kusimba in 1996. Most of the samples were collected by Monge and Williams during the 2010 season, however, when excavations focused on the cemetery located near the mosque. Tooth samples were collected for radiocarbon dating, stable isotope, and genetic analyses. Long bone fragments and rib samples were collected for stable isotope analyses. After each excavation season, the human remains were reburied, except for one individual remains are curated at the NMK. Any sample material remaining after analysis either has been or will be returned to Kusimba for curation at the NMK.

*Manda*

All samples were collected during a single excavation season in 2011-2012. Tooth samples were collected by co-authors J. Monge and S. Williams for radiocarbon dating, stable isotope, and genetic analyses. Long bone fragments and rib samples were collected for stable isotope analyses. The human remains were reburied the following year when osteological analyses were completed. Any sample material remaining after analysis either has been or will be returned to co-author C.M. Kusimba for return to the NMK.

*Makwasinyi*

Tooth fragment samples were collected by co-author C.M. Kusimba. The crania were left undisturbed in their respective localities. Any remaining sample material has been returned to co-author C.M. Kusimba for return to the NMK.

### Archaeological site summaries

Latitude in degrees, longitude in degrees, and altitude in meters above sea level (m asl) are approximate. Calibrated dates (cal BP or cal CE) were modeled in OxCal version 4.3.2 (Bronk Ramsey (2009), using either the IntCal20 (Reimer et al 2020) or SHCal20 (Hogg et al. 2020) calibration latitude depending on whether the site was north (Kenya) or south (Tanzania) of the equator.

*Mtwapa*

*Lat: 3.954S Long: 39.757E*

The medieval town of Mtwapa is located on the Mtwapa Creek some 15 km north of Mombasa, Kenya. It was first reported by Emery [1]. The present site consists of abandoned houses, collapsed walls, and dried water wells. The remains include five mosques and 64 houses, two workshops, four possible commercial houses, and 20 mounds of undetermined structures [2]. The stone town section of Mtwapa covered approximately eight hectares, out of which four still contain standing architecture. A team involving multiple co-authors of this study surveyed, mapped, and excavated 15 trenches at nine localities and two cemeteries at Mtwapa since 1986. The town’s chronological context, drawn from more than 70 radiocarbon dates, dates the site from 1732 BC to 1750 CE when Mtwapa was abandoned [3].

The diversity in residential structures, ranging from mud structures mostly located outside the perimeter wall to a cluster of stone houses of varying degrees of size and different materials, attests to a city differentiated by wealth. The stone town itself shows significant class differences. Complementarity in subsistence strategies and economic activities are visible in consumption patterns, whereas differences in households are difficult to discern because of similarities in the distribution of fish and mammalian bone remains.

Archaeological excavations recovered large numbers of diverse artifacts typical of urban society. The finds, including local and trade ceramics, iron and iron slag, rock crystal, spindle whorls, glass, marine and Indo-Pacific beads, reveal a complex organized multi-tiered urban polity with a thriving domestic, regional, and international economy [3]. The ubiquitous local pottery belongs to what Chami [4, 5] has referred to as Zanjian and post-Zanjian ware, but is interchangeably referred to as Tana ware [6-8]. Mtwapa’s local pottery was produced at the household level by potters who exploited clay sources located upstream along the creek. The diverse yet conservative forms, styles, and decorative motifs of local pottery point to habits and traditions in food production dominated by boiling, stewing and only secondarily, baking [9]. In its development from a seasonal camp in ca. 1732 BCE to its abandonment ca. 1750 CE, Mtwapa residents consumed mostly marine animal resources, including fishes and mollusks.

There are 99 skeletons at various stages of preservation from Mtwapa excavated from 1996 to 2010.

Swahili burial practice conforms to the general tradition throughout the Islamic World: burial with the body close to perpendicular to the geographic position of Mecca with the head and body resting on the right side, sometimes with a stone pedestal keeping the head in that position. The knees are flexed, arms rest on the side, and cross near the pelvis area or directly into the pelvis cavity. Other Muslim practices, including shrouding with no coffin and lack of grave goods, are consistent with the Islamic tradition in many world areas. In two cases, we found objects within the burial: a brass finger ring and a chicken femur bone found inside the skeletal mouth of a young child. The symbolic meaning or accidental occurrence of these objects is not understood.

At Mtwapa, one tomb contained at least three individuals. Otherwise, all burials are similar in their mortuary context. There appears to be no distinction between the burial practices associated with male/female or young/old. In several instances, the skeletons were preserved collapsed onto their ventral surface so that they appeared to be lying on their stomachs with the skull face down.

The state of preservation of the remains is varied, but generally, the skeletons from Manda are better preserved than those from Mtwapa, although all recovered materials were fragile and fragmentary. At Mtwapa, the bones, and especially the skull, underwent plastic deformation. So, although the bones are intact, the individual neurocranial bones' position was compressed and distorted.

*Manda*

*Lat: 2.103S Long: 41.021E*

Manda is one of the earliest dated towns on the Swahili coast. The total area of the site was over 40 acres out of which 18 were surrounded by a perimeter wall. The built-up area of the site presently covers about 18 acres. Manda's population at its height was perhaps 3500 people.

The late Neville Chittick surveyed and excavated the site for three seasons [1966, 1970 and 1978] and dated it the site to the 9^th^ century CE based on imported ceramics. His excavations at several key locales at the site revealed that Manda was one of the wealthiest towns in East Africa between 800 and 1600 CE [10]. Like many contemporary towns and cities along the Indian Ocean, the residents of Manda participated in Indian Ocean commerce that connected African, Asian, and Mediterranean mercantile communities [11]. Excavations carried out at the site in December 2012 yielded a wide range of artifacts including two Chinese Yongle coins of the 15^th^ century CE. Radiocarbon dates have now securely placed the earliest levels of the site at 600 CE making it one of the earliest known pre-Islamic urban settlements in sub-Saharan Africa [12].

Manda’s prosperity arose from its role in handling the export-import trade between the Northern East African interior and the Indian Ocean. The bulk exports passing through Manda included poles, cereals, rock crystal, leopard skins, hides and skins, ivory, honey, and beeswax. Bulk imports received at Manda include Indian beads and cloth, Arabian dates and jewelry, and Chinese and Islamic ceramics. The wealth accumulated from this commerce enabled Manda’s community to invest in more long-term permanent residential and public architecture and in vibrant cottage industries, including weaving, bead making, and iron production.

Low one by two-meter mounds characterize the graves at all four cemeteries identified in August 2009, and these all correspond to the mid-second millennium CE. Coral blocks outline some graves while others are enclosed in family graveyards. Few tombs survive. Those still standing are situated near the two mosques at the site. The first cemetery covers an estimated area of 40 square meters located on the northeastern part of the site in area AW. The second cemetery is adjacent to the northern mosque. The third cemetery is found near the town wall, outside the northwestern corner. The fourth cemetery is located outside the perimeter wall on the east side of the Peninsula. Based on previous excavations, the spatial organization of residential and cemetery locales are similarly organized and as expected, the more elite residences are located closest to the main congregational mosque. Elite individuals were buried near the mosque and non-elite individuals including immigrants were buried in cemeteries outside the perimeter wall.

The Manda site yielded the remains of 19 skeletons.

*Mwakwasinyi*

*Lat: 2.169S Long: 38.672E*

Mwakwasinyi is a locality in the Kasigau Hill of the Tsavo Region in inland southeast Kenya. The Mwakwasinyi are Taita people. The Taita are an ethnolinguistic group of diverse origins that reside in a coastal hinterland region of southeastern Kenya and speak Dawida and Saghala, Bantu languages with southern Cushitic loan words. The Taita inhabit the uplands and slopes of three hills, Dawida, Saghala, and Kasigau. The historical and oral traditions of the Taita suggest that they settled in the Tsavo region ca. 1000 years ago, absorbing Bisha pastoralists and Laa or Wawasi hunter-gatherers. Since at least the seventeenth century, they have also incorporated refugees and visitors from diverse areas of southeastern Kenya. Their present-day neighbors include the Akamba and Taveta agro-pastoralists, Somali and Orma pastoralists, and Waata foragers. The Taita have interacted with their neighbors through trade and intermarriage. They also have developed social networks called blood brotherhoods, which facilitate economic exchange and minimize inter-group conflict. They are traditionally an agropastoral people who grew millet, eleusine, and sorghum. They raised cattle and engaged in regional trade with the coastal communities - the Mijikenda and Swahili with whom they exchanged ivory, iron, grains, cattle, animal skins, beads, cloth, cowrie shells, and marine fish. They forged similar networks with the Taveta, Maasai, and Pare peoples of Usambara and Kilimanjaro in the interior. In addition, their patrilineal clans incorporated outsiders from neighboring communities through alliances and intermarriage over at least the last three hundred years. These interactions may have impacted their genealogical composition, which may be detectable through genetic studies.

According to Taita oral traditions, the first inhabitants of the Tsavo region were the Laa or Wawasi hunter-gatherers, and subsequent inhabitants included the Wambisha, who were pastoralists. They lived in autonomous villages along the mountain ridges with boundaries defined by streams and rivers. These villages were organized into large lineages. The lineages were not exogamous, so many of the women did not marry outsiders but instead married within their natal group. Within the large lineages, smaller segments saw the inheritance of cattle and land, for which there was sometimes matrilineal inheritance. Large lineages maintained a ritual focus around a shrine center with a semi-protected repository of the skulls of the dead, normally exhumed two years after burial.

Ecological differences and predictable climatic regimes around the mountains and the wider Tsavo region made inter-community interaction and exchange crucial. Trade was important for dealing with seasonal food shortages and surpluses. Village and regional market systems fostered intra and inter-group social networks, which involved exchanging agricultural and non-agricultural products with seasonal availability. Through these systems of exchange and interaction, the Taita built and maintained strong ties with their neighbors.

We obtained cranial samples representing 13 individuals from exhumed human remains kept since at least 1952 within a rockshelter cranial display in the village of Makwasinyi and associated with the Taita people. The rockshelter has been used since at least the 18th and 19th centuries as an above-ground tomb by a Taita extended family in Makwasinyi Village, Taita County, Kenya. The Taita peoples are the closest hinterland neighbors and trading partners of the Swahili peoples and so we hypothesized that comparison of the Swahili individuals to them might be informative about the African ancestry sources for Swahili people.

*Songo Mnara*

*Lat: 9.040S Long: 39.552E ~16 m asl, Kilwa District, Lindi Region, Tanzania*

The site of Songo Mnara is located on the northwest coast of Songo Mnara Island and is one of the best-preserved examples of a 15th-16th century CE Swahili stone town [13]. The site was extensively mapped in 1961 [14], and Chittick carried out test excavations in the 1960s (unreported). Further excavations were carried out by Pradines in 2004 [15]. Four seasons of archaeological research were carried out from 2009 to 2016 by co-authors S. Wynne-Jones and J. Fleisher, exploring the use of domestic and public space at the site under the Songo Mnara Urban Landscape Project [16, 17]. Based on local and imported ceramics and beads, the occupation dates from the late 14th to early 16th centuries, with some evidence of modest reoccupation in the 17th century [18].

For a relatively small Swahili stone town, Songo Mnara contains a large number of burials in three separate cemeteries (*Figure S1*). Two cemeteries are located at the margins of the site, one to the west and one to the east. The western cemetery is a walled graveyard associated with a mosque, set on a high bluff southwest of the town. The eastern cemetery is located just outside the eastern wall of the settlement. The main cemetery is located at the center of the site, surrounded by domestic structures on the north, south, and east, and open space to the west that extends to the entrance zone of the settlement. Most graves are marked with sandstone head and footstones, with a small number in the central cemetery marked with coral rag tombs. Tombs are walled enclosures that often surround head and footstones, sometimes built just at the ground surface and up to a meter or more. There is evidence of commemorative practice at some burial sites, with locally-minted copper coins and small water-worn quartzite pebbles apparently left on the grave sites [17].

Fieldwork in 2011 included excavations of four areas with burials (*Figure S1*); these excavations revealed human remains of 14 individuals (Table S1). Two trenches, SM024 and SM025, were excavated in the central cemetery. SM024 incorporated an area with head and foot stone markers, while SM025 was situated in a coral rag tomb enclosure. Trench SM026 was located within a walled graveyard on the eastern side of the central cemetery, just south of a small coral rag mosque; this trench also included head and foot stone markers. Trench SM027 was located in a cemetery outside the eastern wall of the settlement, an area marked by head and foot stone markers; the trench incorporated an area with a head and foot stone. Seven individuals from these burial excavations are included in this aDNA project: SK3, SK4, SK7, SK8, SK9, SK12, and SK13. One additional individual, represented by sample SF20151, is the mandible of an infant found in the mixed fills of trench SM020; no dentition was recovered.

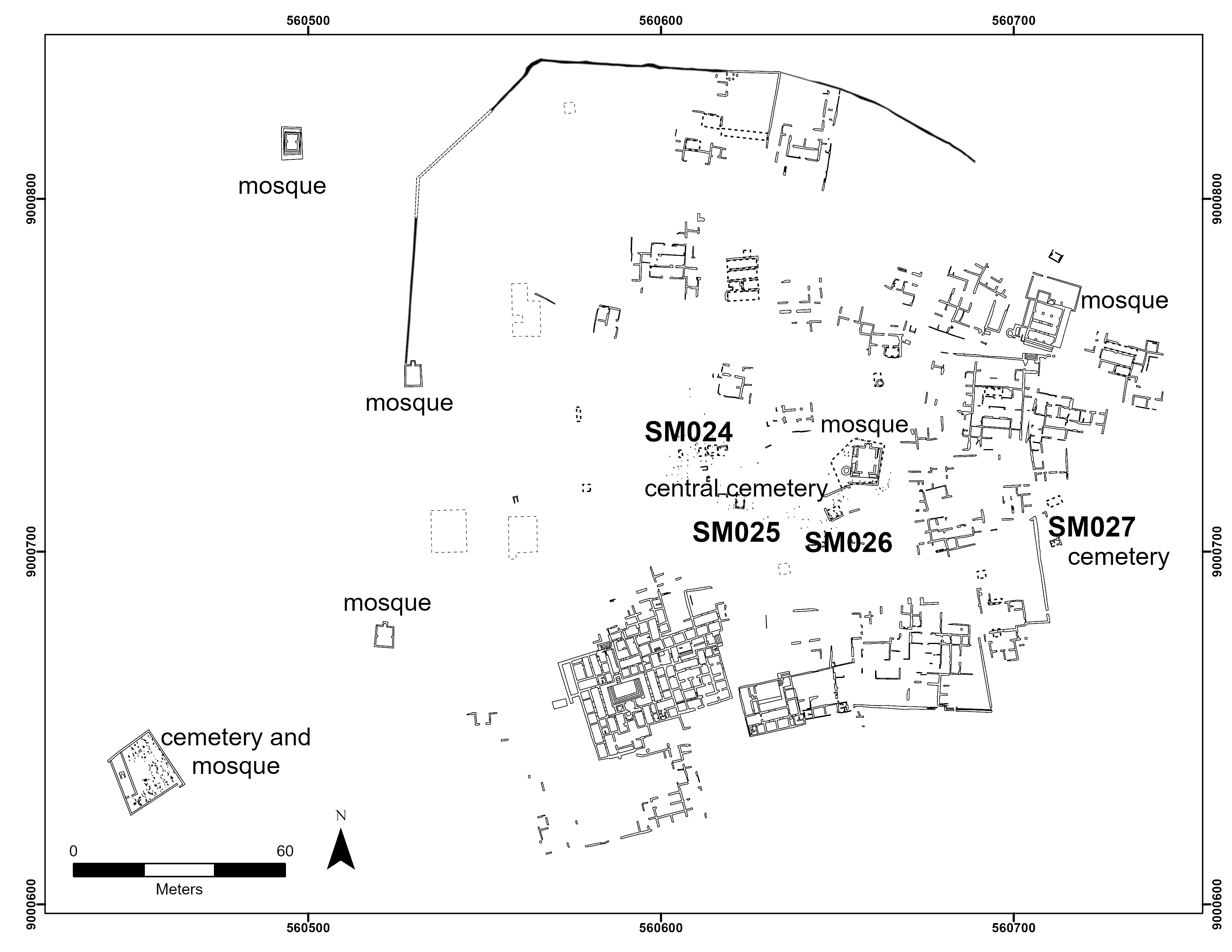

*Figure S1:* Site map of Songo Mnara with locations of excavations associated with human remains.

Radiocarbon dating of the individuals reported here provides evidence of the chronology of burial at Songo Mnara (Table S1). One burial dates to the 14th century CE (SK12); this individual was buried in the central part of the central cemetery and was stratified below three other burials interred sequentially in this walled tomb. Three of the excavated burials (SK7, SK9, SK13) date firmly to the first half of the 15th century CE; these burials were located in the far west of the central cemetery, the far east of the central cemetery (in the walled graveyard), and in the cemetery east of the town wall. This suggests that these main cemeteries were in use by the beginning of the 15th century and that internment was occurring across the site. Finally, three individuals date from the 16th or 17th centuries CE. Two of these (SK4, SK8) were located side by side, and the aDNA data presented in this study suggests they were 2nd or 3rd degree relatives. One additional individual, an infant represented only by a mandible (without dentition), was found in the mixed fill of excavations in House 47 along the site’s northern wall and dates to the 16th and 17th centuries. Together, these individuals support other chronological indicators that a few structures in the northern part of the site were inhabited into the 16th or 17th centuries.

| **SK#** | **Trench#** | **Burial marker type** | **Position** | **Skeletal completeness and bone preservation** | **Sample taken for ancient DNA analysis** | **Estimated age at death** | **Morphologically determined sex** | **Genetically determined sex** |
| --- | --- | --- | --- | --- | --- | --- | --- | --- |
| SK1 | SM025 | head/foot stones | n/a | metacarpal only | not sampled | adult |  |  |
| SK2 | SM025 | rectangular tomb (northern section) | semi-flexed on right side |  | not sampled | neonate, 6-9 months |  |  |
| SK3 | SM025 | rectangular tomb (southern section) | extended on right side | 85% complete, good | metacarpal | adult | M |  |
| SK4 | SM024 | head/foot stones | semi-flexed on right side | nearly complete, excellent | right metatarsal | adult | M | F |
| SK5 | SM025 | rectangular tomb (southern section) | semi-flexed on right side |  | not sampled | neonate |  |  |
| SK6 | SM026 | head/foot stones | semi-flexed on right side |  | not sampled | neonate, 6-9 mo |  |  |
| SK7 | SM024 | head/foot stones |  | just metatarsals | second left metatarsal | adult |  | M |
| SK8 | SM024 | head/foot stones | extended on right side | nearly complete, excellent | right metacarpal | adult | M | M |
| SK9 | SM026 | head/foot stones | extended on right side | 80% complete, fair | metacarpal | adult, 18-23 | F | F |
| SK10 | SM027 | head/foot stones | semi-flexed on right side |  | not sampled | child, 5-6 years |  |  |
| SK11 | SM026 | head/foot stones | semi-flexed on right side |  | not sampled | child, 6-12 mo |  |  |
| SK12 | SM025 | rectangular tomb (southern section) | semi-flexed on right side | 70% complete, very poor | right and left metatarsal | adult | M | M |
| SK13 | SM027 | head/foot stones | extended, on right side, partially facing downward | nearly complete, excellent | metacarpal | adult | M | F |
| SK14 | SM026 | head/foot stones | extended, on right side |  | not sampled | child |  |  |
| SF  20151 | SM020 | in fill | N/A | isolated mandible | mandible | neonate |  | M |

Table S1: Skeletons excavated at Songo Mnara in 2011. Individuals with genetically determined sexes have genetic data.

Excavations of human remains were carried out after consultation with the local community, represented by the Village Ruins Committee [19]. We approached the VRC with a document that described research questions that were possible to address through the excavation of human remains, including issues of chronology, diet, paleopathology, and aDNA. The community's primary concern was that the excavated individuals not be removed from the site. We, therefore, devised a protocol in which skeletons were exposed, excavated, removed, documented, and sampled in one day and reinterred the following day. At the end of the field season, the local imam performed a blessing over the burials. Analysis of the skeletal remains was supervised in the field by Prof. Kate Robson Brown (University of Bristol) and Dr. Francesca Migliaccio (now University of Bath).

*Kilwa Kisiwani*

*Lat: 8.959S Long: 39.497E 14 m asl, Kilwa District, Lindi Region, Tanzania*

Kilwa Kisiwani, on Kilwa Island, is designated a UNESCO World Heritage Site jointly with Songo Mnara, immediately to its south. Kilwa is renowned for coral architecture, including the 11th (enlarged 14th) century Great Mosque that still stands today, and Husuni Kubwa, a grand 13th-14th century palace, both testifying to Kilwa’s role as a major center of medieval and early modern Indian Ocean trade (for a general overview, see [20]). Occupied since the 9th century [21], Kilwa attained its peak of power in the 13th-15th centuries and remains occupied to this day. Elite merchants controlled trade routes that moved gold, copper, ivory, and other goods from the interior to the coast at Kilwa, and the port minted its own copper currency for centuries. Kilwa’s trading networks extended along the coast to Sofala (Mozambique), and as far inland as Great Zimbabwe (Zimbabwe), and most notably across the Indian Ocean interaction sphere, with connections as distant as Arabia and China. Portuguese occupation in 1505 and control of this and other ports in the 16th century precipitated Kilwa’s loss of status.

Excavations at Kilwa in the 1950s-60s by Chittick [22] produced much of what we know of the site’s archaeology, but his interpretations were shaped through a colonialist lens wherein non-African actors were seen as responsible for the city-state’s achievements, even though he recognized that African settlements marked its earliest stratigraphic levels. Subsequent research has shown Kilwa and other coastal city-states to be fundamentally African in origin, with important external connections including foreign merchants (reviewed by Wynne-Jones & Fleisher 2015 [23]; see also main text).

Chittick’s chronology, which was based mainly on ceramics, was recently reevaluated through a limited re-excavation of the site [24]. Unfortunately, linking the burials in this study to site chronology is challenging, due to the lack of contextual information for those burials. None of the remains in this study can be unambiguously connected to the skeletal elements listed in Appendix II of Chittick (1974), which reports incomplete burials mixed with faunal remains, in the southern courtyard of the Great Mosque.

Two boxes of human remains from Kilwa are presently curated at the British Institute in Eastern Africa (BIEA) in Nairobi. An inventory of these remains, which represent several individuals, was made by forensic anthropologist Shari Eppel in an unpublished 2016 report for the BIEA. We found the remains to be precisely as Eppel described them. As Eppel notes in her report, the Kilwa remains appear to have been repackaged in plastic bags at some point prior to 2016, and some contextual information may thus have been lost.

Kilwa Box 1 contains remains with bags reading “SEB ZLL SS” and “SEB ZLL SS 9/9a/7b,” thus connecting this to the Period Ia levels of trench ZLL, located near the center of the site and south of the Great Mosque and Great House. A disturbed burial was found in this trench, and its excavation is documented in photographic archives [25]. A new neighboring trench confirms a chronology for the burial of c. 800-1000 CE [24]. Eppel’s report notes multiple bags of fragmented remains of at least one juvenile, represented by the left and right maxillae with partial dentition and fragments of the mandible, long bones, and phalanges, and one older individual (only represented by a cranial vault fragment). We collected two samples potentially belonging to the same juvenile: a deciduous right upper central incisor (KS.01.01), and a cranial fragment taken as backup for radiocarbon dating (KS.01.02). The incisor was selected because it was isolated and attached to the reconstructed maxilla with putty because its antimere was present. We sampled only KS.01.01 for aDNA and this sample failed to produce working data; we therefore returned the remainder of this sample, and the intact KS.01.02 sample, and no radiocarbon dates were generated per our protocol.

Kilwa Box 2 contains more abundant human remains, but with far less context information. All that is known is from a single bag on which “Bones from BB” is written with marker, and Eppel notes additional notation of “KM skeleton”. “BB” could refer to multiple spaces within structures at Kilwa, according to Chittick’s (1974) nomenclature system [22]. A ‘lobby’ space in Husuni Kubwa was labeled BB, but the excavation notes do not note any human remains, or indeed sub-floor deposits. Perhaps more likely are the ‘western buildings’ of the Makutani complex, where exploratory excavations were labeled BB; these explorations are not documented in detail, but the complex is thought to be 15th century [22].

At least two individuals are identified in Eppel’s inventory in Box 2 based on size, repeating elements, and skeletal development: one adult, and one smaller subadult who died before age 18, based on the presence of a scapula fragment of glenoid fossa still in the process of fusing. Both individuals are fragmentary and incomplete, with limited osteobiographic details. We attempted to collect a sample from each of the commingled individuals. The first sample (KS.02.01) was a fragment of left temporal preserving the petrous portion labeled “part of larger cranial fragment,” which we interpreted as associated with the larger, adult individual. The other sample, a lower canine (KS.03.01), showed some wear but was plausibly from the younger individual. Both produced readable aDNA, although not upon initial screening, and do in fact represent separate individuals. Because our sampling protocol sought to minimize destruction and expedite sample return, we did not radiocarbon date these individuals, because at the time of return, it was thought that there was no working aDNA data. By the time the lab had achieved better results, the samples were already returned to Kenya. This means that the two individuals with working aDNA from Kilwa are undated, and future research should endeavor to obtain direct dates.

*Lindi*

*Lindi town approximate location: Lat: 9.99S Long: 39.71E, Kilwa District, Lindi Region, Tanzania*

Lindi town is located on the coast of southernmost Tanzania, c. 90 km from the Mozambican border. The broader Lindi region, and Mtwara region to its south, have numerous archaeological sites spanning the Middle Stone Age [26-28] to the Later Stone Age and Early Iron Age (e.g., [29-31]). During the medieval era, Lindi would have been situated along key maritime routes and at times fallen within Kilwa’s sphere of influence [32-34]. In the 19th century, it became a terminus of slave-raiding caravans stretching from Lake Malawi to the coast and played a key role in resistance to colonial rule, in particular during the early 20th century Maji Maji rebellion (e.g., [35]).

During construction of a hotel near the beach at Lindi town in 2013 (exact location unknown), a burial was uncovered (and severely damaged) by a bulldozer. The remains, accompanied by sparse material culture, were brought to the National Museum of Tanzania (NMT) in Dar es Salaam, where they are still curated. A handwritten note dated to December 12, 2013 records the provenience information as coming from 2.5 meters below an old house floor, and indicates that the donated skeleton was accompanied by coconut fibers, five potsherds, and a piece of rubble originating in the house floor.

Given the absence of contextual information and associated archaeology for the remains, taken together with the archaeological and historic importance of the broader region, curators at NMT wanted to know more about this individual and requested sampling for aDNA and radiocarbon dating. We also conducted a brief skeletal inventory and osteobiographic assessment.

The remains are consistent with a single individual. The skull is mostly complete but broken in two roughly along the coronal plane. Parts of both the cranium and mandible have been reconstructed using glue. Postcranial remains include lateral parts of both scapulae, a complete right humerus, most of the right radius and ulna, the proximal left ulna, a left lunate. most of the left innominate, the right ilium, the sacrum, and parts of both femora. Part of the spine including the atlas and axis, all twelve thoracic vertebrae, and one lumbar vertebra are also present. The right ilium was previously glued to the sacrum. Sexually dimorphic traits of the pelvis and skull are all consistent with male sex. Age at death was most likely in middle adulthood. The left pubic symphysis shows no billowing but some transverse organization and microporosity, consistent with Suchey-Brooks phase 2 or 3, suggesting the individual was in their 20s or 30s. One rib shows lipping on the costal facet, as well as a shallow, u-shaped sternal end also consistent with middle age. The dentition shows advanced signs of dental disease focused on the cheek teeth, including antemortem tooth loss of several molars, two necrotic root stumps, and caries on the *in situ* right mandibular third molar. By contrast, the anterior teeth are present and worn but not carious. Bone preservation was excellent in the sense that there was little post-depositional diagenesis or damage from the burial environment, but the skeleton had evidently been severely damaged by the bulldozer, with numerous fresh breaks and many skeletal elements missing.

Two samples were chosen from this individual: the right upper central incisor (LS1.01), and the left petrous portion of temporal bone (LS1.02). Only LS1.02 was sampled, producing working aDNA data (individual number I14001) and a direct radiocarbon date of 305±15 uncalibrated years before present (PSUAMS-5718). LS1.01 was returned unsampled.

### PCA setup

We aimed to build a PCA (Figure 1B, Figure 3B, and Extended Data Figure 3) [36, 37] that would allow differentiation along at least 5 ancestry poles of interest: East African and Bantu-associated, European, Near Eastern, South Asian, and East Asian. The goal of distinguishing these ancestry sources is that it would allow us to differentiate different types of plausibly Eurasian ancestry.

We carried out PCA using the following set of 1286 present-day individuals from 73 African and Eurasian populations genotyped on the Affymetrix Human Origins single nucleotide polymorphism (SNP) array (hereafter the “Human Origins (HO) dataset”). The population labels of the individuals in the dataset are listed with sample size in parentheses in what follows: Armenian (10), Balochi (20), BantuKenya (6), Belarusian (10), Brahui (21), Bulgarian (10), Burusho (23), Cameroon_Mbo (21), Croatian (10), Cypriot (8), Dinka (7), Druze (39), Egyptian (18), English (10), Ezid (8), Finnish (7), French (61), Georgian (23), Hungarian (20), IBS_CanaryIslands (2), India_Non_Zoroastrian_Hindu (11), India_Zoroastrian (13), Iranian (38), Iranian_Bandari (8), Iran_Non_Zoroastrian (17), Iran_Zoroastrian (24), Italian_Sardinian (2), Jain (3), Jordanian (9), Juang (26), Kalash (17), Kamboj (31), Khomani (11), Kikuyu (4), Kumyk (8), Kurd (8), Lebanese (8), Lebanese_Christian (9), Lebanese_Muslim (11), Lezgin (9), Libyan (5), Lithuanian (10), Luhya (8), Luo (8), Makrani (20), Malawi_Chewa (11), Malawi_Ngoni (4), Malawi_Tumbuka (10), Malawi_Yao (9), Maltese (8), Masai (12), Mende (8), Muslim_Jat (4), Ossetian (14), Pathan (17), Romanian (10), Russian (71), Sandawe (22), Sardinian (25), Saudi (8), Shia_Iranian_Hyderabad (4), Sicilian (11), Sikh_Jatt (41), Sindhi_Pakistan (14), Spanish (172), Tajik (31), Tamta (1), Turkish (50), Turkmen (6), Ulladan (17), Uzbek (27), Yemeni (6), and Yoruba (21) [21, 38-47]. Eigenvector 1 separates individuals from sub-Saharan Africa and other individuals. Eigenvectors 2 and 3 form a plane separating European-associated, Near Eastern-associated, South Asian-associated, and East Asian-associated poles. We chose populations for constructing axes to ensure that the South Asian – Near Eastern cline becomes the focus of the variation in eigenvectors 2 and 3. We then projected all other individuals.

On the PCA in Figure 1B, the people from medieval burials at Manda and Mtwapa in Kenya together form a cline with one pole pointing toward the historic Makwasinyi individuals and the other pole falling somewhere along a Eurasian gradient from the Near East to South Asia. The Mtwapa individuals vary in their degree of genetic affinity to these Eurasians, raising the possibility that we are sampling individuals from an ongoing admixture event. The inland Makwasinyi group falls along a gradient that has at one extreme ancient and present-day Bantu-speaking populations and their West African genetic relatives, plausibly reflecting the spread of peoples speaking Bantu languages from West/Central Africa after 2000 BCE [48, 49]; we call this “Bantu-associated ancestry” in what follows, without claiming that the people who carried this ancestry spoke Bantu languages, as genetics cannot reveal what languages people spoke. At the other extreme of the cline were ancient African pastoralists (based on associated archaeological evidence for livestock herding) who lived in Kenya and Tanzania primarily between 1000 BCE - 500 CE (we call this “Pastoral Neolithic (PN)-associated ancestry” in what follows). The Kilwa individual and at least one Songo Mnara individual (I19550) also appear near the Mtwapa-Manda cline, but with less proximity to Eurasian groups. The rest of the Songo Mnara individuals have heterogeneous genetic affiliations, with some individuals appearing to have little, if any, relatedness to Eurasians. The Tanzanian Lindi individual appears to have nearly entirely Bantu-associated ancestry with little or no Eurasian affinity.

### *qpAdm* modeling

#### Modeling medieval coastal groups from Mtwapa, Manda, Songo Mnara, Kilwa, and Lindi as well as the inland Makwasinyi group

Unless otherwise specified, our modeling of ancient population uses the “1240k dataset”, a term we use to identify a set of mostly ancient and historic individuals genotyped for the full set of sites targeted by the 1240k in-solution enrichment reagent. Because the individuals in the “Human Origins dataset” are genotyped at only about half the positions targeted by the 1240k reagent, the present-day individuals genotyped on the Affymetrix Human Origins dataset are not included in the 1240k dataset. Whenever we coanalyze the two datasets, we do so by pulling down the 1240K dataset on the HO SNP set. We additionally coanalyze the 1240K dataset with shotgun sequenced individuals (that have ‘.DG’ or ‘.SG’ suffixes) on the 1240K SNP set.

*Makwasinyi*

Using the *qpAdm* framework [39], the Makwasinyi group can be modeled ($P=0.55$) as an admixture of a 21.3±1.2% contribution from Pastoral Neolithic (PN)-associated ancestry, represented in our modeling by Kenya_PastoralN individuals (KPN) [48], and 78.7±1.2% Bantu-associated ancestry most closely matching Bantu-speaking source populations in eastern and southern Africa, represented in our modeling by Tanzania_Pemba_600BP_published, a 600BP individual from Makangale Cave on Pemba Island [21]. But other PN-associated groups and Bantu-associated individuals, including the individual buried at Lindi, produced working models as well (*Table S2* shows some working models).

| **Pastoral group** | **Bantu-associated individual** | **P value** | **Proportion of Bantu-associated ancestry** | **Coefficient of pastoral ancestry** |
| --- | --- | --- | --- | --- |
| Kenya_PastoralN | Tanzania_Pemba_600BP_published | 0.27 | 0.79±0.01 | 0.21±0.01 |
| Kenya_PastoralN_Elmenteitan | Tanzania_Pemba_600BP_published | 0.22 | 0.79±0.01 | 0.21±0.01 |
| Kenya_LukenyaHill_PastoralN | Tanzania_Pemba_600BP_published | 0.77 | 0.78±0.01 | 0.22±0.01 |
| Kenya_HyraxHill_PastoralN | Tanzania_Pemba_600BP_published | 0.18 | 0.79±0.01 | 0.21±0.01 |
| Tanzania_PN | Tanzania_Pemba_600BP_published | 0.12 | 0.8±0.01 | 0.2±0.01 |
| Kenya_PastoralN | Tanzania_Lindi_Swahili | 0.12 | 0.8±0.01 | 0.2±0.01 |
| Kenya_PastoralN_Elmenteitan | Tanzania_Lindi_Swahili | 0.17 | 0.8±0.01 | 0.2±0.01 |
| Kenya_LukenyaHill_PastoralN | Tanzania_Lindi_Swahili | 0.54 | 0.8±0.01 | 0.2±0.01 |
| Kenya_HyraxHill_PastoralN | Tanzania_Lindi_Swahili | 0.87 | 0.8±0.01 | 0.2±0.01 |
| Tanzania_PN | Tanzania_Lindi_Swahili | 0.16 | 0.81±0.01 | 0.19±0.01 |

Table S2: Successful qpAdm 2-way models with Bantu-associated and PN-associated sources.

Individual proportions of admixture can be seen in Table S*3*. Reference or “right” group populations in all *qpAdm* runs for modeling the Makwasinyi group are Altaian.DG, Iranian.DG, Jew_Yemenite.DG, Onge.DG, Brahmin.DG, Spanish.DG, Han.DG, Ami.DG, Kalash.DG, Dusun.DG, Igorot.DG, Yoruba.DG, Dinka.DG, Mozabite.DG, Mende.DG, Hadza_1.DG, Mbuti.DG, Mursi.DG, Luo.DG, South_Africa_1900BP.SG, Druze.DG, and Sandawe.DG.

| **Makwasinyi individual** | **P value** | **Proportion of Bantu-associated ancestry** | **Coefficient of pastoral group** |
| --- | --- | --- | --- |
| I14781 | 0.48569 | 0.89±0.02 | 0.12±0.02 |
| I13872 | 0.53364 | 0.77±0.02 | 0.23±0.02 |
| I13875 | 0.37509 | 0.79±0.02 | 0.22±0.02 |
| I13874 | 0.38109 | 0.81±0.02 | 0.19±0.02 |
| I13871 | 0.69126 | 0.79±0.02 | 0.21±0.02 |
| I17404 | 0.01377 | 0.76±0.03 | 0.24±0.03 |
| I17402 | 0.37174 | 0.77±0.02 | 0.23±0.02 |
| I17401 | 0.31615 | 0.79±0.02 | 0.21±0.02 |
| I17405 | 0.13964 | 0.81±0.03 | 0.19±0.03 |
| I17403_d | 0.35854 | 0.86±0.04 | 0.14±0.04 |

Table S3: qpAdm determined proportions of ancestry for each Makwasinyi individual. The Bantu-associated proxy source is Tanzania_Pemba_600BP_published and the PN-associated source is Kenya_PastoralN.

We sought to challenge the 2-way admixture model for the Makwasinyi individuals by cycling through additional ancient populations among the right-hand outgroups in *qpAdm*. The individuals that we cycled are those from a 1240K dataset of African individuals that have at least 200,000 SNPs on the 1240K SNP set*.* No population (except Kenya_PastoralN_published, which would have issues of bi-directional gene-flow) completely breaks the model for Makwasinyi, and all P values are above 0.1.

*Mtwapa and Manda*

In the PCA, the inland Makwasinyi group appears be an appropriate proximal surrogate sub-Saharan African-associated source for the Mtwapa and Manda sampled individuals. We initially tried to form a 2-way model between Makwasinyi and a Eurasian population from the Human Origins dataset (with Tatar_Siberian, Onge, Juang, Palliyar, Italian_Central, Cameroon_Mbo, Han, and Ami as the initial “right” group reference populations). Out of all Eurasian HO populations tested, only Shia_Iranian_Hyderabad, an Iranian-Indian admixed population, provided a working model (p>0.1).

This led us to try a 3-way model for the Mtwapa and Manda groups (with Tatar_Siberian, Onge, Juang, Palliyar, Italian_Central, Cameroon_Mbo, Han, Ami, Turkish, Lebanese, Jew_Yemenite, Kalash, Murut, and Kankanaey as the final “right” set of reference populations), using the Makwasinyi population as an African proxy source and the Iranian population as a West Eurasian proxy source, and cycling through all other Eurasian populations in the Human Origins dataset. We find working models with South Asian populations, with the best fits for Indian populations (3-way model of Makwasinyi, Iranian, and the Indian population of Sahariya_MP has $P=0.23$ or the Indian population of Ulladan has $P=0.13$). We tested an Indian population (Sahariya_MP) against a population from Bangladesh (BEB) that also produced a working model and found that the Indian population produced the more successful model (Table S4).

| **Left group source population** | **Right group reference population** | **P value without reference population** | **P value with reference population added** | **P value change** |
| --- | --- | --- | --- | --- |
| Sahariya_MP | BEB | 0.228 | 0.189 | 17% |
| BEB | Sahariya_MP | 0.155 | 0.077 | 50% |

Table S4: Comparing South Asian populations in a 3-way model for the ancestry of the Mtwapa group.

Some of the best fitting Indian populations include ancestry that is maximized in East Asia (Figure S2), which led us to wonder whether this implies a possible East Asian component of ancestry in addition to or in place of the Indian component in the Mtwapa group. We tested the robustness of the Indian-associated component of ancestry by trying to break the 3-way Makwasinyi-Iranian-Indian model with more eastern Asian populations from China, Indonesia, Myanmar, Thailand, Cambodia, Korea, Vietnam, Singapore, and the Philippines as reference populations. Since Indian populations themselves include varying amounts of Iranian-associated ancestry, the closest Indian population is hard to determine as it is masked by the Iranian source population. As a surrogate Indian source, we choose Ulladan, a population which resides near the southwestern coast of India and has little of the East Asian ancestry that some other Indian populations have (determined via ADMIXTURE in Figure S2). The East Asian sources tested are not able to completely break the Indian model, whereas the Indian population breaks a model with the East Asian populations (Table S5). We also try adding the East Asian population in addition to the Indian population, but the P value is not necessarily indicative of a successful model (Table S5). Since the dimensions of left and right populations change, we cannot compare the P value when both populations are on the left with the P value when one of the populations is on the left and one on the right. We conclude that there is a significant component of ancestry in Mtwapa that stems from the Indian subcontinent, but more East Asian ancestry is not certain. In subsequent modeling, we represent the Indian component by Ulladan or Sahariya_MP, although there is no evidence that these groups rather than other Indian groups are most closely related to the correct source.

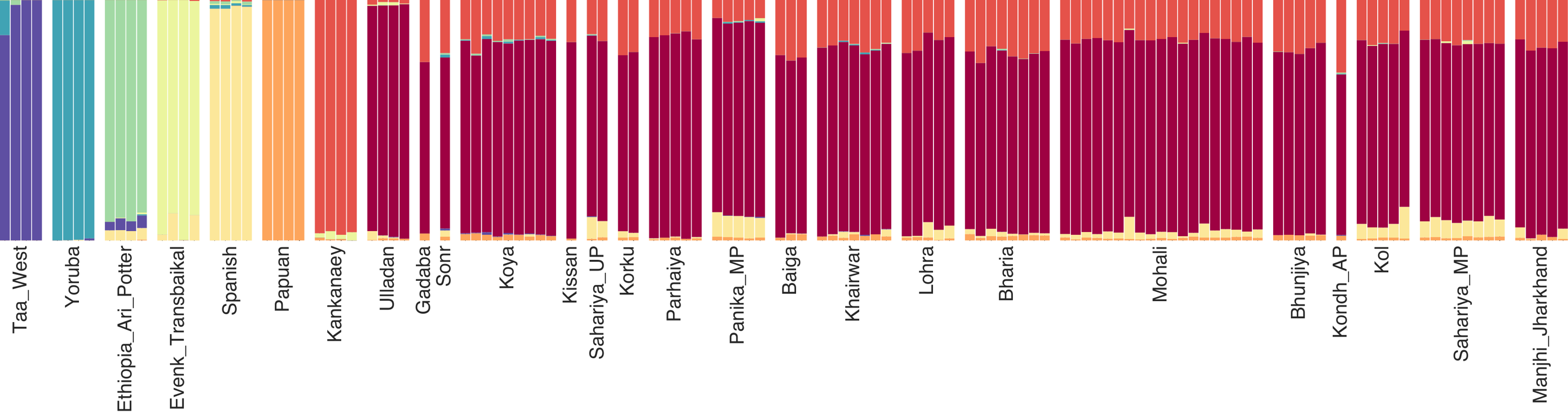

Figure S2: Best-fitting Indian populations have ancestry that is maximized in East Asian populations (ancestry represented by the color maximized by Kankanaey). Ulladan contains little, if any, of this East Asian reference ancestry.

| East Asian population | P value for East Asian population on left, Ulladan on right | P value for East Asian population on right, Ulladan on left | P value for both the East Asian population and Ulladan on the left |
| --- | --- | --- | --- |
| Dong | 0.000000 | 0.037832 | 0.001016 |
| Tibetan | 0.000000 | 0.036751 | 0.000479 |
| Zhuang | 0.000000 | 0.028833 | 0.000377 |
| Mulam | 0.000000 | 0.039024 | 0.000791 |
| Semende | 0.000000 | 0.085224 | 0.003609 |
| Barito | 0.000000 | 0.063535 | 0.005102 |
| Lebbo | 0.000000 | 0.109238 | 0.006306 |
| Thai | 0.000000 | 0.054518 | 0.000934 |
| CoLao | 0.000000 | 0.079455 | 0.000954 |
| Kinh | 0.000000 | 0.046736 | 0.001061 |
| HaNhi | 0.000000 | 0.080331 | 0.003677 |
| Hmong | 0.000000 | 0.078724 | 0.001535 |
| Vietnamese | 0.000000 | 0.043227 | 0.000314 |
| PhuLa | 0.000000 | 0.071494 | 0.001726 |
| Korean | 0.000000 | 0.043593 | 0.000522 |
| Tagalog | 0.000000 | 0.048293 | 0.011007 |
| Ilocano | 0.000000 | 0.106148 | 0.005554 |
| Mamanwa | 0.000000 | 0.103052 | 0.002416 |
| Cambodian | 0.000000 | 0.070054 | 0.002386 |
| Malay | 0.000000 | 0.085806 | 0.008148 |
| Burmese | 0.000000 | 0.086605 | 0.000681 |
| Atayal | 0.000000 | 0.047053 | 0.004201 |
| Bajo | 0.000000 | 0.044197 | 0.004304 |
| Ede | 0.000000 | 0.090621 | 0.003866 |

Table S5: Perturbing the Makwasinyi-Iranian-Ulladan model for Mtwapa with East Asian populations.

We next remove the Iranian population as a proxy West Eurasian source. Then, along with Makwasinyi and Sahariya_MP as sources, we iterate through all other West Eurasian HO populations. We obtain feasible, working models only with Persian-associated populations (Table S*6*). We thus continue to use Iranian as a surrogate Persian-associated source.

| **Eurasian Population** | **P value** | **Makwasinyi coefficient** | **Eurasian population coefficient** | **Sahariya_MP coefficient** |
| --- | --- | --- | --- | --- |
| Iranian | 0.227795 | 0.58±0.01 | 0.36±0.01 | 0.06±0.01 |
| Iran_Zoroastrian | 0.094571 | 0.58±0.01 | 0.35±0.01 | 0.07±0.01 |
| Iran_Non_Zoroastrian | 0.070514 | 0.58±0.01 | 0.36±0.01 | 0.06±0.01 |

Table S6: Feasible models for Mtwapa showing that the West Eurasian source is Persian-associated. Only 3 populations have $P>0.05$, and only Iranian has $P>0.1$.

We then test Makwasinyi as a proxy African source first by cycling through a set of 1240K ancient African populations or individuals that have at least 200,000 SNPs when pulled down on the HO SNP set and that do not have the qualifier (‘_all’ or ‘_contam’) (1240K coanalyzed African dataset, first column of Table S7). Only four populations (highlighted in gray in Table S7) produce feasible and successful models. Kenya_Kakapel_LIA falls on a PCA between Bantu-associated and PN-associated poles [50] similar to the Makwasinyi population (Figure 1B and Extended Data Figure 3). Kenya_LSA and Kenya_Kakapel_LSA_Kansyore have forager ancestry. Together, this likely indicates that the African source population or populations are a mixture of Bantu-associated, PN-associated, and forager-associated ancestries. The Makwasinyi group has the highest SNP count and the most individuals, making its fit more impressive. We continue our analysis using the Makwasinyi group as the African proxy source.

| **Ancient African 1240K population** | **P value** | **POP** | **Iranian** | **Sahariya_MP** |
| --- | --- | --- | --- | --- |
| Kenya_Manda_Swahili | 0.75 | 1.66±0.05 | -0.61±0.05 | -0.05±0.03 |
| Kenya_LSA | 0.44 | 0.63±0.01 | 0.37±0.01 | 0.01±0.01 |
| Kenya_Kakapel_LSA_Kansyore | 0.31 | 0.61±0.01 | 0.38±0.01 | 0.01±0.01 |
| Kenya_Makwasinyi_Swahili | 0.23 | 0.58±0.01 | 0.36±0.01 | 0.06±0.01 |
| Kenya_Kakapel_LIA | 0.21 | 0.56±0.01 | 0.39±0.01 | 0.05±0.01 |
| Tanzania_Lindi_Swahili | 0.08 | 0.52±0.01 | 0.42±0.01 | 0.06±0.01 |
| Tanzania_Kilwa_Swahili | 0.05 | 0.69±0.01 | 0.24±0.01 | 0.06±0.01 |
| Tanzania_SongoMnara_Swahili | 0.02 | 0.68±0.01 | 0.27±0.01 | 0.05±0.01 |
| Botswana_Xaro_EIA | 0.00 | 0.55±0.01 | 0.39±0.01 | 0.07±0.01 |
| Tanzania_Pemba_600BP_published | 0.00 | 0.52±0.01 | 0.42±0.01 | 0.07±0.01 |
| Congo_Kindoki_Protohistoric | 0.00 | 0.51±0.01 | 0.43±0.01 | 0.06±0.01 |
| Cameroon_SMA_published | 0.00 | 0.54±0.01 | 0.42±0.01 | 0.05±0.01 |
| Egypt_ThirdIntermediatePeriod | 0.00 | 7.19±0.75 | -7.34±0.85 | 1.14±0.14 |
| Kenya_EarlyPastoralN | 0.00 | 1.1±0.02 | -0.2±0.02 | 0.1±0.01 |
| Kenya_Historic_2 | 0.00 | 0.73±0.01 | 0.21±0.01 | 0.06±0.01 |
| Kenya_HyraxHill_PastoralN | 0.00 | 1.01±0.02 | -0.1±0.03 | 0.09±0.02 |
| Kenya_IA_Pastoral | 0.00 | 0.73±0.01 | 0.21±0.01 | 0.06±0.01 |
| Kenya_LukenyaHill_PastoralN | 0.00 | 0.92±0.02 | -0.01±0.02 | 0.1±0.02 |
| Kenya_MoloCave_PastoralN | 0.00 | 0.87±0.01 | 0.06±0.02 | 0.07±0.01 |
| Kenya_PastoralN | 0.00 | 0.95±0.01 | -0.04±0.01 | 0.09±0.01 |
| Kenya_PastoralN_Elmenteitan | 0.00 | 0.97±0.01 | -0.05±0.01 | 0.09±0.01 |
| Kenya_PastoralN_published | 0.00 | 0.94±0.01 | -0.03±0.02 | 0.09±0.01 |
| Morocco_Iberomaurusian | 0.00 | 1.32±0.02 | -0.37±0.03 | 0.04±0.02 |
| Tanzania_Luxmanda_3000BP | 0.00 | 0.95±0.02 | -0.03±0.02 | 0.08±0.02 |
| Tanzania_PN | 0.00 | 0.98±0.01 | -0.08±0.02 | 0.11±0.01 |
| Tanzania_PN_IA_o | 0.00 | 0.99±0.02 | -0.08±0.02 | 0.1±0.01 |

Table S7: Testing African populations for use as the surrogate African source in the Mtwapa qpAdm model.

We then put Makwasinyi on the left and each of the three other model-producing populations on the right in turn. We also tested the opposite scenario of Makwasinyi on the right and each of the other model-producing populations on the left in turn. Each of these runs reduced the P value by approximately one order of magnitude without completely breaking the model. There was no significant preference for any of the populations. The Makwasinyi population is likely a close proxy but has inexact proportions of Bantu-associated, PN-associated, and forager-associated ancestry components. It is important to note that PN-associated populations or Bantu-associated populations alone do not produce working models.

We cycled through 153 African populations from the HO dataset and added each one in turn to the right reference set. Many of the populations, particularly those with all types of forager ancestry, degraded the model generally by one two orders of magnitude. But none of these HO African population provides a working alternative model (all with $P<0.0001$). We tried adding an African HO population along with Makwasinyi to the left, but this produced no working models and caused extreme uncertainty in the coefficients of the African populations. Given this, the Makwasinyi population is the best African proxy source, albeit likely without the correct or enough forager-associated ancestry.

A potential caveat to these findings about uniquely good fits for Makwasinyi is that they may in part reflect technical issues. The Makwasinyi, Manda, and Mtwapa data were generated using the 1240k in-solution enrichment process, so that there are no systematic differences in data type between the source and target populations. In contrast, when a population genotyped on the Affymetrix Human Origins array is used as a source with Makwasinyi as a reference population, technical biases in the 1240k data generation process may cause artifactual attraction between Manda/Mtwapa on the one hand, and Makwasinyi on the other, which could cause the model to fail even if in truth the population used as a source is just as valid a source as Makwasinyi. Regardless of whether this technical issue is contributing to our result of uniquely good fits for Makwasinyi, our analyses are sufficient to demonstrate that Makwasinyi is at least as good a source population as any other African population we tested, which is the most important observation.

The same Makwasinyi-Persian-Indian 3-way model fits the Manda population ($P= 0.54$ for Ulladan and $P= 0.61 for Sahariya\_MP$). We cycled 745 Eurasian and African present-day groups from the Human Origins genotyping dataset (Eurasian and African populations from Online Table 5 | coanalyzed_modern_samples that did not have ‘_all’ or ‘_contam’ qualifiers and that were not relatives) as reference populations, and only four degraded the Makwasinyi-Iranian-Sahariya_MP models without completely breaking them $0.01<P<0.1$: Brahmin_Nepal, India_Non_Zoroastrian_Hindu, Himba, and Mbukushu. Brahmin_Nepal and India_Non_Zoroastrian_Hindu likely indicate that the Indian surrogate is not the exact source. Himba and Mbukushu degrade the model to $0.09<P<0.1$, and could indicate an inexact African surrogate population. The multiple comparisons problem means that the more hypotheses we test, the more likely we are to obtain nominally significant inferences. If we use the common approach of relaxing our threshold to accommodate the multiple testing, our Makwasinyi-Persian-Indian model for Mtwapa and Manda is robust. After adding Dinka to the right reference set (giving a base 3-way model $P= 0.26$ for Makwasinyi, Iranian, and Sahariya_MP), we added each of the 151 African HO populations as a source along with Makwasinyi, Iranian, and Sahariya_MP, but none produced a working model with $P>0.1$. We also tried switching between each of the African populations and Makwasinyi, putting Makwasinyi as a reference population and each HO African population as a source population, but none produced a working model with $P>0.1$.

We repeated the process of searching for alternative source populations for 487 Eurasian HO populations (with Dinka added to the right reference set), adding them as a fourth source or switching between Iranian and the Eurasian populations or between Sahariya_MP and the Eurasian populations. No Eurasian population worked as a fourth source. No other Eurasian population provided a working model with Iranian on the right, however other Persian-associated populations came close: Iran_Zoroastrian ($P=0.077$), Shia_Iranian_Hyderabad ($P=0.054$), India_Zoroastrian ($P=0.038$), and Iran_Non_Zoroastrian ($P=0.036$). Many Indian populations provided working models when switched with Sahariya_MP as well as populations from two other south Asian present-day countries, Pakistan and Bandladesh, that were historically part of Hindu empires. This indicates that Sahariya_MP is not the exact source, but simply a proxy for Indian-associated ancestry in the Manda group.

The average ancestry proportions for Mtwapa are 0.58±0.01 Makwasinyi-associated, 0.36±0.01 Persian-associated, and 0.06±0.01 Indian-associated. The average ancestry proportions for Manda are 0.34±0.01 Makwasinyi-associated, 0.59±0.02 Persian-associated, and 0.07±0.02 Indian-associated. The major proportions of ancestry for each Mtwapa and Manda individual using the final right reference set and a 3-way model with source populations of Makwasinyi, Iranian, and Sahariya_MP are summarized in Figure 2 and Extended Data Table 2 and Table S8. There are three highlighted outliers, which may have slightly different ancestry from the main group, or alternatively small amounts of undetected contamination (either could cause the general models to fail for these specific individuals).

| **Population** | **Mtwapa/Manda individual** | **P value** | **Makwasinyi coefficient** | **Iranian coefficient** | **Sahariya_MP coefficient** |
| --- | --- | --- | --- | --- | --- |
| Mtwapa | I17410 | 0.773537 | 0.44±0.02 | 0.46±0.03 | 0.1±0.02 |
|  | I19381 | 0.025539 | 0.6±0.02 | 0.34±0.02 | 0.06±0.02 |
|  | I19387 | 0.239045 | 0.59±0.02 | 0.34±0.03 | 0.07±0.02 |
|  | I19388 | 0.766829 | 0.55±0.03 | 0.41±0.05 | 0.05±0.05 |
|  | I19394 | 0.001969 | 0.62±0.02 | 0.37±0.02 | 0.01±0.02 |
|  | I19392 | 0.205746 | 0.76±0.02 | 0.19±0.03 | 0.05±0.02 |
|  | I19384 | 0.054742 | 0.5±0.02 | 0.42±0.02 | 0.08±0.02 |
|  | I19414 | 0.276103 | 0.61±0.01 | 0.32±0.02 | 0.07±0.02 |
|  | I19416 | 0.825797 | 0.7±0.02 | 0.23±0.03 | 0.07±0.02 |
|  | I19420 | 0.312661 | 0.66±0.02 | 0.26±0.02 | 0.08±0.02 |
|  | I19411 | 0.271986 | 0.61±0.02 | 0.29±0.03 | 0.1±0.02 |
|  | I19423 | 0.712401 | 0.46±0.02 | 0.48±0.04 | 0.05±0.04 |
|  | I19408 | 0.767363 | 0.54±0.03 | 0.41±0.05 | 0.05±0.05 |
|  | I19413 | 0.422078 | 0.57±0.01 | 0.36±0.02 | 0.06±0.02 |
|  | I19417 | 0.427363 | 0.65±0.02 | 0.31±0.02 | 0.04±0.02 |
|  | I19401 | 0.046828 | 0.57±0.01 | 0.39±0.02 | 0.04±0.02 |
|  | I23660 | 0.105636 | 0.62±0.03 | 0.34±0.04 | 0.04±0.04 |
|  | I23662 | 0.002357 | 0.53±0.02 | 0.41±0.02 | 0.05±0.02 |
|  | I23548 | 0.762237 | 0.64±0.02 | 0.3±0.03 | 0.06±0.03 |
|  | I17409 | 0.402058 | 0.5±0.01 | 0.43±0.02 | 0.07±0.02 |
|  | I24975 | 0.474807 | 0.47±0.03 | 0.42±0.06 | 0.1±0.06 |
|  | I19391 | 0.26526 | 0.48±0.02 | 0.44±0.02 | 0.08±0.02 |
|  | I19409 | 0.234715 | 0.45±0.02 | 0.47±0.03 | 0.08±0.03 |
|  | I13611 | 0.505993 | 0.59±0.02 | 0.36±0.02 | 0.05±0.02 |
|  | I17412 | 0.061578 | 0.49±0.01 | 0.41±0.02 | 0.11±0.02 |
|  | I17413 | 0.166367 | 0.67±0.02 | 0.26±0.02 | 0.07±0.02 |
|  | I23558 | 0.000323 | 0.42±0.01 | 0.48±0.02 | 0.1±0.02 |
|  | I19415 | 0.763372 | 0.63±0.01 | 0.32±0.02 | 0.05±0.02 |
|  | I23561 | 0.21538 | 0.5±0.01 | 0.45±0.02 | 0.05±0.02 |
|  | I19386 | 0.419929 | 0.55±0.02 | 0.47±0.03 | -0.02±0.03 |
|  | I21475 | 0.254544 | 0.71±0.01 | 0.27±0.02 | 0.03±0.01 |
| Manda | I7934 | 0.700058 | 0.3±0.01 | 0.63±0.02 | 0.08±0.02 |
|  | I7939 | 0.39307 | 0.57±0.02 | 0.38±0.04 | 0.05±0.04 |
|  | I7941 | 0.640198 | 0.46±0.02 | 0.42±0.04 | 0.12±0.04 |
|  | I7938 | 0.439132 | 0.42±0.02 | 0.54±0.03 | 0.04±0.03 |

Table S8: P values and coefficients for the Makwasinyi, Iranian, and Sahariya_MP source populations when individually modeling each of the Mtwapa and Manda individuals using the final right reference set.

*Lindi*

The individual from Lindi (I14001) can be modeled as having forager-associated and Bantu-associated ancestry, without Eurasian-associated ancestry. The right set of reference populations is Tatar_Siberian, Onge, Juang, Palliyar, Italian_Central, Cameroon_Mbo, Han, Ami, Turkish, Lebanese, Jew_Yemenite, Kalash, Murut, Kankanaey, Mbuti, Taa_West, Ju_hoan_North, Tunisian, Dinka, Ethiopia_Mursi, and Masai. A number of southern or eastern African forager-associated populations provide working models as do a number of Bantu-associated populations. A model with Malawi_Yao (89.2±0.3%) and Hadza (10.8±0.3%) as surrogate sources fits ($P=$0.60) even if we challenge it by adding other present-day African HO populations to the reference set in *qpAdm*. While the model cannot be broken by adding other Bantu-associated populations to the right, some other Bantu-associated populations do provide working alternative models in place of Malawi_Yao, but with different proportions of ancestry. Since many Bantu-associated populations have some forager-associated ancestry as well, it is difficult to determine the exact proportion of each type of ancestry.

*Songo Mnara*

The PCA and ADMIXTURE plots indicate that the Songo Mnara individuals are diverse in their major ancestry components; some of the individuals also lived at different times from one another, in contrast with other studied sites. We therefore model each individual separately.

Individual I7944, from 1516-1667 calCE, appears on the PCA to be on the Pastoral Neolithic to Pastoral Iron Age cline, without West Eurasian-associated admixture. This individual can be modeled as having about 95-98% PIA and 2-5% PN associated ancestry, depending on the specific PN source populations used for the model. We then tested $f4\left( S\text{ongo Mnara I7994},\text{ Kenya\_IA\_Pastoral\_published}, \text{X, Brazil\_MHolocene} \right)$, and no population tested was able to provide evidence that there is a significant genetic ancestry difference between the Songo Mnara individual and IA pastoral individuals (Table S9).

| X | f4 | Z score | SNPs used |
| --- | --- | --- | --- |
| Tanzania_Lindi_Swahili | -0.000653 | -0.938 | 399652 |
| Russia_IA_EarlySarmatian | -0.000448 | -0.786 | 316819 |
| Pakistan_H_Aligrama | -0.000246 | -0.527 | 434048 |
| Pakistan_Loebanr_IA_published | -0.000206 | -0.441 | 419236 |
| Egypt_ThirdIntermediatePeriod | -0.000287 | -0.425 | 292214 |
| China_YR_LBIA | -0.000101 | -0.243 | 434152 |
| Kenya_Manda_Swahili | -0.000163 | -0.234 | 245057 |
| Tanzania_Kilwa_Swahili | -0.000127 | -0.186 | 408427 |
| Taiwan_Gongguan | -0.000082 | -0.146 | 357677 |
| Kenya_Mtwapa_Swahili_o | -0.000317 | -0.106 | 11054 |
| Kazakhstan_Tasmola_Saka_IA | -0.000041 | -0.085 | 428173 |
| Uzbekistan_Dzharkutan_BA_1 | -0.000001 | -0.003 | 431226 |
| Vanuatu_150BP | 0.000011 | 0.024 | 433993 |
| Italy_Sardinia_Medieval | 0.000035 | 0.061 | 304661 |
| Italy_North_EarlyMedieval_Langobards_1 | 0.000033 | 0.068 | 430167 |
| Kenya_PastoralN | 0.000040 | 0.083 | 434741 |
| Kenya_PastoralN_Elmenteitan | 0.000045 | 0.088 | 431063 |
| Italy_Sardinia_IA_Punic_1 | 0.000051 | 0.092 | 375905 |
| Spain_IA | 0.000087 | 0.201 | 434753 |
| Kazakhstan_Saka_IA | 0.000107 | 0.224 | 405308 |
| Kazakhstan_Maitan_MLBA_Alakul | 0.000105 | 0.242 | 434532 |
| Pakistan_Loebanr_IA | 0.000110 | 0.271 | 435072 |
| Congo_NgongoMbata_Protohistoric | 0.000261 | 0.290 | 175112 |
| Tanzania_PN_IA_o | 0.000203 | 0.297 | 378627 |
| Greece_BA_Mycenaean | 0.000141 | 0.298 | 433496 |
| Spain_Roman | 0.000136 | 0.303 | 434355 |
| Congo_Kindoki_Protohistoric | 0.000206 | 0.307 | 345935 |
| Pakistan_Gogdara_IA | 0.000169 | 0.330 | 410128 |
| Kenya_Mtwapa_Swahili | 0.000154 | 0.337 | 434825 |
| Botswana_Xaro_EIA | 0.000222 | 0.338 | 422593 |
| Cameroon_SMA_published | 0.000212 | 0.351 | 434721 |
| Spain_EBA | 0.000179 | 0.415 | 435025 |
| Kenya_EarlyPastoralN | 0.000253 | 0.425 | 357424 |
| Kazakhstan_LIA_Georgievsky_published | 0.000282 | 0.464 | 371346 |
| Tanzania_Zanzibar_1300BP | 0.000490 | 0.489 | 145555 |
| Italy_Sardinia_BA_Nuragic | 0.000232 | 0.495 | 434867 |
| Pakistan_Udegram_IA | 0.000216 | 0.504 | 434658 |
| Pakistan_H_Barikot | 0.000247 | 0.534 | 428945 |
| China_WLR_BA | 0.000337 | 0.561 | 258145 |
| Kenya_LukenyaHill_PastoralN | 0.000422 | 0.566 | 243937 |
| Spain_Islamic | 0.000242 | 0.575 | 435106 |
| Kenya_Kakapel_LSA_Kansyore | 0.000425 | 0.583 | 284979 |
| Iran_C_TepeHissar | 0.000279 | 0.601 | 428596 |
| Taiwan_Hanben_IA | 0.000245 | 0.606 | 435099 |
| Kazakhstan_Berel_IA | 0.000242 | 0.618 | 435092 |
| Italy_North_EarlyMedieval_Langobards_2 | 0.000275 | 0.619 | 434097 |
| Uzbekistan_SappaliTepe_BA | 0.000272 | 0.622 | 434950 |
| Pakistan_Medieval_Udegram_Ghaznavid | 0.000388 | 0.626 | 330557 |
| Israel_MLBA | 0.000282 | 0.651 | 432315 |
| Jordan_LBA | 0.000306 | 0.691 | 435014 |
| Kazakhstan_Tasmola_EIA | 0.000280 | 0.725 | 435105 |
| India_RoopkundB | 0.000331 | 0.731 | 431721 |
| Tanzania_Luxmanda_3000BP | 0.000513 | 0.738 | 373152 |
| China_Upper_YR_IA | 0.000347 | 0.760 | 424953 |
| Kenya_Historic_2 | 0.000447 | 0.763 | 430267 |
| Kenya_Makwasingi_Swahili | 0.000452 | 0.783 | 426544 |
| Turkey_Alalakh_MLBA | 0.000353 | 0.804 | 434938 |
| Kenya_Kakapel_LIA | 0.000599 | 0.834 | 335694 |
| India_RoopkundA | 0.000347 | 0.865 | 434764 |
| Turkmenistan_Gonur_BA_1 | 0.000402 | 0.875 | 421862 |
| Kazakhstan_IA_Chanchar_published | 0.000517 | 0.879 | 367741 |
| Syria_Ebla_EMBA | 0.000412 | 0.885 | 430198 |
| Hungary_Langobard | 0.000392 | 0.886 | 433714 |
| Spain_IA_published | 0.000532 | 0.913 | 317336 |
| Pakistan_Katelai_IA | 0.000373 | 0.923 | 434901 |
| Iran_GanjDareh_Historic | 0.000561 | 0.925 | 391091 |
| Kenya_PastoralN_published | 0.000592 | 0.971 | 384493 |
| Spain_Islamic_Zira | 0.000521 | 0.975 | 421238 |
| China_YR_LN | 0.000407 | 0.983 | 435066 |
| Kazakhstan_Sarmatian_IA | 0.000435 | 0.994 | 434653 |
| Morocco_Iberomaurusian | 0.000546 | 0.998 | 434399 |
| Pakistan_H_SaiduSharif | 0.000418 | 0.999 | 434976 |
| Spain_Medieval_published | 0.000627 | 1.001 | 276670 |
| Italy_North_EarlyMedieval_Langobards_3 | 0.000478 | 1.008 | 430970 |
| Tanzania_PN | 0.000536 | 1.027 | 424019 |
| Kazakhstan_Georgievsky_MBA_published | 0.000672 | 1.038 | 330966 |
| Kazakhstan_Shoendykol_MLBA_Fedorovo | 0.000526 | 1.058 | 428362 |
| Iran_IA_HajjiFiruz | 0.000736 | 1.086 | 254520 |
| Spain_Visigoth_Granada | 0.000532 | 1.107 | 424837 |
| Pakistan_Aligrama_IA_published | 0.000591 | 1.132 | 416098 |
| Kenya_MoloCave_PastoralN | 0.000765 | 1.135 | 397301 |
| Pakistan_H_Butkara | 0.000565 | 1.185 | 412095 |
| Tanzania_Pemba_600BP_published | 0.000910 | 1.262 | 360300 |
| Kenya_LSA | 0.000934 | 1.319 | 344251 |
| Uganda_Munsa_LIA | 0.001262 | 1.487 | 192029 |
| Kazakhstan_Chanchar_MBA_published | 0.000978 | 1.550 | 375396 |
| Pakistan_Katelai_IA_published | 0.000739 | 1.617 | 411630 |
| Pakistan_Barikot_IA | 0.000932 | 1.791 | 377195 |
| Kenya_HyraxHill_PastoralN | 0.001430 | 1.882 | 252924 |

Table S9: Results of an f4 statistic of the form $f4\left( S\text{ongo Mnara I7994},\text{ Kenya\_IA\_Pastoral\_published}, \text{X, Brazil\_MHolocene} \right)$. No Z score is greater than 3 or less than -3, which indicates that based on the populations tested, there is no evidence of a significant ancestry difference between Songo Mnara individual I7994 and Kenya_IA_Pastoral_published.

Individual I19547, from 1508-1648 calCE, can be modeled with *qpAdm* ($P=0.13$) as having 84.2±1.9% Bantu-associated, 7.8±2.3% Malagasy, and 8.1±1.3% Arabian ancestry using the HO dataset (modeled with Malawi_Ngoni, Madagascar_North, and Yemeni_Highlands_Raymah), using the final set of right reference populations with Dinka, Georgian, Jordanian, BedouinB, Russian, and Mbuti added to the right as well. Some southern African populations with forager ancestry cause the model to degrade by two orders of magnitude when cycled on the right, but when placed on the left do not produce a working model. This suggests that the Bantu-associated surrogate, Malawi_Ngoni, does not have the correct type or the correct amount of forager ancestry.

Individual I19549, from 1629-1794 calCE can be modeled as having Bantu-associated and Arabian-associated ancestry sources. Using Tanzania_Pemba_600BP_published, the individual buried at Makangale cave in Pemba ca. 600BP, as the Bantu-associated source and the same final set of right reference populations as with Mtwapa and Manda, the two sources that fit with $P>0.1$ are Emirati and Yemeni_Highlands_Raymah. If we lower the P value threshold to 0.04, a number of other Arabian peninsula populations provide working models as well. Given that many populations on the Arabian Peninsula already have varying amounts of Bantu-associated ancestry (see Figure 1C) and that we do not know the exact Arabian source, we cannot determine the exact proportions of admixture. Modeling with the Pemba (Makangale) individual from 600BP as the Bantu-associated source and several present-day genotyped Arabian Peninsula populations, we receive a range of 20-27% Arabian-associated ancestry. If, alternatively, we use this individual’s 2^nd^ or 3^rd^ degree relative, individual I19547 buried at Songo Mnara, as one source and if we add Georgian, Jordanian, BedouinB, and Russian to the final right set of reference populations, we obtain feasible, successful models with Yemeni_Desert and Yemeni_Highlands_Raymah. When we model individual I19549 on the 1240K dataset using individual I19547 (the 2^nd^ or 3^rd^ degree relative) as a source and a right reference set of Altaian.DG, Iranian.DG, Jew_Yemenite.DG, Onge.DG, Brahmin.DG, Spanish.DG, Han.DG, Ami.DG, Kalash.DG, Dusun.DG, Igorot.DG, Yoruba.DG, Dinka.DG, Mozabite.DG, Mende.DG, Hadza_1.DG, and Mbuti.DG, we get a working model of 59% ancestry from individual I19547 and 41% ancestry from Kenya_IA_Pastoral_published ($P=0.33$). When we try to break this model by cycling through 88 African or Eurasian populations form the 1240K dataset, only Kenya_LSA is able to degrade the model to $0.01<P<0.1$, indicating that we are including too much or too little or not the right type of forager ancestry in our model. The HO dataset does not have sufficient PN- or PIA-associated proxy sources and our 1240K dataset does not have sufficient Yemeni-associated proxy sources, and so we are not able to compare between these two potential sources without introducing dataset-specific biases.

Individual I19550, from 1412-1446 calCE, can be modeled ($P=0.21$) as deriving from Bantu-associated (67.3±2.1% Lindi), Persian-associated (25.2±3.7% Iranian), and Indian-associated (7.5±3.9% Sahariya_MP) sources. The set of right group reference populations in this model is the same as the final right set used for Mtwapa and Manda individuals. This model is robust to adding any other Eurasian or African HO population to the reference set. Out of 748 cycled populations, only two populations degrade the model to $0.01<P<0.1$, Mende, with West African-associated ancestry, and Dogra, with Indian-associated ancestry. This likely indicates that the surrogate sources are not the historically exact sources, but rather proxies for the correct sources. Individuals 5 (I19548), 6 (I19552), and 7 (I19551), appear in the PCA to have similar Eurasian and Indian ancestry (Figure 1B); however, their coverage is too low to confidently model them.

*Kilwa*

The single Kilwa individual can be modeled ($P= 0.27$) as an admixture of 24.3±1.2% Persian-associated ancestry and 75.7±1.2% Bantu-associated ancestry, with the latter proxied by the individual from Lindi, who also has forager ancestry. The Human Origins genotyped Iranian population was used as the Persian proxy source and the same final right reference set as with Mtwapa and Manda. Other present-day HO populations from areas associated with ancient Persia, including present-day Iran and the Caucasus, as well as an Iranian-Indian population living in India, also provide working models with similar proportions of ancestry (*Table S9*). Here, too, we represent the Persian-associated ancestry component with the HO Iranian population.

| **Populations that provide working models** | **P value** | **Bantu-associated coefficient** | **Persian-associated coefficient** | **Approximate latitude** | **Approximate longitude** |
| --- | --- | --- | --- | --- | --- |
| Avar_outlier2 | 0.301 | 0.77±0.01 | 0.24±0.01 | 41.22 | 48.30 |
| Iranian | 0.274 | 0.76±0.01 | 0.24±0.01 | 33.70 | 51.25 |
| Iran_Non_Zoroastrian | 0.265 | 0.76±0.01 | 0.24±0.01 | 34.01 | 51.76 |
| Azeri | 0.228 | 0.76±0.01 | 0.24±0.01 | 41.07 | 47.68 |
| Avar_outlier1 | 0.227 | 0.76±0.01 | 0.24±0.01 | 42.41 | 46.43 |
| Iran_Zoroastrian | 0.216 | 0.76±0.01 | 0.24±0.01 | 32.75 | 52.78 |
| Ezid | 0.171 | 0.76±0.01 | 0.24±0.01 | 36.90 | 46.60 |
| Kurd | 0.170 | 0.76±0.01 | 0.24±0.01 | 44.58 | 39.42 |
| Lezgin | 0.157 | 0.77±0.01 | 0.23±0.01 | 42.12 | 48.18 |
| Ossetian | 0.156 | 0.76±0.01 | 0.24±0.01 | 43.06 | 44.18 |
| Russia_Abkhasian | 0.145 | 0.77±0.01 | 0.23±0.01 | 43.00 | 41.02 |
| Karachai | 0.137 | 0.76±0.01 | 0.24±0.01 | 43.94 | 42.52 |
| Adygei | 0.137 | 0.77±0.01 | 0.24±0.01 | 44.39 | 39.31 |
| Kumyk | 0.127 | 0.76±0.01 | 0.24±0.01 | 43.25 | 46.58 |
| Tabasaran | 0.126 | 0.77±0.01 | 0.23±0.01 | 41.96 | 47.82 |
| Abkhasian | 0.125 | 0.77±0.01 | 0.23±0.01 | 43.00 | 41.02 |
| Shia_Iranian_Hyderabad | 0.125 | 0.76±0.01 | 0.24±0.01 | 17.39 | 78.49 |
| Avar | 0.123 | 0.77±0.01 | 0.23±0.01 | 41.82 | 47.37 |
| Ingushian | 0.123 | 0.77±0.01 | 0.24±0.01 | 43.52 | 44.58 |
| Iranian_Bandari | 0.112 | 0.74±0.01 | 0.26±0.01 | 27.18 | 56.27 |
| Balkar | 0.109 | 0.76±0.01 | 0.24±0.01 | 43.48 | 43.62 |
| Abazin | 0.108 | 0.76±0.01 | 0.24±0.01 | 44.15 | 42.20 |
| Circassian | 0.102 | 0.76±0.01 | 0.24±0.01 | 44.20 | 41.83 |

Table S11: Persian-associated population that produce working models for a 2-way model for the individual buried at Kilwa.

Initially, four African individuals or groups from the 1240K African dataset provided feasible and successful models (using the final right reference set). But we then took each of these four groups and added it as a source with the other three as references in addition to the final right reference set. Only the Lindi individual provided a working model.

| 1240K African groups | P value of model with this 1240K African group on the left | P value of model with this 1240K African group on the left and with the other three 1240K African groups on the right |
| --- | --- | --- |
| Cameroon_SMA_published | 0.279 | 0.094 |
| Tanzania_Lindi_Swahili | 0.274 | 0.230 |
| Kenya_Makwasingi_Swahili | 0.152 | 0.056 |
| Kenya_Kakapel_LIA | 0.138 | 0.047 |

Table S12: Testing for the best-fitting African proxy source for modeling the individual buried at Kilwa.

We tested the robustness of the African proxy source being the individual buried at Lindi by cycling through the 1240K coanalyzed African dataset on the left and on the right. Given that there are only a few published ancient individuals of high coverage on the 1240K SNP set with Bantu-associated ancestry in eastern Africa, it is expected that our Bantu-associated source would be a proxy rather than an accurate source for the correct population. Only populations with pastoral-associated ancestry on the right reduce the P value to below 0.01. But the Kilwa individual cannot be modeled well with the addition of PN-associated ancestry, and so it is likely indicative of too much PN-associated ancestry in our modeling. In fact, if a PN population is added as a third source, the P value is low, indicating a poor fit, and the PN coefficient is negative.

We next tested the robustness of the Eurasian component of the model. We challenged it by cycling populations into the right group reference set. Populations with South and East Asian ancestry added as a reference population slightly reduce the P value ($P<0.1$), but do not completely break the model ($P>0.01$). Adding an East Asian or a South Asian population as a third source does not significantly improve the model, but also does not significantly degrade the model. Because there are a different number of populations on the left in a two-source versus in a three-source model, the degrees of freedom differ, and so the P values cannot be compared. Altogether, using *qpAdm*, we cannot rule out nor claim either South Asian or East Asian admixture within the Kilwa sample. But if there is South or East Asian ancestry, the coefficient in such a case provides about 0.1-1.2% of the ancestry and is very close to zero within a single standard error. No HO African population completely breaks the model when cycled on the right, but a couple of northern African populations slightly reduce the P value ($0.05<P<0.1$).

While there is also no separate detectable Indian component of ancestry, if instead of a Persian source, we use a mixed Persian-Indian source, we do produce working models (Figure 2C). Populations along the Indian-Iranian cline as well as artificially produced populations of about 80% or more Iranian individuals and the rest Indian Pulliyar individuals do provide working models when substituted in place of the Iranian source (see **Supplementary qpAdm modeling: qpAdm Iranian/Indian proportions**). This could be explained by the presence of a small percentage of Indian ancestry (approximately 0.1% as determined by qpAdm runs with the individual buried at Lindi as the African proxy source, Iranian as the Persian proxy source and Sahariya_MP as the Indian proxy source), which falls below the limits of our ability to detect it definitively. Thus, the Kilwa individual may have a single homogeneous Persian-Indian admixed population as the source of Eurasian-associated ancestry as is the case in the Manda and Mtwapa groups and in Individual I19550 from Songo Mnara.

*Present-day populations from Kilifi, Lamu, and Mombasa*

We assembled published data from people living in Lamu (31 individuals), Kilifi (35 individuals), and Mombasa (23 individuals) [51]. A PCA of these present-day individuals projected onto the same PCA space as in Figure 1, shows many of the coastal individuals close to the Bantu-PN cline with some bearing higher proportions of Eurasian ancestry (Figure 3B). We remove the two individuals with the greatest proportion of West Eurasian ancestry as determined by the ancestral reference source maximized by Israel_Natufian_published in the ADMIXTURE graph of Extended Data Figure 1, and we treat them as outliers.

When we model the present-day coastal populations in a two-way model with a Bantu-associated source and cycle through ancient and present-day populations, we find that the Kilifi, Lamu, and Mombasa groups are well modeled with pastoralist populations – both PN and PIA. We also find that the Kilifi and Lamu groups can also be well modeled with medieval Swahili populations as the second source. So, we fixed both a Bantu-associated source (the individual buried at Lindi or the individual from 600BP buried at Pemba) and a PIA source (Kenya_IA_Pastoral_published) and cycled through ancient and modern populations allowing for a third source. The only similar 3-way model for Kilifi, Lamu, and Mombasa is with a medieval Swahili-associated source (the individual from Kilwa) as a third population. To represent the Pastoral Iron Age ancestry, we used two individuals: I8892 (Nakuru, Ilkek Mounds, GsJj66) and I12381 (Laikipia District Burial Site, GoJl45).

| **Present-day coastal population** | **P value** | **Kilwa** | **Lindi** | **PIA** |
| --- | --- | --- | --- | --- |
| Kilifi | 0.592 | 0.14±0.04 | 0.84±0.03 | 0.03±0.03 |
| Lamu | 0.359 | 0.12±0.04 | 0.86±0.03 | 0.02±0.03 |
| Mombasa | 0.346 | 0.06±0.04 | 0.79±0.03 | 0.15±0.03 |
| **Present-day coastal population** | **P value** | **Kilwa** | **Pemba 600BP** | **PIA** |
| Kilifi | 0.155 | 0.14±0.04 | 0.8±0.03 | 0.06±0.05 |
| Lamu | 0.142 | 0.1±0.04 | 0.84±0.03 | 0.06±0.05 |
| Mombasa | 0.094 | 0.05±0.03 | 0.76±0.03 | 0.19±0.05 |

In all *qpAdm* runs with present-day coastal populations, our right populations are Tatar_Siberian, Onge, Juang, Palliyar, Italian_Central, Cameroon_Mbo ,Han, Ami, Turkish, Lebanese, Jew_Yemenite, Kalash, Murut, Kankanaey, Dinka, Ethiopia_Mursi, Mende, Hadza1, Georgian, Masai, Egyptian, Yoruba, and Iran_Zoroastrian. And the populations that we investigated as potential sources are Botswana_Xaro_EIA, Cameroon_SMA_published, Congo_Kindoki_Protohistoric, Congo_NgongoMbata_Protohistoric, Egypt_ThirdIntermediatePeriod, Kenya_EarlyPastoralN, Kenya_Historic_2, Kenya_HyraxHill_PastoralN, Kenya_IA_Pastoral_published, Kenya_Kakapel_LIA, Kenya_Kakapel_LSA_Kansyore, Kenya_LSA, Kenya_LukenyaHill_PastoralN, Kenya_Makwasingi_Swahili, Kenya_Manda_Swahili, Kenya_MoloCave_PastoralN, Kenya_Mtwapa_Swahili, Kenya_PastoralN, Kenya_PastoralN_Elmenteitan, Kenya_PastoralN_published, Malawi_Chencherere_LSA_5200BP, Morocco_Iberomaurusian, Tanzania_Kilwa_Swahili, Tanzania_Lindi_Swahili, Tanzania_Luxmanda_3000BP, Tanzania_Pemba_600BP_published, Tanzania_PN, Tanzania_SongoMnara_Swahili, Tanzania_Zanzibar_1300BP, Uganda_Munsa_LIA, Malawi_Yao, Emirati, Iranian, Shia_Iranian_Hyderabad, Spanish, Italian_Central, Iranian_Bandari, Iran_DinkhaTepe_BA_IA_1, Iran_DinkhaTepe_BA_IA_2, Yemeni_Highlands, Yemeni_Desert, Yemen_Modern, Yemeni, Saudi, and Ethiopia_4500BP_published.SG.

None of the cycled populations, when added to the right-hand outgroups, were able to completely break the Lamu, Kilifi, and Mombasa models. Iran_DinkhaTepe_BA_IA_2 was able to reduce the P value of the model by two orders of magnitude, perhaps indicating that the individual buried at Kilwa is not the exact medieval Swahili source for present-day Swahili groups, but rather a close proxy. Otherwise, the Mtwapa group, the Makwasinyi group, and KPN are able to degrade the model by one order of magnitude ($0.01<P<0.1$), indicating that the proxy pastoralist source and proxy medieval Swahili source are not the exact sources, but rather proxies.

#### Modeling present-day coastal and inland present-day groups

*Present-day inland populations from present-day Kenya and Tanzania*

We also present *qpAdm* admixture models for present-day inland populations in present-day Kenya and Tanzania. The goal here is not to show the full landscape of present-day Kenyan and Tanzanian variation, but rather to represent the very complex situation with approximate surrogate Bantu-associated, pastoralist-associated, and forager-associated ancestry components (see SI for proportions). We caution that there exists high covariance between the sources, and thus it is in some cases more informative to assess the total sum of their proportions of ancestry. But because there is no ideal distinction about which populations are best combined, the models should be viewed as representative estimates, rather than unique sets of ancestral origins.

We modeled numerous present-day samples that were shotgun sequenced, and so for consistency, both our left and right groups consisted solely of shotgun sequenced samples. Our right reference set comprised Han.DG, Onge.DG, Spanish.DG, Turkish.DG, Armenian.DG, Jew_Yemenite.DG, Khomani_San.DG, Mozabite.DG, Cameroon_SMA.DG, and Iranian.DG and our left set of sources was Yoruba.DG, Amhara.DG, Ethiopia_4500BP.DG, and Mbuti.DG. Yoruba represents West African-associated ancestry. Amhara represents pastoralist-associated ancestry. The similarity between some Ethiopian populations and pastoralist populations can be seen in Extended Data Figure 1. Amhara might also include some West Eurasian ancestry as well. Ethiopia_4500BP.DG, also known as Mota or Bayira, and Mbuti.DG are forager ancestries that exist is small proportions in the area. Mbuti.DG has high covariance with Ethiopia_4500BP.DG and Amhara.DG, and so we combine all three populations into a single pastoralists-and-foragers surrogate population. Note that there was also high covariance between Ethiopia_4500BP.DG and Yoruba.DG, and so we could have also reasonably chosen to combine those two populations and have Bantu-forager with pastoralist-forager sources. There is no ideal combination, likely because while these surrogate sources are very distinct, the actual Bantu-associated and pastoralist-associated sources present in these regions of Kenya and Tanzania likely both had various forager-associated admixture, making either distinction arbitrary. Thus, we caution against using these proportions as exact models.

|  | **P value** | **Yoruba.DG** | **pastoralists + foragers** |
| --- | --- | --- | --- |
| Kenya_Makwasinyi_Swahili | 0.51 | 0.69±0.05 | 0.31±0.09 |
| Kikuyu.DG | 0.25 | 0.58±0.06 | 0.42±0.11 |
| Maasai (Masai.DG) | 0.25 | 0.39±0.06 | 0.61±0.12 |
| Iraqw.DG | 0.15 | 0.14±0.07 | 0.86±0.14 |
| Okiek (Ogiek.DG) | 0.81 | 0.46±0.06 | 0.55±0.11 |
| Elmolo.DG | 0.41 | 0.46±0.06 | 0.55±0.11 |
| Rendille.DG | 0.56 | 0.2±0.06 | 0.8±0.12 |
| Sengwer.DG | 0.10 | 0.54±0.06 | 0.46±0.11 |
| LWK.SG | 0.02 | 0.82±0.04 | 0.18±0.07 |
| Somali.DG | 0.50 | 0.25±0.08 | 0.75±0.16 |
| Luo.DG | 0.21 | 0.81±0.06 | 0.19±0.11 |
| Luhya.DG | 0.48 | 0.92±0.06 | 0.08±0.11 |
| BantuKenya.DG | 0.06 | 0.82±0.06 | 0.18±0.11 |
| Hadza_1.DG | 0.04 | 0.14±0.08 | 0.86±0.16 |
| Hadza_2.DG | 0.59 | 0.26±0.08 | 0.75±0.15 |
| Sandawe.DG | 0.12 | 0.16±0.06 | 0.84±0.12 |

Table S13: Uniform four-source model for present-day inland Kenyan and Tanzanian populations, where three of the sources are combined as pastoralists+foragers because of their high covariance.

#### *qpAdm* on the X-chromosome and autosomes for sex-bias determination

We calculate the Z-score for each sex-bias calculation, which tells us how many standard deviations we are away from the mean of equal male and female ancestry contributions

$Z= \frac{p_{A}-p_{X}}{\sqrt{\sigma_{A}^{2}+\sigma_{X}^{2}}}$.

If there is an extreme sex bias, where the population under question contributed excess male ancestry, we would expect to observe significant differences between the X-chromosome and autosomal admixture proportions ($Z\gtrsim2$). If the population under question contributed excess female ancestry, we would expect $Z\lesssim-2$. To compute a plausible standard error on the X-chromosome, we used smaller jackknife bins in the *qpAdm* analysis with option blgsize: 0.005 and fancyf4: yes. However, given the low number of measured SNPs on the X-chromosome (fewer than 30,000), standard errors are generally large, and thus the power to measure bias is limited.

When calculating sex-bias in the Makwasinyi population with the individual buried at Pemba ca. 600BP as the Bantu proxy, the number of SNPs used for *qpAdm* calculations on all the autosomes was 688,274 and the number of SNPs used for *qpAdm* calculations on the X chromosome was 22,485. The model coefficients for the autosomes are $C_{A}=0.787\pm0.010$Bantu-associated and $C_{A}=0.213\pm0.010$KPN-associated. The Model coefficients for chromosome X are $C_{X}=0.753\pm0.037$Bantu-associated and $C_{X}=0.247\pm0.037$PN-associated. This gives $Z= -0.89$ for the Bantu-associated proportion of ancestry.

For calculating sex-bias in the Mtwapa and Manda groups, we apply a three-source *qpAdm* model with the 1240K-capture Makwasinyi source and the diploid shotgun Iranian.DG and Indian Mala.DG sources to the autosomes and X-chromosome for both Mtwapa and Manda. The set of reference populations was Russian.DG, Onge.DG, Brahmin.DG, Italian_North.DG, Cameroon_SMA.DG, Han.DG, Ami.DG, Kalash.DG, Dusun.DG, Igorot.DG, Iran_TepeAbdulHosein_N.SG, and Indian_GreatAndaman_100BP.SG. Mtwapa and Manda autosomal *qpAdm* calculations use 919,225 and 540,549 SNPs, respectively, whereas their X chromosome *qpAdm* calculations use 31,682 and 21,233 SNPs, respectively. In the Manda group, we calculate entirely female Makwasinyi ancestry. We also infer male Iranian-associated ancestry and primarily female Mala-associated ancestry, however the uncertainty estimates are extremely high. In the Mtwapa group, the Iranian ancestry is derived from males, but while most of the Makwasinyi ancestry derives from females, a small portion seems to come from males as well. The Indian ancestry seems to be entirely or predominantly deriving from females.

To run sex-bias analyses in the Kilwa sample, we used another 2-way Bantu-Persian admixture model with higher SNP coverage using Pemba_600BP_published as the Bantu-associated proxy source, and the shotgun Iranian.DG as the Persian proxy source. The *qpAdm* model uses 678,103 SNPs for autosomal calculations and 23,891 SNPs for X chromosomal calculations. The set of reference populations was the same as for Mtwapa and Manda. Like Manda and Mtwapa, the Iranian component of the Kilwa individual is consistent with being male-derived and the Bantu-associated ancestry is consistent with being derived predominantly from females.

#### Ternary plot linear regression to determine the two admixing source populations

The percentages of ancestry in the two admixing source populations are calculated by finding the intersection point (depicted by the left circle in Figure 2B and Figure S3) between the Makwasinyi ternary axis (yellow line in Figure S3) and the linear regression line (red line in Figure S3) at $x_{m}=\frac{0.01301}{\sqrt{3}-0.09856}$ and $y_{m}=\sqrt{3} x_{m}$. The distance along the Makwasinyi ternary axis is then 1-$\sqrt{x_{m}^{2}+y_{m}^{2}}=0.9841$, which indicates that a nearly 100% Makwasinyi-like group admixed with a mixed Iranian-Indian group.

The intersection point (depicted by the right circle in Figure 2B and Figure S3) between the Indian ternary axis (which is the axis where the proportion of Makwasinyi ancestry is 0 and is depicted as the blue line in Figure S3) and the linear regression line (red line in Figure S3) is at $x_{i}=\frac{\sqrt{3}-0.01301}{\sqrt{3}+0.09856}$ and $y_{i}=-\sqrt{3} x_{i}+\sqrt{3}$. The distance along the Indian ternary axis is then $\sqrt{\left( 1-x_{i} \right)^{2}+y_{i}^{2}}=0.1219$. The intersection on the Iranian axis (green full and dotted lines in Figure S3) would be 0.8781. This indicates that the Iranian-Indian mixed group had approximately 12% Indian-associated ancestry and 88% Persian-associated ancestry.

We apply a Block Jackknife approach to correct for correlation among individuals and to acquire a more accurate standard error. We create 603 bins of 991 SNPs each and run the *qpAdm* analyses and calculate the Makwasinyi and Indian axis interception points 603 times, where each time we remove one bin of 991 SNPs from consideration. The Block Jackknife gives a mean and standard error of the Makwasinyi axis intersection point as $0.9843\pm0.0253$ and those of the Indian axis intersection point as $0.1221\pm0.0276$. The standard error was calculated as ${SE}_{jackknife} = \sqrt{\frac{n-1}{n}\sum_{1}^{n} \left( x_{i}-\underline{x} \right)}$. Any proportion greater than 100% Makwasinyi is impossible, and results from noise arising from issues such as inexact surrogate source populations and the low coverage of samples.

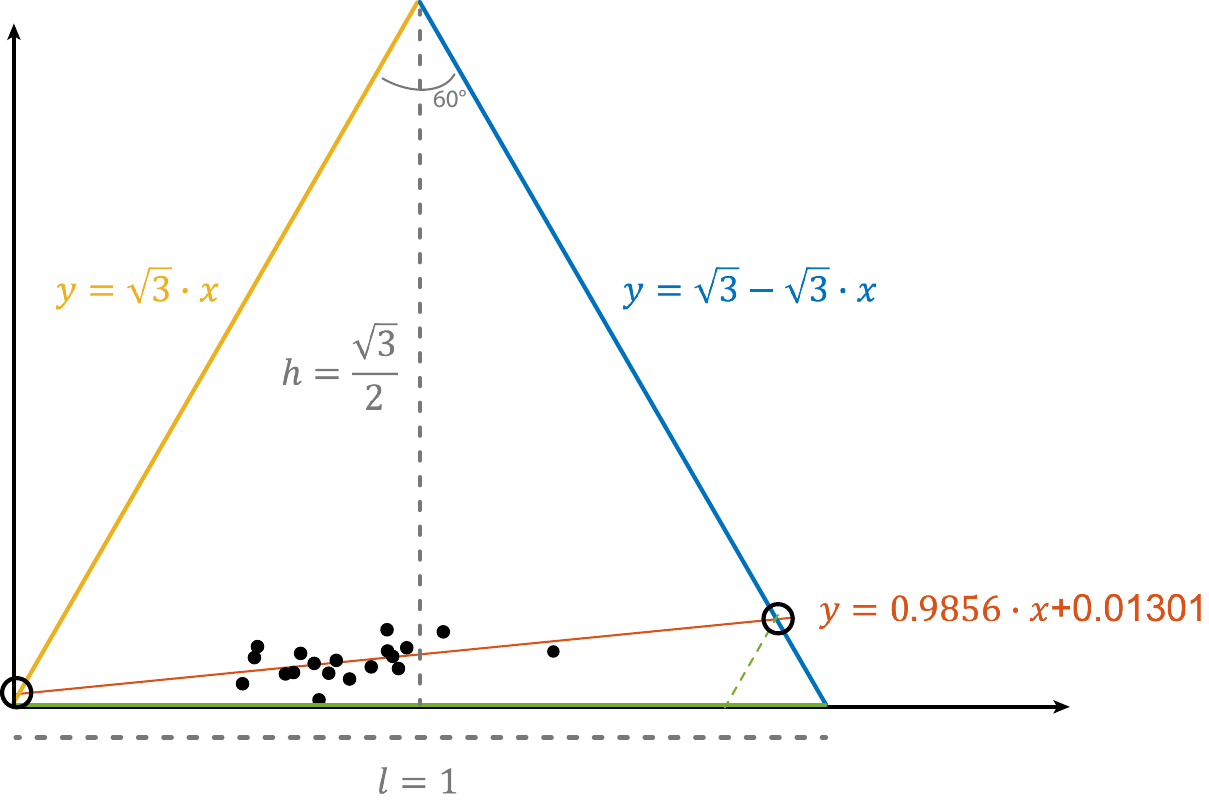

Figure S3: Schematic depicting the linear regression of the Mtwapa and Manda high coverage points on the ternary plot.

#### *qpAdm* Iranian/Indian proportions

We estimated the ratio of Persian to Indian ancestry in Mtwapa, Manda, and Kilwa by creating a mixed population with varying numbers of Iranian individuals and Pulliyar individuals from India. We formed a 2-source model with this mixed population and Makwasinyi for Manda and Mtwapa or the individual buried at Lindi for the individual buried at Kilwa and for Songo Mnara individual I19550. We analyzed the *qpAdm* P values for each Iranian/Indian proportion level of this 2-source model to find possible Iranian/Indian proportions (see Figure 2C). The Mtwapa group is determined to allow for $84.2\%\pm3.8\%$to $86.8\%\pm3.8\%$ Iranian ancestry with the rest being Indian ancestry. The Manda group is determined to allow for $84.2\%\pm3.8\%$to $97.4\%\pm3.8\%$ Iranian ancestry, the Kilwa individual allows for $89.5\%\pm3.8\%$to $100\%$ Iranian ancestry, and the Songo Mnara individual I19550 allows for 55.3%$\pm3.8\%$ to 100% Iranian ancestry. The Songo Mnara individual has much lower SNP count than the other medieval Swahili coastal groups and so the range of acceptable proportions is much larger. We would expect some amount of variation resulting from differences in the exact Iranian and Indian ancestry proportions in the admixing individuals in each location, as well as inconsistencies arising from having *qpAdm* surrogate populations and not having the precise source populations.

#### *qpAdm* Mtwapa_lowI to Mtwapa_highI or Manda

We detected inhomogeneity in Eurasian- and African-associated ancestry in the Mtwapa individuals (Figure 2B), with a base of around 30% Persian-associated ancestry, above which this component of ancestry is highly variable, sometimes with as much as 50% or more West Eurasian ancestry. To determine if the West Eurasian ancestry is consistent with deriving from a single source or could have derived from heterogeneous sources, we divided the Mtwapa individuals into Mtwapa_lowI and Mtwapa_highI. Mtwapa_lowI individuals have less than 40% Persian-associated ancestry and Mtwapa_highI individuals had greater than 40% Persian-associated ancestry according to the *qpAdm* model.

We used Mtwapa_lowI as a source population for Mtwapa_highI in further *qpAdm* modeling and cycled through 495 West Eurasian present-day populations (using the final right reference set). We found that Iranian, Iran_Non_Zoroastrian, and Iran_Zoroastrian, were not viable proxy sources for the more recent West Eurasian ancestry. However, Emirati and Iranian_Bandari, which border the Strait of Hormuz, did provide working models. Makrani and Shia_Iranian_Hyderabad, which have Indian-associated and Persian-associated ancestry also provide working models. We also used Mtwapa_lowI as a source in a two-way model for Manda, and this analysis instead favors Persian-associated populations for the extra West Eurasian-relatedness in Manda. The individuals buried at Manda pre-date the samples buried at Mtwapa, suggesting that a Persian-associated population was the predominant West Eurasian source population among Manda-like populations along the Northeast Swahili coast up until approximately 1600 CE (Figure S9). And the later Mtwapa-like population, has a different added ancestry component, which may resemble the ancestry components in the Emirati population. We caution that the Emirati samples we analyze are genetically heterogenous, which must reflect modern movements of people in the United Arab Emirates. Thus, the fact that the Emirati sample we have is a good proxy for the additional West Eurasian ancestry in Mtwapa_highI compared to Mtwapa_lowI, cannot mean that it is the true source population, but instead that the genetic heterogeneity that is observed today in the Emirati samples is part of a long-standing process of interaction between people in the region, which just happens to capture the additional Eurasian ancestry components that distinguish Mtwapa_highI from Mtwapa_lowI and are not found in Manda. Such a scenario is in fact plausible, since Oman itself was a site of interaction between Persian, Arabian, African, and Indian people.

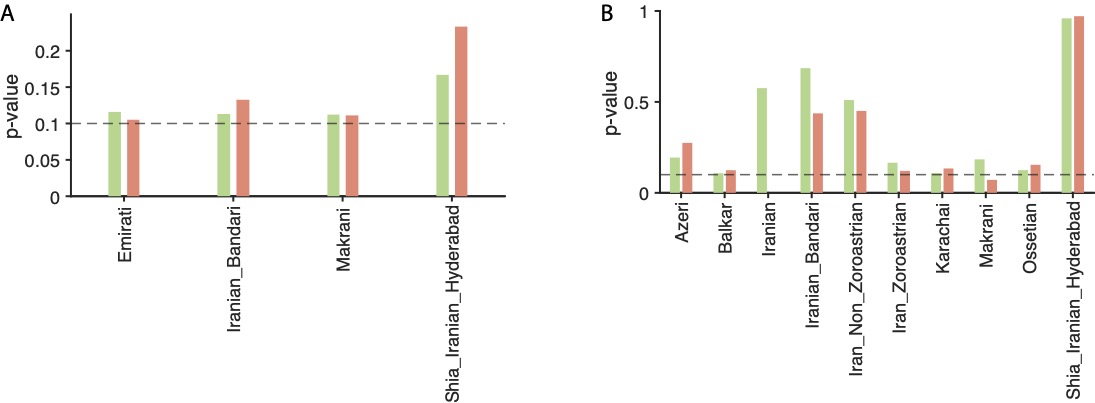

Figure S4: A) Additional WE ancestry source in Mtwapa_highI relative to Mtwapa_lowI; *qpAdm* models with and without Iranian as part of the right-hand reference set are orange and green, respectively. Emirati is a sample from an area that was under Omani control when Oman also controlled part of the Swahili coast. Another successful proxy for the additional West Eurasian ancsetry is Shia_Iranian_Hyderabad, which has a mixture of Persian-associated and South Asian-associated ancestry components, but Iranian by itself did not provide a successful model. This could be indicative of Persian-Indian related ancestry but also of a combination of Persian-Indian and Omani ancestry, as Oman had many South Asian and Persian individuals since its control included parts of present-day Pakistan and present-day Iran. Models with and without Iranian as part of the right-hand reference set are orange and green, respectively. B) Additional WE ancestry source in Manda relative to Mtwapa_lowI shows that Persian-associated ancestry, including Iranian, provide working models.

In order to investigate the differences between Arabian and Iranian ancestry in the Mtwapa and Manda groups, we analyzed another dataset from Fernandes et al. of genotyping data from present-day individuals from Saudi Arabia, Yemen, Dubai, Oman, and Iran [52]. Many individuals in this dataset had significant recent African mixture [52], reflecting events in recent centuries after the date range of the medieval Swahili coast individuals we analyze. We removed individuals who had a value of less than -0.0006 on eigenvector 1 when projected onto the same PCA as in Fig 1 in order to try to limit to individuals with the smallest proportions of African ancestry. We then used Mtwapa individual I17413 as a source for all other Mtwapa individuals and the Manda group. We chose Mtwapa I17413 because this individual has high coverage and low Eurasian admixture. We cycled through the five Persian-associated or Arabian-associated source populations on the left, using the other four as the right reference group. We also applied the same analysis using Manda as the source population. Since Mtwapa generally has less Eurasian ancestry compared to Manda, we also use the individual from 600BP buried at Pemba Island as a third proxy source. When Manda is used a source population in feasible and successful ($P>0.1$) models, most Mtwapa individuals allow for added Arabian ancestry (Table S14). Only a handful allow for added Iranian ancestry, with only a single individual allowing for only added Iranian ancestry. When Mtwapa individual I17413 is used as a source, nearly every feasible and successful model allows Iran as a source, and Iran is the only allowed source in six cases (Table S15). In many individuals both where Manda and where Mtwapa individual I17413 are used as sources, Arabian populations are allowed for other Mtwapa individuals (including individual I17413), but not for Manda. We repeated this analysis with other Mtwapa individuals (Table S16 and Table S17) and overall obtained similar results. Because of the overlap of ancestry and shared IBD segments among these present-day individuals [52], we cannot use them to determine the correct added ancestry source in some individuals of the Mtwapa group, and it appears that it could plausibly be Arabian rather than Persian or somewhere along the Arabian-Persian cline compared to in the Manda group.

| **Arabian/Persian source** | **Target Mtwapa Individual** | **P value** | **Arabian/Persian source coefficient** | **Manda source coefficient** | **Tanzania_Pemba_600BP_published source coefficient** |
| --- | --- | --- | --- | --- | --- |
| Iran | I19381 | 0.85899 | 0.05±1.02 | 0.57±1.33 | 0.39±0.33 |
| Oman | I19387 | 0.927352 | 0.11±0.26 | 0.49±0.27 | 0.4±0.12 |
| Dubai | I19388 | 0.485193 | 0.54±1.44 | 0.07±1.03 | 0.39±0.52 |
| Iran | I19388 | 0.957457 | 0.25±0.77 | 0.1±0.62 | 0.65±0.46 |
| SaudiArabia | I19388 | 0.973913 | 0.15±0.43 | 0.49±0.23 | 0.37±0.46 |
| Dubai | I19394 | 0.343121 | 0.13±0.36 | 0.45±0.42 | 0.43±0.12 |
| Oman | I19394 | 0.92502 | 0.19±0.37 | 0.42±0.41 | 0.39±0.12 |
| Yemen | I19394 | 0.672465 | 0.01±0.11 | 0.48±0.19 | 0.52±0.17 |
| Dubai | I19392 | 0.162854 | 0.25±0.33 | 0.18±0.29 | 0.57±0.12 |
| Oman | I19392 | 0.304369 | 0.05±0.4 | 0.31±0.35 | 0.64±0.14 |
| SaudiArabia | I19392 | 0.29808 | 0.35±0.27 | 0.34±0.11 | 0.31±0.27 |
| Yemen | I19392 | 0.873135 | 0.14±0.11 | 0.48±0.16 | 0.38±0.17 |
| Dubai | I19384 | 0.753386 | 0.02±0.27 | 0.6±0.29 | 0.38±0.09 |
| Iran | I19384 | 0.795347 | 0.26±0.58 | 0.29±0.74 | 0.45±0.19 |
| Oman | I19384 | 0.835009 | 0.02±0.34 | 0.59±0.33 | 0.39±0.09 |
| SaudiArabia | I19384 | 0.895073 | 0.07±0.18 | 0.63±0.1 | 0.31±0.2 |
| Yemen | I19384 | 0.733201 | 0±0.08 | 0.64±0.15 | 0.36±0.14 |
| Dubai | I19414 | 0.860362 | 0.06±0.18 | 0.63±0.2 | 0.31±0.07 |
| Oman | I19414 | 0.178234 | 0.07±0.24 | 0.63±0.25 | 0.3±0.07 |
| SaudiArabia | I19414 | 0.172835 | 0.03±0.16 | 0.7±0.08 | 0.27±0.17 |
| Dubai | I19416 | 0.409524 | 0.16±0.29 | 0.43±0.41 | 0.41±0.19 |
| Oman | I19416 | 0.743443 | 0.14±0.31 | 0.42±0.41 | 0.44±0.18 |
| SaudiArabia | I19416 | 0.573988 | 0.33±0.32 | 0.58±0.18 | 0.09±0.35 |
| Yemen | I19416 | 0.336536 | 0.09±0.13 | 0.61±0.28 | 0.3±0.31 |
| Dubai | I19420 | 0.881424 | 0.12±0.28 | 0.41±0.29 | 0.47±0.1 |
| Oman | I19420 | 0.329535 | 0.23±0.33 | 0.29±0.32 | 0.48±0.1 |
| Yemen | I19420 | 0.303193 | 0.06±0.12 | 0.5±0.19 | 0.45±0.24 |
| SaudiArabia | I19411 | 0.098168 | 0.3±0.27 | 0.6±0.13 | 0.1±0.3 |
| Yemen | I19411 | 0.55578 | 0.08±0.11 | 0.81±0.19 | 0.11±0.2 |
| Oman | I19423 | 0.461957 | 0.05±0.64 | 0.79±0.47 | 0.17±0.25 |
| Dubai | I19408 | 0.568215 | 0.09±0.61 | 0.66±0.59 | 0.25±0.22 |
| Iran | I19408 | 0.686144 | 0.57±1.16 | 0.07±1.27 | 0.36±0.25 |
| Oman | I19408 | 0.868629 | 0.08±0.62 | 0.65±0.56 | 0.26±0.22 |
| SaudiArabia | I19408 | 0.831443 | 0.2±1.51 | 0.68±0.31 | 0.12±1.3 |
| Yemen | I19408 | 0.877843 | 0.09±0.35 | 0.84±0.46 | 0.07±0.66 |
| Dubai | I19413 | 0.462456 | 0.01±0.18 | 0.55±0.2 | 0.43±0.07 |
| Oman | I19413 | 0.730511 | 0.06±0.21 | 0.52±0.22 | 0.42±0.07 |
| Dubai | I19417 | 0.81294 | 0.17±0.24 | 0.48±0.27 | 0.34±0.12 |
| Oman | I19417 | 0.456249 | 0.19±0.33 | 0.45±0.34 | 0.35±0.14 |
| SaudiArabia | I19417 | 0.571363 | 0.03±0.2 | 0.62±0.13 | 0.35±0.2 |
| Yemen | I19417 | 0.556677 | 0.08±0.11 | 0.62±0.16 | 0.3±0.17 |
| Dubai | I23662 | 0.983607 | 0.09±0.44 | 0.66±0.49 | 0.25±0.14 |
| Oman | I23662 | 0.23798 | 0.31±0.49 | 0.42±0.52 | 0.27±0.14 |
| Yemen | I23662 | 0.20674 | 0.03±0.12 | 0.69±0.2 | 0.28±0.21 |
| Dubai | I17409 | 0.033731 | 0.05±0.24 | 0.63±0.26 | 0.31±0.07 |
| Yemen | I17409 | 0.155351 | 0.06±0.07 | 0.8±0.13 | 0.14±0.13 |
| Dubai | I19409 | 0.948918 | 0.07±0.28 | 0.66±0.27 | 0.27±0.14 |
| Iran | I19409 | 0.654787 | 0.25±0.63 | 0.44±0.83 | 0.32±0.26 |
| Oman | I19409 | 0.629958 | 0.07±0.32 | 0.66±0.3 | 0.27±0.15 |
| Dubai | I13611 | 0.923116 | 0.23±0.19 | 0.49±0.21 | 0.28±0.08 |
| Oman | I13611 | 0.507684 | 0.27±0.22 | 0.45±0.23 | 0.28±0.08 |
| SaudiArabia | I13611 | 0.302684 | 0.07±0.19 | 0.69±0.09 | 0.24±0.21 |
| Yemen | I13611 | 0.362064 | 0.06±0.08 | 0.64±0.13 | 0.29±0.13 |
| SaudiArabia | I17413 | 0.916624 | 0.02±0.16 | 0.47±0.07 | 0.51±0.17 |
| Dubai | I19415 | 0.888268 | 0.4±0.16 | 0.23±0.17 | 0.37±0.07 |
| Oman | I19415 | 0.290958 | 0.5±0.19 | 0.15±0.19 | 0.35±0.07 |
| SaudiArabia | I19415 | 0.484393 | 0.05±0.18 | 0.55±0.08 | 0.41±0.18 |
| Yemen | I19415 | 0.378026 | 0.16±0.07 | 0.52±0.11 | 0.32±0.12 |
| Dubai | I23561 | 0.621793 | 0.09±0.19 | 0.66±0.21 | 0.25±0.07 |
| SaudiArabia | I23561 | 0.002533 | 0.06±0.22 | 0.79±0.1 | 0.15±0.22 |
| Dubai | I19386 | 0.616448 | 0.22±0.39 | 0.24±0.43 | 0.55±0.2 |
| Iran | I19386 | 0.526766 | 0.25±0.52 | 0.1±0.92 | 0.66±0.48 |
| Oman | I19386 | 0.987653 | 0.23±0.36 | 0.23±0.34 | 0.54±0.19 |
| SaudiArabia | I19386 | 0.538959 | 0.39±0.96 | 0.38±0.2 | 0.23±1.02 |
| Yemen | I19386 | 0.547339 | 0.1±0.18 | 0.35±0.23 | 0.56±0.28 |
| Dubai | I21475 | 0.575435 | 0.36±0.16 | 0.13±0.17 | 0.51±0.06 |
| Oman | I21475 | 0.409734 | 0.44±0.2 | 0.08±0.2 | 0.48±0.06 |
| SaudiArabia | I21475 | 0.846687 | 0±0.15 | 0.43±0.07 | 0.57±0.16 |
| Yemen | I21475 | 0.514369 | 0.13±0.06 | 0.38±0.1 | 0.5±0.11 |

Table S14: Table of successful and feasible models using Manda as a source along with an Arabian or Persian-associated population and a Bantu-associated population.

| **Arabian/Persian source** | **Target Mtwapa Individual or Manda group** | **P value** | **Arabian/Persian source coefficient** | **I17413 source coefficient** |
| --- | --- | --- | --- | --- |
| Iran | I17410 | 0.97 | 0.5±0.13 | 0.5±0.13 |
| Iran | I19381 | 0.23 | 0.06±0.15 | 0.94±0.15 |
| Dubai | I19387 | 0.32 | 0.07±0.18 | 0.93±0.18 |
| Iran | I19387 | 0.44 | 0.25±0.16 | 0.75±0.16 |
| Oman | I19387 | 0.23 | 0.21±0.22 | 0.8±0.22 |
| Dubai | I19388 | 0.87 | 0.44±0.38 | 0.56±0.38 |
| Iran | I19388 | 0.89 | 0.37±0.44 | 0.63±0.44 |
| Oman | I19388 | 0.95 | 0.63±0.49 | 0.37±0.49 |
| Yemen | I19388 | 0.76 | 0.08±0.26 | 0.93±0.26 |
| Dubai | I19394 | 0.52 | 0.06±0.18 | 0.94±0.18 |
| Iran | I19394 | 0.40 | 0.22±0.15 | 0.79±0.15 |
| Oman | I19394 | 0.19 | 0.11±0.21 | 0.89±0.21 |
| SaudiArabia | I19392 | 0.32 | 0.28±0.19 | 0.72±0.19 |
| Yemen | I19392 | 0.75 | 0.04±0.13 | 0.97±0.13 |
| Dubai | I19384 | 0.15 | 0.24±0.11 | 0.76±0.11 |
| Iran | I19384 | 0.94 | 0.25±0.1 | 0.75±0.1 |
| Oman | I19384 | 0.20 | 0.3±0.13 | 0.7±0.13 |
| Dubai | I19414 | 0.35 | 0.27±0.1 | 0.74±0.1 |
| Iran | I19414 | 0.50 | 0.31±0.09 | 0.7±0.09 |
| Oman | I19414 | 0.16 | 0.31±0.12 | 0.69±0.12 |
| Yemen | I19414 | 0.12 | 0±0.07 | 1±0.07 |
| Dubai | I19416 | 0.92 | 0.03±0.24 | 0.97±0.24 |
| Iran | I19416 | 0.50 | 0.14±0.21 | 0.86±0.21 |
| SaudiArabia | I19416 | 0.34 | 0.08±0.21 | 0.92±0.21 |
| Yemen | I19416 | 0.33 | 0.01±0.13 | 0.99±0.13 |
| Dubai | I19420 | 0.24 | 0.09±0.18 | 0.91±0.18 |
| Iran | I19420 | 0.35 | 0.16±0.15 | 0.85±0.15 |
| Oman | I19420 | 0.67 | 0.2±0.19 | 0.8±0.19 |
| Dubai | I19411 | 0.83 | 0±0.22 | 1±0.22 |
| Iran | I19411 | 0.97 | 0.05±0.19 | 0.95±0.19 |
| Oman | I19411 | 0.86 | 0.05±0.25 | 0.95±0.25 |
| SaudiArabia | I19411 | 0.79 | 0.02±0.18 | 0.99±0.18 |
| Yemen | I19411 | 0.81 | 0.01±0.11 | 0.99±0.11 |
| Dubai | I19408 | 0.19 | 0.85±0.78 | 0.15±0.78 |
| Oman | I19408 | 0.43 | 0.2±2.38 | 0.8±2.38 |
| Dubai | I19413 | 0.35 | 0.18±0.11 | 0.82±0.11 |
| Iran | I19413 | 0.78 | 0.22±0.1 | 0.78±0.1 |
| Oman | I19413 | 0.74 | 0.26±0.12 | 0.74±0.12 |
| Yemen | I19413 | 0.97 | 0±0.07 | 1±0.07 |
| Dubai | I19417 | 0.54 | 0.06±0.19 | 0.94±0.19 |
| Iran | I19417 | 0.64 | 0.08±0.18 | 0.93±0.18 |
| Oman | I19417 | 0.84 | 0.18±0.23 | 0.82±0.23 |
| Yemen | I19417 | 0.67 | 0.05±0.11 | 0.95±0.11 |
| Dubai | I19401 | 0.17 | 0.07±0.12 | 0.93±0.12 |
| Iran | I19401 | 0.36 | 0.08±0.11 | 0.92±0.11 |
| Oman | I19401 | 0.17 | 0.11±0.14 | 0.89±0.14 |
| Iran | I23660 | 0.66 | 0.21±0.43 | 0.79±0.43 |
| Oman | I23660 | 0.14 | 0.17±0.49 | 0.83±0.49 |
| SaudiArabia | I23660 | 0.15 | 0.26±0.27 | 0.74±0.27 |
| Yemen | I23660 | 0.15 | 0.19±0.17 | 0.81±0.17 |
| Dubai | I23662 | 0.74 | 0.34±0.14 | 0.66±0.14 |
| Iran | I23662 | 0.83 | 0.36±0.13 | 0.64±0.13 |
| Oman | I23662 | 0.70 | 0.39±0.15 | 0.61±0.15 |
| Yemen | I23662 | 0.54 | 0.06±0.09 | 0.94±0.09 |
| Dubai | I23548 | 0.14 | 0.21±0.15 | 0.79±0.15 |
| Iran | I23548 | 0.18 | 0.4±0.14 | 0.6±0.14 |
| Dubai | I17409 | 0.19 | 0.29±0.1 | 0.71±0.1 |
| Iran | I17409 | 0.86 | 0.29±0.09 | 0.71±0.09 |
| Oman | I17409 | 0.23 | 0.33±0.11 | 0.67±0.11 |
| Dubai | I24975 | 0.47 | 0.48±2.12 | 0.52±2.12 |
| Iran | I24975 | 0.28 | 0.34±0.81 | 0.66±0.81 |
| Oman | I24975 | 0.55 | 0.25±1.4 | 0.75±1.4 |
| Iran | I19391 | 0.17 | 0.57±0.13 | 0.43±0.13 |
| Dubai | I19409 | 0.11 | 0.23±0.28 | 0.78±0.28 |
| Iran | I19409 | 0.68 | 0.37±0.21 | 0.63±0.21 |
| Oman | I19409 | 0.10 | 0.3±0.38 | 0.7±0.38 |
| Dubai | I13611 | 0.68 | 0.36±0.1 | 0.64±0.1 |
| Iran | I13611 | 0.47 | 0.32±0.11 | 0.68±0.11 |
| Oman | I13611 | 0.76 | 0.4±0.12 | 0.6±0.12 |
| Yemen | I13611 | 0.16 | 0.07±0.07 | 0.93±0.07 |
| Iran | I17412 | 0.26 | 0.38±0.09 | 0.62±0.09 |
| Iran | I23558 | 0.21 | 0.56±0.09 | 0.44±0.09 |
| Dubai | I19415 | 0.35 | 0.27±0.12 | 0.73±0.12 |
| Iran | I19415 | 0.25 | 0.22±0.11 | 0.78±0.11 |
| Oman | I19415 | 0.61 | 0.35±0.13 | 0.65±0.13 |
| Yemen | I19415 | 0.89 | 0.11±0.07 | 0.89±0.07 |
| Dubai | I23561 | 0.18 | 0.23±0.1 | 0.77±0.1 |
| Iran | I19386 | 0.73 | 0.04±0.31 | 0.96±0.31 |
| SaudiArabia | I19386 | 0.72 | 0.15±0.25 | 0.85±0.25 |
| Dubai | I21475 | 0.18 | 0.01±0.11 | 1±0.11 |
| Iran | I21475 | 0.89 | 0.07±0.1 | 0.93±0.1 |
| Oman | I21475 | 0.26 | 0.07±0.13 | 0.93±0.13 |
| Yemen | I21475 | 0.20 | 0.04±0.07 | 0.96±0.07 |
| Iran | Manda | 0.93 | 0.66±0.09 | 0.34±0.09 |

Table S15: Table of successful and feasible models using Mtwapa I17413 as a source along with an Arabian or Persian-associated population.

| **Arabian/Persian source** | **Target Mtwapa Individual or Manda group** | **P value** | **Arabian/Persian source coefficient** | **I19414 source coefficient** |
| --- | --- | --- | --- | --- |
| Iran | I17410 | 0.383738 | 0.45±0.14 | 0.55±0.14 |
| Oman | I17410 | 0.34686 | 0.44±0.17 | 0.56±0.17 |
| SaudiArabia | I19381 | 0.830074 | 0.02±0.12 | 0.98±0.12 |
| Dubai | I19388 | 0.424408 | 0.44±0.86 | 0.56±0.86 |
| Iran | I19388 | 0.869522 | 0.14±1.11 | 0.86±1.11 |
| SaudiArabia | I19388 | 0.789645 | 0.32±0.22 | 0.68±0.22 |
| Yemen | I19388 | 0.690431 | 0.21±0.21 | 0.79±0.21 |
| Iran | I19394 | 0.595972 | 0.02±0.22 | 0.99±0.22 |
| Yemen | I19392 | 0.260598 | 0.15±0.1 | 0.85±0.1 |
| Oman | I19384 | 0.408639 | 0.02±0.18 | 0.98±0.18 |
| SaudiArabia | I19384 | 0.363376 | 0.03±0.11 | 0.97±0.11 |
| Yemen | I19384 | 0.406977 | 0.02±0.07 | 0.98±0.07 |
| Iran | I19416 | 0.538569 | 0.06±0.24 | 0.94±0.24 |
| SaudiArabia | I19420 | 0.228109 | 0.09±0.14 | 0.91±0.14 |
| SaudiArabia | I19411 | 0.510538 | 0.2±0.13 | 0.8±0.13 |
| Dubai | I19408 | 0.982535 | 0.36±0.43 | 0.64±0.43 |
| Iran | I19408 | 0.606262 | 0.48±0.33 | 0.52±0.33 |
| Oman | I19408 | 0.470859 | 0.66±0.51 | 0.34±0.51 |
| SaudiArabia | I19408 | 0.271368 | 0.01±0.32 | 1±0.32 |
| Yemen | I19408 | 0.383491 | 0.04±0.22 | 0.96±0.22 |
| Oman | I19417 | 0.491141 | 0.18±0.27 | 0.82±0.27 |
| Iran | I23660 | 0.249224 | 0.23±0.42 | 0.77±0.42 |
| Iran | I23662 | 0.961625 | 0.04±0.2 | 0.97±0.2 |
| Iran | I17409 | 0.949402 | 0±0.14 | 1±0.14 |
| Dubai | I24975 | 0.436355 | 0±1.05 | 1±1.05 |
| Iran | I24975 | 0.505158 | 0.23±0.72 | 0.77±0.72 |
| Oman | I24975 | 0.449584 | 0.93±1.97 | 0.07±1.97 |
| Dubai | I19409 | 0.797635 | 0.29±0.17 | 0.71±0.17 |
| Iran | I19409 | 0.944135 | 0.34±0.18 | 0.66±0.18 |
| Oman | I19409 | 0.848239 | 0.34±0.19 | 0.66±0.19 |
| Yemen | I19409 | 0.982947 | 0.09±0.11 | 0.91±0.11 |
| Dubai | I13611 | 0.80128 | 0.02±0.16 | 0.98±0.16 |
| Oman | I13611 | 0.706668 | 0.05±0.19 | 0.95±0.19 |
| SaudiArabia | I13611 | 0.804539 | 0.05±0.1 | 0.95±0.1 |
| Yemen | I13611 | 0.832972 | 0.05±0.07 | 0.95±0.07 |
| Iran | I17412 | 0.115688 | 0.15±0.13 | 0.85±0.13 |
| Yemen | I17413 | 0.119445 | 0±0.07 | 1±0.07 |
| Iran | I23558 | 0.384101 | 0.33±0.13 | 0.67±0.13 |
| Dubai | I19415 | 0.116973 | 0.04±0.16 | 0.96±0.16 |
| Iran | I19415 | 0.24438 | 0.01±0.15 | 0.99±0.15 |
| Oman | I19415 | 0.359015 | 0.15±0.18 | 0.85±0.18 |
| SaudiArabia | I19415 | 0.181691 | 0.01±0.12 | 0.99±0.12 |
| Yemen | I19415 | 0.188982 | 0.07±0.07 | 0.93±0.07 |
| Iran | I23561 | 0.363935 | 0.07±0.15 | 0.93±0.15 |
| SaudiArabia | I19386 | 0.295919 | 0.13±0.2 | 0.87±0.2 |
| Yemen | I19386 | 0.286443 | 0.07±0.14 | 0.93±0.14 |
| Iran | Manda | 0.786577 | 0.44±0.12 | 0.56±0.12 |

Table S16: Table of successful and feasible models using Mtwapa I19414 as a source along with an Arabian or Persian-associated population.

| **Arabian/Persian source** | **Target Mtwapa Individual or Manda group** | **P value** | **Arabian/Persian source coefficient** | **I21475 source coefficient** |
| --- | --- | --- | --- | --- |
| Iran | I17410 | 0.949414 | 0.29±0.2 | 0.71±0.2 |
| Oman | I17410 | 0.109409 | 0.4±0.29 | 0.6±0.29 |
| Dubai | I19387 | 0.168615 | 0.07±0.16 | 0.93±0.16 |
| Iran | I19387 | 0.524836 | 0.14±0.17 | 0.86±0.17 |
| Oman | I19387 | 0.166607 | 0.08±0.23 | 0.92±0.23 |
| Dubai | I19388 | 0.994973 | 0.31±0.35 | 0.69±0.35 |
| Iran | I19388 | 0.884677 | 0.18±0.44 | 0.82±0.44 |
| Oman | I19388 | 0.974447 | 0.37±0.43 | 0.63±0.43 |
| SaudiArabia | I19388 | 0.845169 | 0.02±0.39 | 0.98±0.39 |
| Yemen | I19388 | 0.825592 | 0.1±0.23 | 0.9±0.23 |
| Iran | I19394 | 0.195522 | 0.22±0.16 | 0.78±0.16 |
| SaudiArabia | I19392 | 0.261064 | 0.2±0.22 | 0.8±0.22 |
| Yemen | I19392 | 0.447341 | 0.02±0.13 | 0.98±0.13 |
| Iran | I19384 | 0.834285 | 0.14±0.1 | 0.86±0.1 |
| Dubai | I19416 | 0.825154 | 0.15±0.17 | 0.86±0.17 |
| Iran | I19416 | 0.981525 | 0.16±0.17 | 0.84±0.17 |
| Oman | I19416 | 0.849444 | 0.17±0.18 | 0.83±0.18 |
| SaudiArabia | I19411 | 0.36654 | 0.16±0.15 | 0.84±0.15 |
| Iran | I19423 | 0.223275 | 0.64±0.24 | 0.36±0.24 |
| Dubai | I19408 | 0.695407 | 0.36±0.6 | 0.64±0.6 |
| Iran | I19408 | 0.680982 | 0.48±0.32 | 0.52±0.32 |
| Oman | I19408 | 0.428727 | 0.93±0.82 | 0.07±0.82 |
| Dubai | I19413 | 0.203482 | 0.11±0.1 | 0.89±0.1 |
| Iran | I19413 | 0.692975 | 0.12±0.1 | 0.88±0.1 |
| Oman | I19413 | 0.221392 | 0.12±0.13 | 0.88±0.13 |
| Dubai | I19417 | 0.842231 | 0.18±0.16 | 0.82±0.16 |
| Iran | I19417 | 0.721989 | 0.09±0.18 | 0.91±0.18 |
| Oman | I19417 | 0.616157 | 0.26±0.22 | 0.74±0.22 |
| Yemen | I19417 | 0.470485 | 0.03±0.1 | 0.98±0.1 |
| SaudiArabia | I23660 | 0.467253 | 0.13±0.27 | 0.87±0.27 |
| Iran | I23662 | 0.12112 | 0.21±0.18 | 0.79±0.18 |
| Oman | I23662 | 0.165724 | 0.05±0.34 | 0.95±0.34 |
| Iran | I23548 | 0.287654 | 0.01±0.23 | 0.99±0.23 |
| Iran | I17409 | 0.677278 | 0.24±0.1 | 0.76±0.1 |
| Dubai | I24975 | 0.246996 | 0.51±0.31 | 0.49±0.31 |
| Iran | I24975 | 0.899644 | 0.49±0.26 | 0.51±0.26 |
| Oman | I24975 | 0.260865 | 0.6±0.41 | 0.4±0.41 |
| Iran | I19391 | 0.237279 | 0.43±0.14 | 0.57±0.14 |
| Dubai | I19409 | 0.322903 | 0.35±0.18 | 0.65±0.18 |
| Iran | I19409 | 0.450252 | 0.42±0.19 | 0.59±0.19 |
| Oman | I19409 | 0.305953 | 0.3±0.26 | 0.7±0.26 |
| Dubai | I13611 | 0.16218 | 0.32±0.11 | 0.68±0.11 |
| Iran | I13611 | 0.170518 | 0.25±0.12 | 0.75±0.12 |
| Oman | I13611 | 0.485048 | 0.27±0.15 | 0.73±0.15 |
| Dubai | I17413 | 0.179518 | 0.01±0.11 | 0.99±0.11 |
| SaudiArabia | I17413 | 0.170767 | 0.01±0.11 | 0.99±0.11 |
| Iran | I23558 | 0.30264 | 0.49±0.1 | 0.51±0.1 |
| Dubai | I19415 | 0.925049 | 0.2±0.12 | 0.8±0.12 |
| Iran | I19415 | 0.572521 | 0.1±0.13 | 0.9±0.13 |
| Oman | I19415 | 0.696052 | 0.26±0.15 | 0.74±0.15 |
| Yemen | I19415 | 0.369341 | 0.04±0.07 | 0.96±0.07 |
| Dubai | I19386 | 0.127306 | 0.16±0.23 | 0.84±0.23 |
| Iran | I19386 | 0.418348 | 0.19±0.26 | 0.81±0.26 |
| Oman | I19386 | 0.207068 | 0.3±0.25 | 0.7±0.25 |
| Iran | Manda | 0.737615 | 0.6±0.09 | 0.4±0.09 |

Table S17: Table of successful and feasible models using Mtwapa I21475 as a source along with an Arabian or Persian-associated population.

### DATES (Distribution of Ancestry Tracts of Evolutionary Signals)

We applied DATES [53] to the Makwasinyi set of individuals using KPN and the Yoruba population as a proxy for the Bantu-associated source. We estimate that admixture occurred 38.4±5.0 generations (here and what follows we quote one standard error) prior to the time the Makwasinyi individuals lived, with the result consistent with a single admixture pulse to the limits of our resolution. Using a 28±2 year-to-generation conversion estimate [54], we estimate that the average date of the admixture event occurred 1075±217 years before the estimated date for the Makwasinyi individuals, which of the dated samples, averaging to $1799\pm53$ CE. This puts the average admixture date at $724\pm223$ CE.

It should be noted that the admixture linkage disequilibrium in the DATES curve does not converge at zero, which indicates that despite the admixture occurring many generations ago, there remains some amount of substructure within this population. This likely results from close relatedness among Makwasinyi individuals.

We applied DATES to the Mtwapa individuals with a two-source admixture using Makwasinyi and Iranians as proxy sources (Extended Data Figure 2). The average number of elapsed generations between the admixture event and date of the Mtwapa population, as determined with DATES, is 18.999±0.972. By multiplying the elapsed generations by a generation to year estimate of 28±2, we calculate the number of elapsed years as 532±65. The average date for the isotopically dated Mtwapa samples is 1622±42 BP. This puts the admixture event as occurring on average at 1090±77 CE.

A similar DATES analysis for the Manda individuals results in a standard error that is over 70% the value of elapsed generations and so we substitute the Yoruba population in place of the Makwasinyi population, which provides a curve consistent with a single admixture pulse 17.961±5.750 generations ago, which is approximately 503±197 before the time of the Manda individuals. The average radiocarbon date of Manda individuals is 1523±34 CE, placing an average single pulse for the admixture event at 1020±200 CE. We applied a similar DATES curve to the Kilwa sample and obtained a curve corresponding to admixture at 16.963±2.592 generations ago. Unlike the Mtwapa and Manda individuals, we have no radiocarbon date of the Kilwa individual to use as a reference, and so we used archaeological context dte range of 1300-1600 CE. This puts the admixture at approximately 975±184 CE.

These DATES calculations are consistent with the admixture for each of the Swahili populations possibly occurring at roughly the same time.

1.	Emery, J.B., *A Short Account of Mombasa and Neighbouring Coast of Africa.* Journal of the Royal Geographical Society, 1883: p. 280-282.

2.	Kusimba, C.M., *The archaeology and ethnography of iron metallurgy on the Kenya coast*. 1993, Bryn Mawr College.

3.	Kusimba, C.M., S.B. Kusimba, and L. Dussubieux, *Beyond the Coastalscapes: Preindustrial Social and Political Networks in East Africa.* African Archaeological Review, 2013. **30**(4): p. 399-426.

4.	Chami, F.A., *A review of Swahili archaeology.* African Archaeological Review, 1998. **15**(3): p. 199-218.

5.	Chami, F., *The Tanzanian Coast in the Early First Millennium AD: an archaeology of the iron-working, farming communities*. Studies in African Archaeology. Vol. 7. 1994, Uppsala: Societas Archaeologica Uppsaliensis.

6.	Abungu, G., *Communities on the River Tana, Kenya: an archaeological study of relations between the delta and the river basin.* 1990.

7.	Fleisher, J. and S. Wynne-Jones, *Ceramics and the early Swahili: deconstructing the Early Tana Tradition.* African Archaeological Review, 2011. **28**(4): p. 245-278.

8.	Horton, M., *Shanga: The archaeology of a Muslim trading community on the coast of East Africa (British Institute in Eastern Africa).* 1996.

9.	Haaland, R., *Emergence of sedentism: new ways of living, new ways of symbolizing.* Antiquity, 1997. **71**(272): p. 374-385.

10.	Chittick, N., *Manda: Excavations at an island port on the Kenya coast.* 1984.

11.	Oka, R. and C. Kusimba, *Archaeology of trading systems 1: A theoretical survey.* Journal of Archaeological Research, 2008. **16**: p. 339-395.

12.	Gannon, M., *600-year-old Chinese coin found in Kenya*, in *NBC News*. 2013.

13.	Wynne-Jones, S., *Kilwa Kisiwani and Songo Mnara*, in *The Swahili World*. 2017, Routledge. p. 253-259.

14.	Garlake, P.S., *The early Islamic architecture of the East African coast*. 1966: London, Oxford U. P.

15.	Pradines, S. and P. Blanchard. *Kilwa al-Mulûk. Premier bilan des travaux de conservation-restauration et des fouilles archéologiques dans la baie de Kilwa, Tanzanie*. in *Annales Islamologiques*. 2005.

16.	Wynne-Jones, S., *The public life of the Swahili stonehouse, 14th–15th centuries AD.* Journal of anthropological archaeology, 2013. **32**(4): p. 759-773.

17.	Fleisher, J., *The complexity of public space at the Swahili town of Songo Mnara, Tanzania.* Journal of anthropological archaeology, 2014. **35**: p. 1-22.

18.	Wynne-Jones, S. and J. Fleisher, *The multiple territories of Swahili urban landscapes.* World Archaeology, 2016. **48**(3): p. 349-362.

19.	Wynne-Jones, S. and J. Fleisher, *Conservation, community archaeology, and archaeological mediation at Songo Mnara, Tanzania.* Journal of Field Archaeology, 2015. **40**(1): p. 110-119.

20.	Wynne-Jones, S., *The Archaeology of the Swahili World*, in *Oxford Research Encyclopedia of African History*. 2020.

21.	Skoglund, P., et al., *Reconstructing Prehistoric African Population Structure.* Cell, 2017. **171**(1): p. 59-71.e21.

22.	Chittick, N., *Kilwa: an Islamic trading city on the East African coast.* 1974.

23.	Wynne-Jones, S. and J. Fleisher, *Fifty years in the archaeology of the eastern African coast: a methodological history.* Azania: Archaeological Research in Africa, 2015. **50**(4): p. 519-541.

24.	Horton, M., et al., *The Chronology of Kilwa Kisiwani, AD 800–1500.* African Archaeological Review, 2022.

25.	Africa, B.I.i.E. 2016; Available from: <https://doi.org/10.5284/1038987>.

26.	Masao, F.T. *A NEWLY DISCOVERED MSA/LSA VARIANT OR MASASIAN: REPORT OF ARCHAEOLOGICAL INVESTIGATION OF SOUTH EASTERN TANZANIA*. 2015.

27.	Saanane, C.B., *Preliminary Report on Early Settlements and Archaeological Materials from Lindi Rural District, Lindi Region, Southeastern Tanzania.* International Journal of Geosciences, 2016. **7**(5): p. 655-668.

28.	Beyin, A. and K.P. Ryano, *Filling the Void: a Study of Sites Characterized by Levallois and Blade Technologies in the Kilwa Basin, Coastal Tanzania.* Journal of Paleolithic Archaeology, 2020. **3**(4): p. 1048-1094.

29.	Chami, F.A. and R. CHAMI, *Narosura pottery from the southern coast of Tanzania: first incontrovertible coastal Later Stone Age pottery.* Nyame akuma, 2001(56): p. 29-35.

30.	Kwekason, A.P., *Nkope: The early ironworking pottery tradition of southern coastal Tanzania.* African Archaeological Review, 2013. **30**(2): p. 145-167.

31.	Kwekason, A., *Holocene archaeology of the southern coast of Tanzania.* Dar es Salaam: E&D Vision Publishing, 2011.

32.	Pollard, E. and E.B. Ichumbaki, *Why land here? Ports and harbors in southeast Tanzania in the early second millennium AD.* The Journal of Island and Coastal Archaeology, 2017. **12**(4): p. 459-489.

33.	Pollard, E., *Safeguarding Swahili trade in the fourteenth and fifteenth centuries: a unique navigational complex in south-east Tanzania.* World Archaeology, 2011. **43**(3): p. 458-477.

34.	Pawlowicz, M., *Modelling the Swahili past: the archaeology of Mikindani in southern coastal Tanzania.* Azania: Archaeological Research in Africa, 2012. **47**(4): p. 488-508.

35.	Becker, F., *Traders,‘big men’and prophets: political continuity and crisis in the Maji Maji rebellion in Southeast Tanzania.* The Journal of African History, 2004. **45**(1): p. 1-22.

36.	Price, A.L., et al., *Principal components analysis corrects for stratification in genome-wide association studies.* Nature Genetics, 2006. **38**(8): p. 904-909.

37.	Patterson, N., A.L. Price, and D. Reich, *Population Structure and Eigenanalysis.* PLOS Genetics, 2006. **2**(12): p. e190.

38.	Nakatsuka, N., et al., *The promise of discovering population-specific disease-associated genes in South Asia.* Nature Genetics, 2017. **49**(9): p. 1403-1407.

39.	Patterson, N., et al., *Ancient Admixture in Human History.* Genetics, 2012. **192**(3): p. 1065-1093.

40.	Jeong, C., et al., *The genetic history of admixture across inner Eurasia.* Nature Ecology & Evolution, 2019. **3**(6): p. 966-976.

41.	Lazaridis, I., et al., *Genomic insights into the origin of farming in the ancient Near East.* Nature, 2016. **536**(7617): p. 419-424.

42.	Lazaridis, I., et al., *Ancient human genomes suggest three ancestral populations for present-day Europeans.* Nature, 2014. **513**(7518): p. 409-413.

43.	Biagini, S.A., et al., *People from Ibiza: an unexpected isolate in the Western Mediterranean.* European journal of human genetics : EJHG, 2019. **27**(6): p. 941-951.

44.	Broushaki, F., et al., *Early Neolithic genomes from the eastern Fertile Crescent.* Science, 2016. **353**(6298): p. 499-503.

45.	Lipson, M., et al., *Ancient West African foragers in the context of African population history.* Nature, 2020. **577**(7792): p. 665-670.

46.	Liu, D., et al., *Extensive Ethnolinguistic Diversity in Vietnam Reflects Multiple Sources of Genetic Diversity.* Molecular Biology and Evolution, 2020. **37**(9): p. 2503-2519.

47.	Skoglund, P., et al., *Genomic insights into the peopling of the Southwest Pacific.* Nature, 2016. **538**(7626): p. 510-513.

48.	Prendergast Mary, E., et al., *Ancient DNA reveals a multistep spread of the first herders into sub-Saharan Africa.* Science, 2019. **365**(6448): p. eaaw6275.

49.	Russell, T., F. Silva, and J. Steele, *Modelling the Spread of Farming in the Bantu-Speaking Regions of Africa: An Archaeology-Based Phylogeography.* PLOS ONE, 2014. **9**(1): p. e87854.

50.	Wang, K., et al., *Ancient genomes reveal complex patterns of population movement, interaction, and replacement in sub-Saharan Africa.* Science Advances, 2020. **6**(24): p. eaaz0183.

51.	Brucato, N., et al., *The Comoros Show the Earliest Austronesian Gene Flow into the Swahili Corridor.* American journal of human genetics, 2018. **102**(1): p. 58-68.

52.	Fernandes, V., et al., *Genome-Wide Characterization of Arabian Peninsula Populations: Shedding Light on the History of a Fundamental Bridge between Continents.* Molecular Biology and Evolution, 2019. **36**(3): p. 575-586.

53.	Chintalapati, M., N. Patterson, and P. Moorjani, *The spatiotemporal patterns of major human admixture events during the European Holocene.* eLife, 2022. **11**: p. e77625.

54.	Moorjani, P., et al., *A genetic method for dating ancient genomes provides a direct estimate of human generation interval in the last 45,000 years.* Proceedings of the National Academy of Sciences, 2016. **113**(20): p. 5652.
